# Discovery of reactive peptide inhibitors of human papillomavirus oncoprotein E6

**DOI:** 10.1101/2023.05.25.542341

**Authors:** Xiyun Ye, Peiyuan Zhang, Jason Tao, John C. K. Wang, Amirhossein Mafi, Nathalie M. Grob, Anthony J. Quartararo, Hannah T. Baddock, Ian Foe, Andrei Loas, Dan L. Eaton, Qi Hao, Aaron H. Nile, Bradley L. Pentelute

## Abstract

Human papillomavirus (HPV) infections account for nearly all cervical cancer cases, which is the fourth most common cancer in women worldwide. High-risk variants, including HPV16, drive tumorigenesis in part by promoting the degradation of the tumor suppressor p53. This degradation is mediated by the HPV early protein 6 (E6), which recruits the E3 ubiquitin ligase E6AP and redirects its activity towards ubiquitinating p53. Targeting the protein interaction interface between HPV E6 and E6AP is a promising modality to mitigate HPV-mediated degradation of p53. In this study, we designed a covalent peptide inhibitor, termed reactide, that mimics the E6AP LXXLL binding motif by selectively targeting cysteine 58 in HPV16 E6 with quantitative conversion. This reactide provides a starting point in the development of covalent peptidomimetic inhibitors for intervention against HPV-driven cancers.

## Introduction

High-risk forms of HPV are causative in multiple cancers, including cervical, vaginal, oropharyngeal, and potentially a subset of prostate cancers^1–4^. Despite the introduction of HPV-directed vaccines over ten years ago, the lack of their widespread distribution, uptake by the general population and availability continues to make HPV positive (HPV+) cancers prevalent^4–7^. Among ∼200 identified HPV strains, high-risk HPV16 and HPV18 are responsible for ∼75% of HPV-associated cervical cancers^8–11^. Current treatment strategies of HPV cancers include radiation therapy^4^, surgery^12^, chemotherapy^13^, monoclonal antibody (mAb)^14^ and checkpoint blockade^15–17^. There are no currently approved targeted therapeutics against HPV. The absence of this viral protein in uninfected tissues within the human proteome and its robust association with cancer makes it an attractive target for chemical intervention.

HPV-encoded early protein 6 (E6) and early protein 7 (E7) are primary transforming viral proteins enhancing cancer cell proliferation and contributing to cancer progression^10,18^. HPV E7 primarily binds and inactivates retinoblastoma protein (pRB) and related pocket proteins, p107 and p130, inducing their proteasome-dependent degradation and promoting cell cycle entry^19^. Multiple host proteins interact with E6 via a leucine-rich LXXLL motif, including E6BP^20^, IRF3^21^, paxillin and tuberin^22^; PDZ proteins such as MAGI-1^23^; and other proteins such as p53, E6AP, MAML1, and p300/CBP^19,24^. Nevertheless, the interaction of E6 with the E3 ubiquitin ligase, E6AP, and p53 is thought to be a central transformative pathway of cell immortalization^25–28^. E6 and E7 are active in different cell stages^18^. E7 promotes cell entry to the division phase S^19^, in turn, E6 prevents E7-induced apoptosis by degrading the apoptosis-inducing protein p53^29,30^. HPV+ tumors mostly contain non-mutant p53^31^, as a result, silencing E6 with siRNA can rescue p53 and initiate apoptosis in HPV+ cancer cell lines^32,33^. Therefore, checkpoint networks are primed in HPV+ cells awaiting E6 disruption, providing support for its suitability as an oncology target.

HPV16 E6 (16E6) hijacks E6AP to form a complex with p53 that promotes ubiquitination of p53, leading to its subsequent proteasome-mediated degradation, while neither E6 nor E6AP interact with p53 alone^34–36^. p53 mediates stress response, cell proliferation, and apoptosis, and its downregulation or mutation is a hallmark of carcinogenesis directly affecting efficacy of cancer therapy^37,38^. It was reported that targeting the ubiquitination catalytical domain of ubiquitin ligase E6AP (HECT domain) represses p53 ubiquitination *in vitro*^39^. However, E6AP is a regulator of the proteostasis signaling network^40^ and has wide distribution patterns in humans^41^. Additionally, E6AP has been associated with Angelman syndrome and Prader-Will syndrome^42,43^, diseases causing developmental defects. As such, targeting E6AP could result in on-target toxicity although acute inactivation has not been assessed. Given the potential toxicity associated with targeting E6AP and the lack of 16E6 in healthy human cells, targeting the viral protein 16E6 can provide a wide therapeutic index in HPV+ cancers to limit on-target side-effects.^28^

Efforts in the past decades targeted the 16E6/E6AP protein-protein interaction (PPI). Ribozymes and gene-silencing siRNA that reduce or remove the activity of the HPV E6 oncogene and its protein product have been shown to induce apoptosis in cancer cells^44–47^. Polyhydroxy flavonoids^48–50^ display low micromolar E6 inhibition IC_50_ values and cytotoxicity in HPV+ cancer cells but have not been successful in clinical trials, possibly due to unclear structure-activity relationships (SAR), poor stability, off-target binding, and low specificity^27^. Biomolecules, including E6-binding antibodies and mini-proteins with dissociation constants (*K*_D_) of 10 nM – 60 nM, exhibit nanomolar affinity to E6. However, their inhibitory effect is unsurprisingly hindered by poor penetration through the cell membrane^51–53^.

The X-ray crystal structure of the ternary complex formed between HPV16 E6, an E6AP-derived LXXLL binding motif-containing peptide fused to MBP, and the p53 core domain was reported (PDB: 4XR8, **Figure 1A**)^54^. Investigation of the 16E6/E6AP interface revealed the main recognition domain as a peptide fragment of E6AP (residues 401-418)^55^. The peptide fragment contains a conserved LXXLL motif, critical for both peptide E6AP[401-418]^56^ and E6AP protein binding^55^. The LXXLL motif identified on E6AP formed the basis of our starting point to design peptide inhibitors for 16E6.

**Figure 1.**
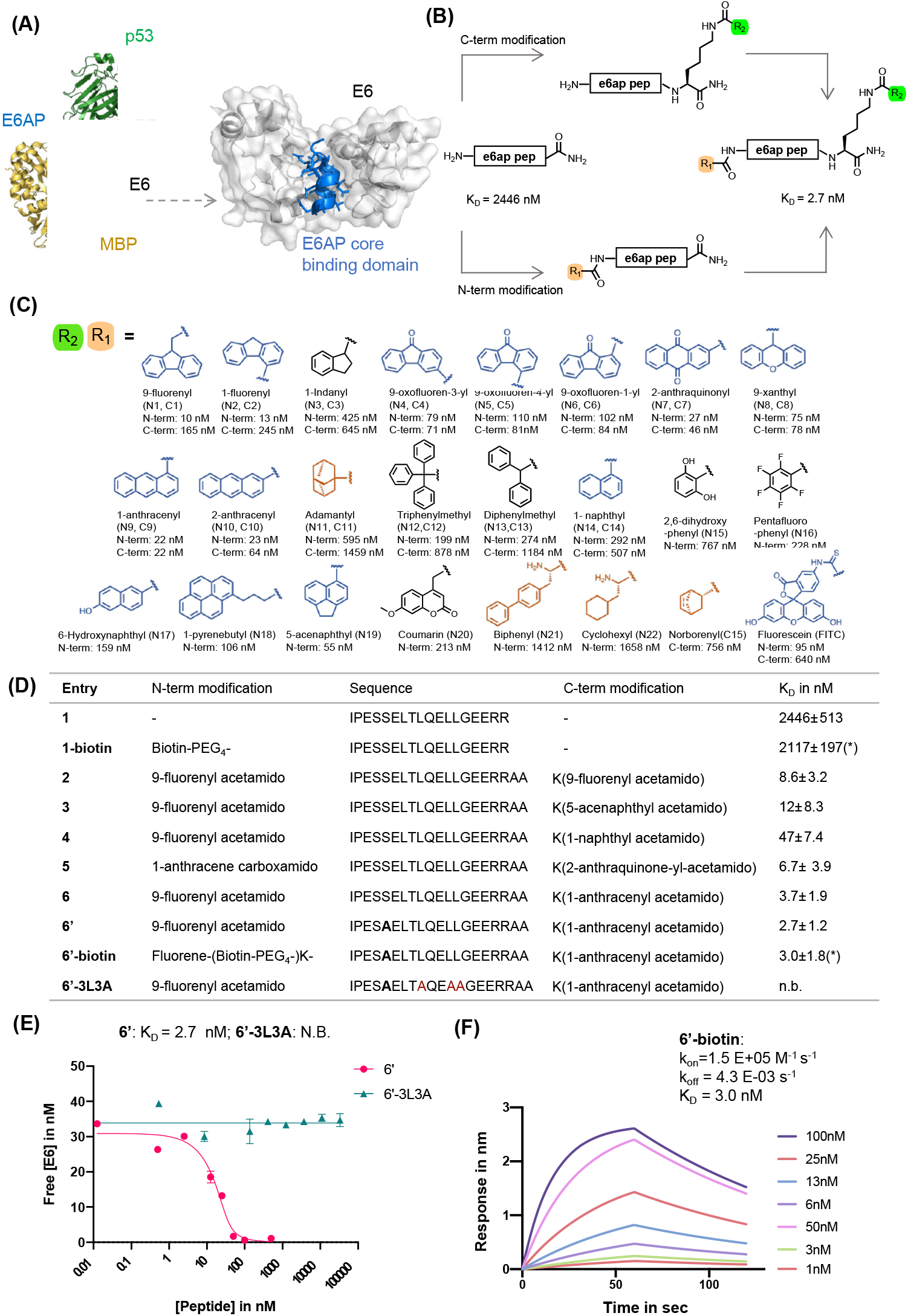
Rational design of E6AP-mimicking peptides. **(A)** Left: X-ray crystal structure of the ternary protein complex formed by 16E6 (white), MBP (yellow) fused to the E6AP LXXLL peptide (blue), and p53 (green). PDB: 4XR8. Right: Interaction interface of 16E6 and the LXXLL motif of E6AP. The E6AP LXXLL peptide binds the 16E6 hydrophobic groove and drives peptide recognition. **(B)** Rational design of a high-affinity E6 binding peptide. The N-terminal and C-terminal lysine side chain primary amines were modified with various small molecules listed in panel C. **(C)** Chemical library of small molecules used for modifying E6AP-based peptides. Planar aromatic residues are in blue, hydrophobic residues in orange, other aromatic groups in black. **(D)** Table of N- and C-terminally modified E6AP peptide mimics. The binding dissociation constant *K*_D_ is measured by bio-layer interferometry (BLI) competition assay, except for \**K*_D_ measured by direct BLI. **(E)** BLI competition assay was used to measure the binding of **6’** and **6’-3L3A**. Streptavidin tips were immobilized with **1-biotin** and dipped into solutions of various concentrations of analyte and 16E6. **(F)** Direct BLI measurement of **6’-biotin**, parameters are estimated to be: k_on_ = 1.5E+05 M^−1^s^−1^, k_off_ = 4.3E-03 s^−1^, and *K*_D_ = 3.0 ± 1.8 nM.

Peptide mimics of E6AP are a promising strategy to overcome the challenges of non-specific binding of small molecules and poor penetration of large biomolecules.^57^ Peptides are of intermediate molecular weight between small molecules and proteins (2-5 kDa) and can be efficient at cell penetration while being endowed with high affinity and specificity to protein targets^58^. A tri-cysteine peptide displayed by phage (*K*_D_ = 118 nM) is prone to aggregation, poorly soluble and unable to compete E6AP-mimicking peptides^59–61^ While an E6AP-mimicking peptide (*K*_D_ = 4 μM)^62^ was reported to target 16E6, it exhibit insufficient affinity toward 16E6, likely due to a rapid *k*_off_ rate^59–61^.

The addition of irreversible crosslinking groups is a suitable strategy for developing inhibitors of ‘undruggable’ proteins. Covalent inhibition, with *k*_off_ values close to zero, is an effective strategy for overcoming resistance in the context of HIV-1 protease inhibitors^63^, where high *k*_off_ values are associated with drug resistance^64,65^. Acrylamide-like warheads are popular in FDA-approved drugs owing to their mild and selective reactivity profile^66–68^. Compared to other highly reactive electrophiles such as chloroacetamides, acrylamide warheads balance stability, reactivity, and substrate selectivity. Warhead-containing peptide inhibitors bind-and-react with targets to establish a covalent bond for irreversible inhibition. For example, acrylamide modified BimBH3 peptide targeting Cys55 on Bcl-2 protein^69^, and acrylamide modified MCL-1 binding peptides crosslink to Cys55^69^. These reactive peptides have multiple advantages compared to traditional peptide inhibitors, typically addressing potency, selectivity, toxicity, dosing, and target scope^66^. Covalent small molecule inhibitors are also exemplified in this regard.

We developed high-affinity 16E6-binding peptides with single digit nanomolar affinity based on the LXXLL binding motif in E6AP, by introducing dehydroalanine (Dha) warheads to irreversibly disrupt the 16E6/E6AP interaction. The N- and C-termini of these peptides were appended to improve 16E6 binding affinity by nearly 1000-fold from 2.4 μM to 2.7 nM. The peptides were designed to crosslink with 16E6 at Cys58 located within the hydrophobic cavity of 16E6 that mediates engagement with the E6AP LXXLL peptide through a Dha warhead. We termed these warhead-containing peptide binders “reactides”. The engineered reactides form covalent peptide-E6 conjugates, irreversibly binding the 16E6 pocket and preventing E6AP association. Based on our findings, we envision reactides as a possible strategy for further development of reactive variants to inhibit the HPV16 E6 protein.

## RESULTS

### Rational design of high-affinity peptide binders to E6

As a starting point for our study, we used the X-ray crystal structure of the 16E6-E6AP-p53 ternary complex (PDB: 4XR8). We selected a 17-mer peptide (**1;** sequence: IPESSELTLQELLGEER, **Figure 1D** and **S1A**) that covers residues 401-417 of E6AP and include the LXXLL motif, as a template for optimization^8^. The binding affinity of peptide **1** was assessed by bio-layer interferometry (BLI) using recombinant MBP-16E6 4C4S protein (MBP-16E6), which contains a maltose-binding protein (MBP) tag and four cysteine to serine mutations to stabilize and solubilize HPV16 E6^70^. The BLI binding assay was run in competition mode by immobilizing biotinylated **1** (**1**-Biotin) onto streptavidin biosensor tips and immersing them across a serial dilution of unlabeled **1** until equilibrium^71^. **1** competed MBP-16E6 binding to **1**-Biotin, revealing a competition *K*_D_ of 2.4 μM (**Figure S1**). The competition *K*_D_ values reported for the peptide variants discussed below were determined by BLI in a similar manner.

While investigating our E6AP peptide designs, we serendipitously observed an increased binding affinity with N-terminal fluorescein isothiocyanate (FITC)-labeled peptide **1** (**N-FITC**) compared to the C-terminal FITC-labeled peptide **1** (**C-FITC**). The resulting **N-FITC** peptide showed a competition *K*_D_ of 95 nM, while **C-FITC** peptide **1** had a competition *K*_D_ of 640 nM. To identify critical residues, we performed alanine (Ala) scanning and synthesized of 17 single Ala mutants of **N-FITC**, peptides **A1** to **A17**, respectively (**Table S1**). The alanine scanning identified hotspot residues Leu9, Leu12, and Leu13 to be critical for binding, as the **A9, A12**, and **A13** peptides did not show measurable binding to MBP-16E6. This observation aligns with L12A and L13A single mutated peptides, which showed decreased E6 binding^72^. Therefore, peptides simultaneously containing the L9A, L12A, and L13A mutations (or **3L3A** triple mutants) were used as negative control peptides in this work (**Figure1D**).

We observe a 900-fold improvement in binding affinity to MBP-16E6 with N-terminal FITC modifications. To investigate the structure-affinity relationships (SAR) contributing to the binding enhancement of terminal FITC modifications, we replaced FITC with a panel of small molecules at the C- or N-terminus of parent peptide **1** (**Figure 1B, C** and **Table S2)**. From 36 synthesized peptides, 8 terminally modified peptides displayed *K*_D_ < 70 nM against MBP-16E6 (Peptides **N1, N2, N7, N9, N10, N19, C4, C9**, and **C10**). At the N-terminus, planar and tricyclic arenes enhance binding by 50 to 250-fold relative to unmodified peptide **1**, while bulky, hydrophobic, or aliphatic small molecules modestly improved affinity by less than 5-fold (**Figure 1C**). The position of fluorene ring substitution has no effect on N- or C-terminal modifications. At the N-terminus, the largest ∼250-fold increase in binding affinity was observed with fluorene modification (**N1**). Removal of a phenyl ring from fluorene (**N1**, *K*_D_ of 10 nM) to indane (**N3**, *K*_D_ of 425 nM) decreases the binding by ∼42-fold. Compared to fluorene-modified **N1**, biphenyl-containing **N21** (*K*_D_ of 1412 nM) decreases the binding by ∼27-fold. These observations suggest that N-terminal planar tricyclic aromatic scaffolds are favorable motifs to enhance binding of parent peptide **1** to 16E6 (**Figure S2A**). The greatest improvements at the C-terminus, from 30 to 111-fold, were found with planar, tricyclic, and polyarenes, while bulky, non-conjugated aromatic or aliphatic small molecules did not significantly improve the binding compared to parent peptide **1** (**Figure 1C** and **Table S3**). The best performing C-terminal modification was anthracene, which showed a competition *K*_D_ value of 22 nM, enhancing binding affinity to MBP-16E6 by ∼111-fold compared to the parent peptide **1** (**Figure S2B**).

Combining the N- and C-terminal modifications with the highest 16E6 binding affinity generated double-modified peptide binders that displayed improvements over single modifications–likely due to synergistic effect from binding features of the two terminal modifications (**Figure 1D**). Peptide **6**, containing N-terminal fluorene and C-terminal anthracene, showed the lowest competition *K*_D_ of 3.7 nM to MBP-16E6 and was selected as our lead for further modifications. Additional optimization was achieved by substituting Ser5 to Ala, as previous Ala scan studies revealed a 2-fold increase in binding affinity (**Table 1**), resulting in peptide **6’** with a competition *K*_D_ of 2.7 nM. To investigate the binding affinity and selectivity of **6’**, biotinylated peptide **6’-biotin** and its 3L3A derivative **6’-3L3A** were synthesized. **6’-3L3A** displayed no observable binding towards MBP-16E6 (**Figure 1E**). **6’-biotin** showed a direct binding *K*_D_ of 3.0 ± 1.8 nM to MBP-16E6(**Figure 1F**). These results demonstrate that **6’** binding to MBP-16E6 is sequence-specific and requires the tri-leucine hotspot motif LXXLL. To assess target selectivity, **6’-biotin** was immobilized and its affinity towards the unrelated proteins murine double minute 2 (MDM2) and an LXXLL motif-binding protein thyroid hormone receptor alpha (THRA)^73^ were measured by BLI. No appreciable signal during the association or dissociation steps was observed by BLI (**Figures S3B and S3C**), indicating that peptide **6’** selectively binds MBP-16E6.

### Development of a dehydroalanine-modified E6 binding peptide that crosslinks to MBP-16E6

Irreversible covalent inhibition is an effective strategy to increase drug potency and selectivity toward inhibiting ‘undruggable’ targets^74^. The binding interface of 16E6 contains the nucleophilic Cys58 residue in proximity to the E6AP LXXLL peptide binding pocket (**Figure 2A**. To exploit this finding, we synthesized reactides **E1–E11** based on **E0**, a truncated version of peptide **1**, where a Cys-reactive acrylamide replaced Gly9, Glu10, or Glu10 of **E0** (**Figure 2B** and **Figure S4A**). We tested multiple electrophiles including phenylacrylamide (Phacr), acrylamide (Acr), propiolamide (Ppa), and Dha (**Figure S4B**)^75^. Liquid chromatography-mass spectrometry (LC-MS) monitored conjugation between MBP-16E6 protein and the reactides, and the area under the total ion peak was used to estimate crosslink yield. Gly9 was found to be the closest residue to Cys58 and its substitution with Dha gave the highest crosslink yield of 74%, and it was thus chosen as our lead reactide scaffold **E3 (Figure 2C)**.

**Figure 2.**
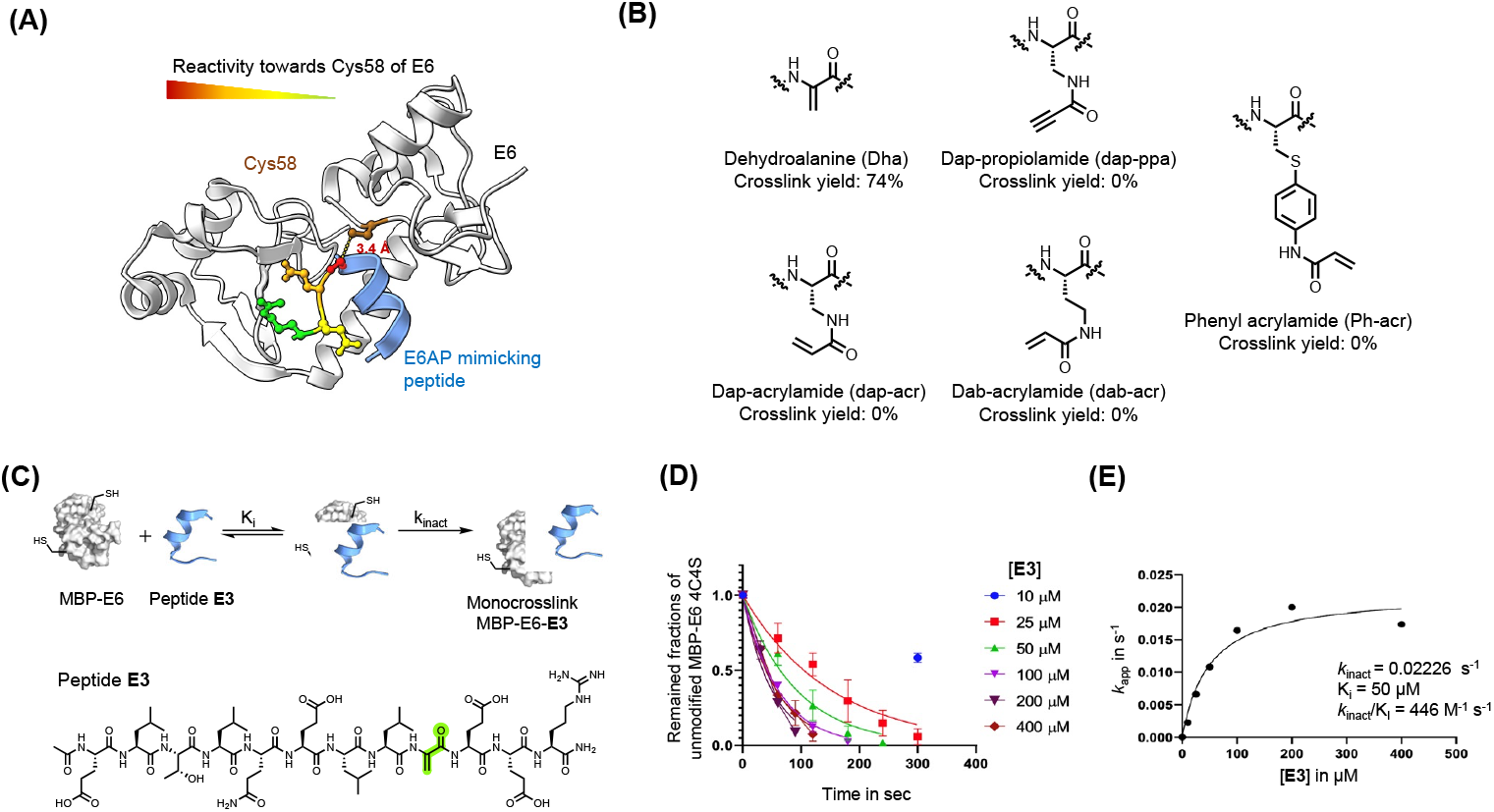
Dha was installed on reactive peptides. **(A)** Analysis of the main binding interface of 16E6 and E6AP. Cys58 of 16E6 is highlighted in brown. Residues of E6AP were colored based on their measured reactivity towards Cys58 of 16E6: Gly9 in red, Glu10 in orange, Glu11 in yellow, and Arg12 in Green. **(B)** Structures of electrophiles. **(C)** A scheme of the bind-and-react strategy. Structure of peptide **E3** containing a Dha electrophile highlighted in green. **(D)** Apparent kinetic constant k_app_ was calculated from the time-course of a protein depletion assay, where 50 nM of MBP-16E6 was incubated with different concentrations of **E3. (E)** A series of *K*_app_ values were plotted against the corresponding **E3** concentration to estimate k_inact_ and *K*_i_.

Kinetic studies were performed with **E3** to estimate the binding constant *K*_i_ and first-order rate constant (*k*_inact_) involved in the two-stage bind-and-react strategy. Crosslink yield was monitored over time by LC-MS (**Figure 2D** and **2E**). *K*inetic parameters were estimated assuming a steady-state approximation, and a *k*_*inact*_ value of 0.022 s^-1^ and *K*_i_ value of 50 μM were obtained. The *k*_inact_/*K*_i_ ratio was calculated to be 446 M^-1^ s^-1^, indicating that improvements in binding affinity will produce a more efficient covalent inhibitor^66^. For comparison, the FDA-approved small molecular drug nirmatrelvir showed a *k*_inact_/*K*_i_ value of 55 mM^-1^ s^-1^.^76^

First we installed Dha onto peptide **6’** to obtain reactide **7**, which has a net charge of −3 that may be detrimental to cell permeability^77,78^. To increase positive charge, improve solubility, retain binding affinity to 16E6, and further reduce molecular weight, we performed residue substitutions and truncations on **7**. The LXXLL proximal RRN*KK* [417-421] segment from the E6AP protein was appended to the C-terminus of reactide **7** to increase the total charge from −3 to −1 (reactide **8**). Next, Glu3Gln, Glu15Gln, and Glu16Ala mutations were applied to increase the total charge to +2, resulting in peptides **9** and **10**. To reduce molecular weight, Ala13, Gln14, and Asn19 residues were omitted from peptide **10** to generate **11, 12**, and **13**. Reactides were evaluated by the BLI competition assay against **1-biotin** to estimate their inhibitory constant, which we report here as an apparent *K*_i_ value. Peptides **8**–**13** showed comparable binding affinity to the parent reactide **7**, with an apparent *K*_i_ ranging from 11 nM to 39 nM (**Figure S5**). Reactide **13** was selected for further characterization since it has the smallest molecular weight, highest net charge, and relatively low apparent *K*_i_ = 17 ± 3.9 nM (**Figure 3A, 3B** and **3C**). The negative control derivative of **13**, (**13-3L3A**) showed no observable binding to MBP-16E6. A kinetic study was performed with **13** and revealed a *k*_*inact*_ of 0.027 s^-1^, a *K*_i_ value of 120 nM and a *k*_inact_/*K*_i_ ratio of 270 mM^-1^ s^-1^ (**Figure 3D and 3E**). Reactide **13** represents a 504-fold improvement over **E3**.

**Figure 3.**
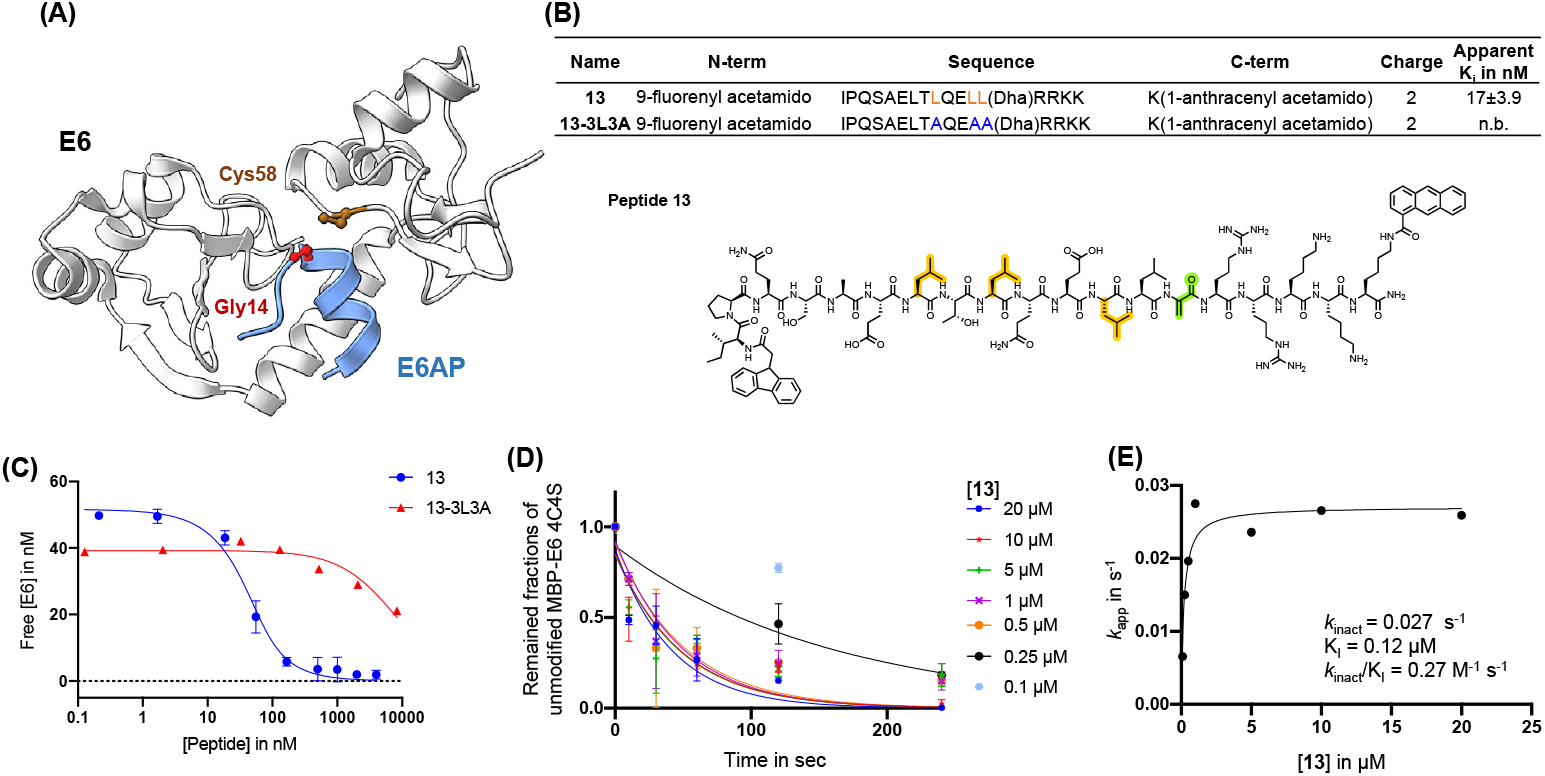
Affinity matured 16E6-binding peptide is endowed with improved reactivity. **(A)** Main binding interface of 16E6 and E6AP. Cys58 of 16E6 is highlighted in brown and targeted by an electrophile substituted at Gly14. **(B)** Sequence table of E6AP-mimicking **13** and **13-3L3A** (control peptide). Apparent *K*_i_ is determined by BLI (n.b., non-binding). Peptides were in competition with immobilized **1-Biotin** following a 30 min incubation with 16E6. Dha: dehydroalanine. Structure of **13**, Dha (green), tri-leucine (orange). **(C)** BLI competition assay measurement of **13** and **13-3L3A** estimated apparent *K*_i_ in nM. **(D)** Apparent kinetic constant k_app_ was calculated from the time-course of protein depletion assay, where 10 nM of MBP-16E6 was incubated with different concentrations of **13. (E)** *K*_app_ was plotted against the corresponding **13** concentration to estimate k_inact_ and *K*_i_.

### Reactide 13 selectively crosslinks to MBP-16E6 in PBS

**13** uses a two-stage bind-and-react strategy to covalently crosslink to MBP-16E6 and block the E6AP binding site. When **13** (3 μM) and MBP-16E6 (1 μM) were incubated in PBS at 37 °C for 2 h, protein deconvolution mass spectra revealed >99% mono-crosslinking of **13** to MBP-16E6 (**Figure 4A**). This was achieved despite the presence of 10 cysteine residues on MBP-16E6. To confirm site-selectivity, **13** was incubated with the MBP-16E6 mutant C58S for 12 h in PBS at 37 °C. No crosslinking was observed between **13** and MBP-16E6 C58S (**Figure 4B**), which supports Cys58 as the sole site of MBP-16E6 modification. **13** showed no crosslink to THRA (**Figure S6)** Additionally, the negative control **13-3L3A** showed no observable crosslinking to MBP-16E6 protein (**Figure 4C**), supporting the selectivity of **13**. Therefore, **13-3L3A** was used as a negative control for subsequent peptides.

**Figure 4.**
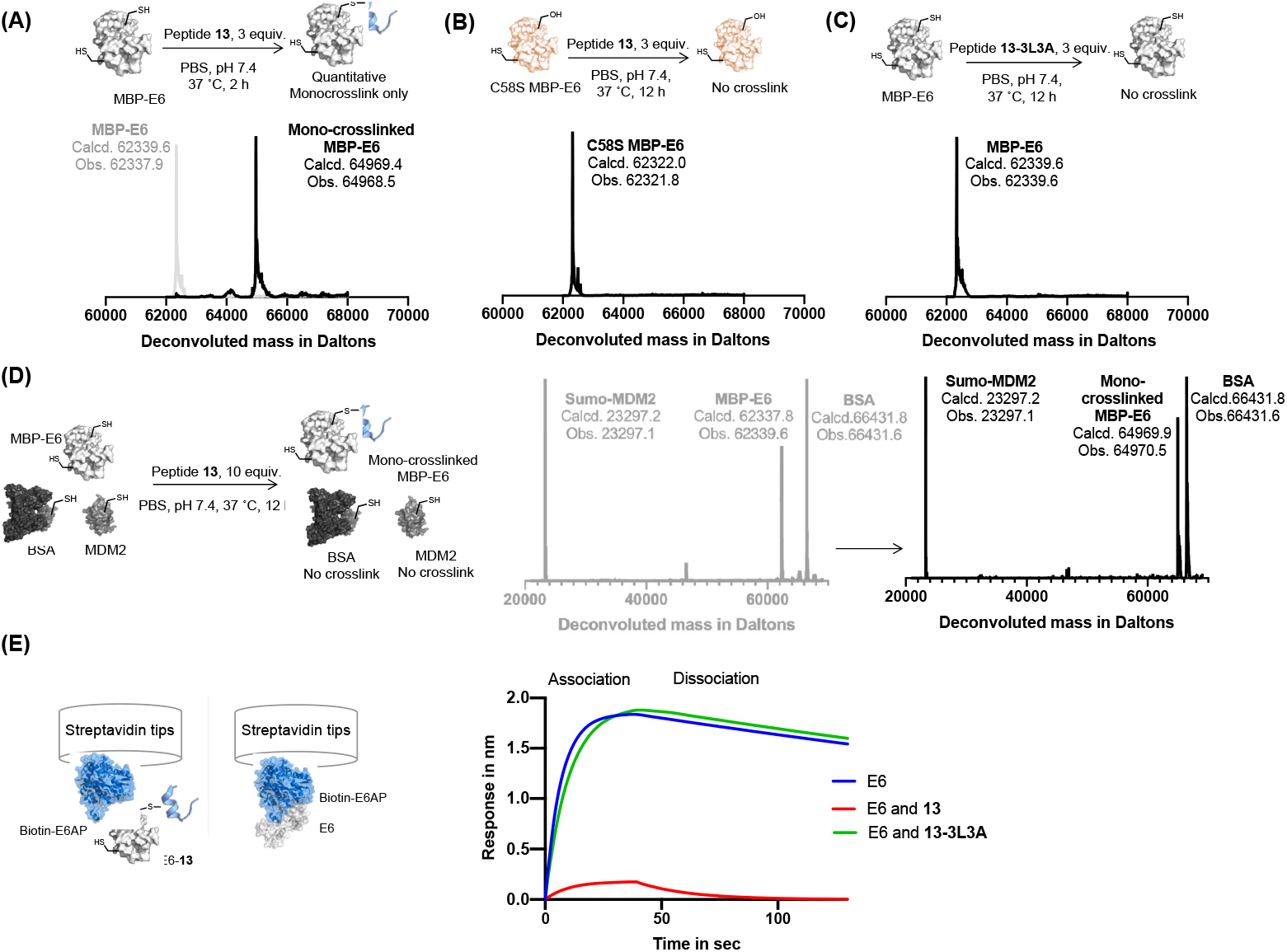
Reactive peptides selectively crosslink to 16E6. **(A)** Crosslinking reaction of MBP-16E6 (1 μM) and reactide **13** (3 μM). Quantitative mono-crosslinked MBP-16E6-**13** was observed at 2 h. **(B)** Crosslinking of MBP-16E6 C58S (1 μM) and peptide **13** (3 μM), no appreciable amount of reaction product was observed after 12 h. **(C)** Crosslinking of MBP-16E6 (1 μM) and peptide **13-3L3A** (3 μM), no appreciable amount of reaction product was observed after 12 h. **(D)** Protein-selective intermolecular crosslinking of MBP-16E6 in a protein mixture. Only 16E6 was modified, as indicated by the LC-MS analysis. **(E)** BLI assay determined 16E6/E6AP complex formation. 1 μM of Biotin-E6AP protein was immobilized onto streptavidin tip and dipped into 1 μM of MBP-16E6 only (blue), MBP-16E6 mixed with 1 μM of **13** (red) or 1 μM of **13-3L3A** (green).

### Reactide 13 selectively crosslinks to MBP-16E6 in a protein mixture

To investigate whether **13** selectively crosslink to 16E6 in a protein mixture we monitored modification of Sumo-MDM2^25-109^ which contains one cysteine and bovine serum albumin (BSA) which has 35 cysteine residues. A mixture of the three proteins, 16E6 (1 μM), BSA (1 μM), and Sumo-MDM2^225-109^ (1 μM) was incubated with **13** (10 μM) at 37 °C for 12 h and resolved by LC-MS (**Figure 4D**). The results indicate 16E6 is mono-crosslinked with >95% conversion with no observable modifications to BSA or MDM2. These results demonstrate crosslinking of **13** is specific to MBP-16E6 and depends on the hotspot motif LXXLL.

### Disruption of 16E6/E6AP interaction by reactide 13

To assess the disruption of 16E6/E6AP interaction by reactide **13**, we set up an 16E6/E6AP BLI binding assay by immobilizing biotinylated E6AP protein onto streptavidin tips. The 16E6/E6AP interaction was evaluated as the response (in nm) during the association step when dipped into recombinant MBP-16E6 solution (1 μM). MBP-16E6 conjugated with reactide **13** had a significantly decreased BLI response signal to the immobilized E6AP protein, while the control reactide **13-3L3A** had no impact (**Figure 4E**). The decrease in BLI response signal indicates that the 16E6-**13** conjugate loses its E6AP-binding ability, as its E6AP binding pocket was occupied by **13**.

### Molecular modeling of E6-Peptide-13 complex

Molecular modeling was conducted to understand the mechanism by which reactide **13** disrupts the 16E6/E6AP interaction using molecular docking and molecular dynamics (MD) simulations. The MBP tag may affect the conformation of the isolated E6AP-based peptide from the ternary complex structure with 16E6 and p53 (PDB: 4XR8).^8^ Therefore, we removed the MBP tag and p53 protein from the structure and ran a 1.1 μs simulation of the 16E6-bound E6AP-LXXLL peptide. The peptide remained stable with an r.m.s.d. of 0.4, indicating **α**-helical features (**Figure S7**). This enabled us to build the model of **13** by incorporating the flexible N- and C-terminal sequences into the alpha helix of the E6AP LXXLL peptide and perform extensive conformational sampling calculations to optimize the reactide **13** model (**Figure S8B**). The covalent bond formed between **13** and 16E6 Cys58 was not considered when initiating the molecular docking to avoid any biases in the calculations. The docking result suggested that the primary driver of **13** binding to 16E6 are electrostatic and van der Waals interactions (**Figure 5A**). By refining the top candidates obtained from the docking calculations using MD simulations (**Figure S8C**), we observed that the Dha (C atoms) resides within 3.7 Å from Cys58 (S atoms), confirming optimal placement for **13** to cross-link with 16E6. The thioester bond between 16E6 and Dha acts as a covalent lock, securing **13** at the 16E6/E6AP interface. Computational modeling confirms the importance of this bond as shown in **Figure S8D** and **Figure S9**. Our trajectory r.m.s.d analysis (**Figure 5B**) for the 16E6 and **13** complex and **13** alone indicates that the covalent bond between Cys58 and Dha is instrumental in stabilizing **13** and effectively restricts the flexibility of the E6AP-LXXLL core sequence with an r.m.s.d of 1.5 ± 0.3 Å. Taken together, our molecular simulations reveal that reactide **13** occupies the same position as the parent E6AP LXXLL peptide (r.m.s.d = 1.9 Å, measured for Cα atoms), leading to the blocking of E6AP/16E6 binding (**Figure 5C**).

**Figure 5.**
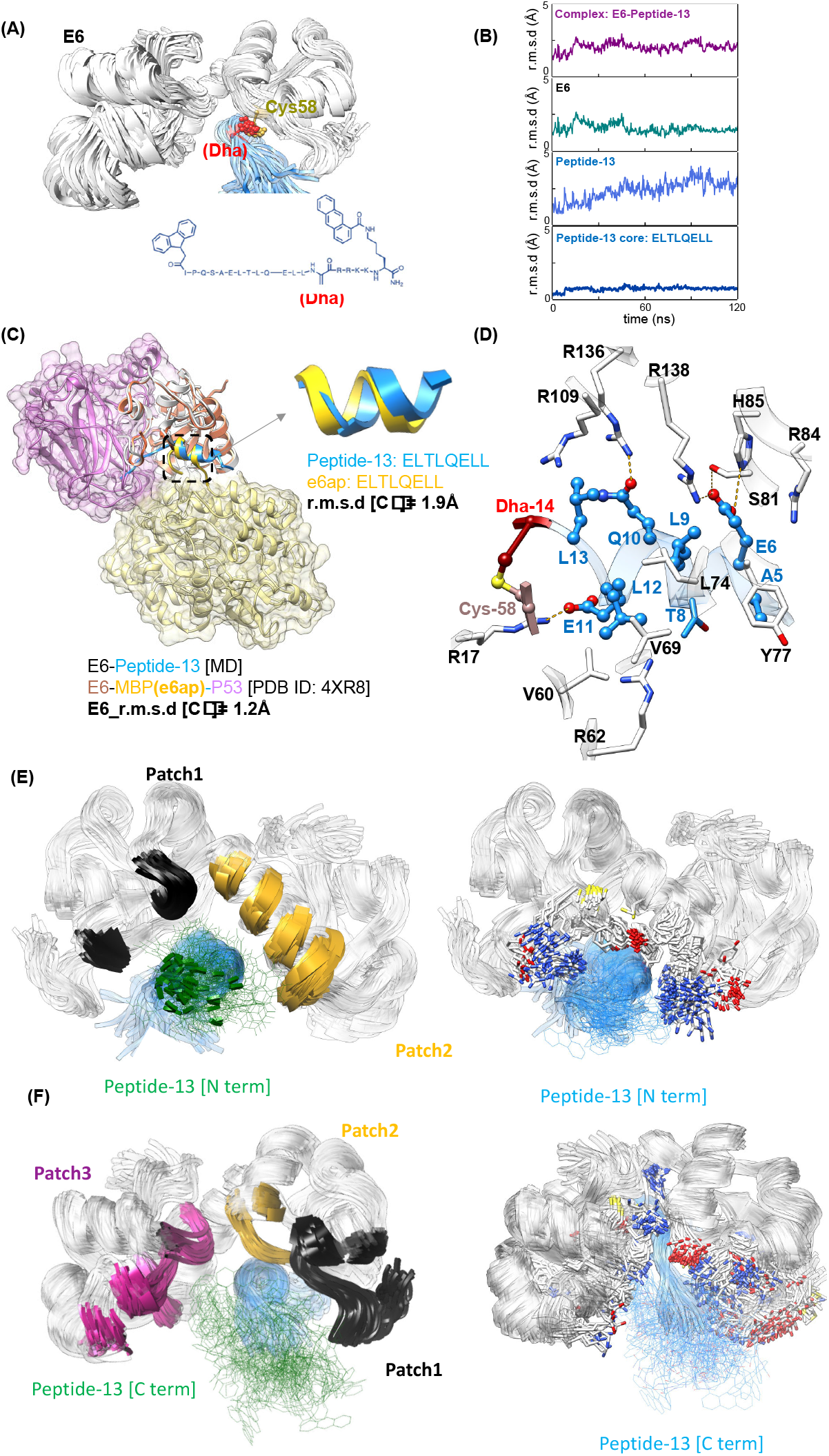
Molecular modeling of the E6-13 conjugate. **(A)** Structural representation of 16E6 (white) bound to **13** (blue) featuring the Dha warhead (red), obtained through ∼120 ns MD simulation. Structure represents the averaged configurations of **13** within calculated clusters (see methods). **(B)** Trajectory analysis of root mean squared differences (r.m.s.d) during MD simulatioms, showing stability and modest conformational changes of the complex. **(C)** Comparison of molecular modeling of the **13**:16E6 complex with the X-ray structure of 16E6 (coral)-E6AP LXXLL peptide (yellow), p53 (plum) complex (PDB ID: 4XR8). All r.m.s.d. were calculated for the Cα atoms. All strucutures were compared to the initial struture. **(D)** Intermolecular interactions between 16E6 (white) and **13** (blue), identified and analyzed from the MD simulation. Interactions shown in (D) are persistent during the ∼120 ns MD simulation. Dotted yellow lines represent hydrogen bonding. **(E)** Covered surfaces by the N-terminus within 5 Å: Patch1 in black [V38, Y39, C40, K41, R62, E63], Patch2 in yellow [C73, F76, Y77, I80, Y83, R84, H85, R136]. Left: E6 and **13** N-term detailed interactions. **(F)** Covered surfaces by the C-terminus within 5 Å: Patch1 [M8, F9, Q10, D11, P12, Q13, E14, R15, P16, R17, K18, L19, P20, Q21, D25]. Patch2 in yellow [A53, D56, R55, L57, C58*]. Patch3 in purple [Y99, K101, D105, L107, I108, R109, C110, C113, Q114, K115, P116, L117, R136, R138, W139, T140]. Left: E6 and C-terminal **13** detailed interactions.

The modeling suggests important side chain interactions between **13** and 16E6 basic residues, including E6AP Glu6/16E6 Arg138 and E6AP Glu11/16E6 Arg11. 16E6 Arg138 is crucial as Arg138 mutation reduces E6AP recruitment and interactions substantially^79^. In addition, 16E6 Arg17 forms a salt bridge with the C-terminus of the E6AP core peptide. The Glu6 carboxylate group is essential for **13** binding and establishes an additional network of hydrogen bonding with Ser81 and His85 from 16E6 (**Figure 5D**). 16E6 Arg136 forms a hydrogen bond with Gln10 in the E6AP core. The keystone 16E6 Arg109 interacts with Leu13 through hydrophobic interactions, consistent with crystallographic observations^54^. Notably, the alanine substitutions of both Arg136^54^ and Arg109^79^ led to impaired peptide binding, highlighting their crucial roles in stabilizing the 16E6/E6AP complex, though R136 exhibited conformational disorder^54^ and was oriented away from the peptide towards MBP. Collectively, peptide **13** targets all residues appearing in the binding pocket of E6 to disrupt the binding interface of 16E6 and E6AP.

The N- and C-terminal groups are located on flexible loops and dynamically bind to 16E6, making it challenging to identify static interactions similar to those found for the core motif. However, despite their dynamic binding, the N- and C-terminal groups still interact with important surfaces of E6. This flexibility and ability to cover essential surfaces of 16E6 may prevent 16E6 from recruiting E6AP (**Figure 5E and 5F**). Notably, the binding interface of the N and C-terminal groups includes several key residues that are essential in the 16E6-E6AP complex. Therefore, these results suggest that hydrophobic modifications contribute to the inhibition of 16E6.

## Discussion

The affinity of a 16E6-binding peptide was improved from ∼2.4 μM to ∼2.7 nM, a nearly 900-fold improvement, by modifying its termini with pharmacophores. We and others have observed binding improvement by terminal fluorescein labeling. For example, Frank et al. reported an unexpected binding improvement with a fluorescein-labeled peptide inhibitor of replication protein A^80^. Similarly, Torner et al. reported that tryptophan addition to the C-terminus of MDM2-binding peptide PMI has increased binding by 50-fold and is likely to engage a secondary binding pocket^81^. Likewise, the terminal small molecule modifications on our E6AP-mimicking peptides have improved binding affinity, with retention of sequence specificity and selectivity for the E6AP/16E6 binding groove. Upon peptide anchoring to the binding groove through LXXLL residues, pharmacophores bearing arenes may adhere to adjacent surface patches to enhance binding. Although the detailed mechanism of interaction of the added aromatic moieties in the 16E6 binding groove remains to be determined by further structural studies, the observations reported in this work should be of interest as insights gained toward strategies of improving peptide binding affinity.

Despite increasing numbers of reports describing peptide-based covalent inhibitors, the strategy is under-exploited to target challenging protein-protein interactions^82^. One major drawback for this strategy is off-target crosslinking^66,68^ which can be mitigated through a bind-and-react strategy. With this approach, the risks of non-specific crosslinking and non-selective inhibition are reduced through specific binding of the base peptide sequence which brings the reactive warhead and its target residue in close proximity. This may allow for the use of less reactive groups to alleviate non-selective modification.

Dha is the simplest dehydroamino acid and is found in some microbial peptides. Due to its electrophilic nature and lack of geometric isomers resulting from the methylidene group, it has been utilized as a reactive probe targeting enzymatic mechanisms.^83–87^ To our knowledge, we exploit for the first-time Dha as an electrophilic warhead to generate a reactide for targeting a cancer-relevant PPI. Reactivity of the reactide was fine-tuned by adjusting the electrophile and distance between the warhead and peptide backbone. Optimal crosslinking efficiency was achieved with Dha placed directly onto the peptide backbone to fit in the tight binding pocket at the target protein surface containing Cys58, as suggested by the co-crystal structure. The reactide is stable and not reactive to 0.4 mM free Cys and GST in cell media (**Figure S11**).

Although it is desirable to inactivate all pathogenic HPV subtypes with Cys58-targeted reactides only a subset contain a cysteine within the LXXLL binding motif (**Figure S12**). Among high-risk types, only HPV35, HPV45 and HPV16 contain Cys58. Further research into effective HPV targeting sites should expand the window of opportunity for HPV+ diseases.

## Conclusion

In this study, we investigated the inhibition of HPV16 E6 by utilizing peptides that mimic the native E6AP binding scaffold. Our findings from alanine scanning mutagenesis indicate that binding depends on the LXXLL sequence motif. Further affinity maturation with terminal small molecular pharmacophores endowed peptides with enhanced binding affinity towards 16E6 while maintaining stability and selectivity. Using these matured peptides, we designed high-affinity, selective, and potent peptide-based covalent inhibitors targeting the 16E6 oncoprotein using a cysteine-reactive acrylamide warhead. We demonstrate reactide **13** as an efficient and selective crosslinking tool and report the first covalent peptide for irreversible inhibition of HPV16 E6. As a result of the high potency of peptide **13**, we anticipate that a bind-and-react strategy can be applied more broadly in the development of irreversible covalent peptide inhibitors. Continued advancement in covalent chemistry compatible with solid-phase peptide synthesis and biological display methods are expected to expand substrate scope and optimize crosslinking efficiency. This opens opportunities for the design of potent and selective inhibitors across a wide range of biological targets including challenging protein-protein interactions.

Challenges remain for the development of reactides with covalent warheads, including peptide stability in a biological context, bioavailability, renal clearance, and biological barrier penetration properties. Possible solutions include the incorporation of unnatural amino acids, albumin-binding modifications, and cyclization methods^57^. Further investigation is warranted into the cellular response, inhibitory mechanism, and cell penetration properties of this new class of 16E6 reactides. Nevertheless, our studies establish a foundation for the next generation of HPV-targeted therapeutics and put forward design principles to utilize in bind-and-react strategies for other high-risk HPV proteins and a broad variety of other oncogenic proteins.

## Acknowledgment

Calico Life Sciences (to B. L. P.) provided financial support for this work.

## Author contributions

B. L. P., A. H. N., Q. H. and X. Y. conceived the project; X. Y., P. Z., J. T., A.M. and J. C. K. W. performed experiments with input from H. T. B., I. F., D. L. E., Q. H., A. H. N., A. J. Q., A. L. and N. M. G. X. Y. wrote the manuscript with input from all authors.

