## Supporting Information for "Discovery of reactive peptide inhibitors of human papillomavirus oncoprotein E6"

Contents

1. Supplementary figures and tables ... 3
1.1 Materials ... 18
1.2 Fast flow synthesis of peptides ... 18
1.3 Automated flow peptide synthesis (AFPS) set-up ... 18
1.4 Method for peptide acetylation ... 19
1.5 Method for Alloc deprotection ... 19
1.6 Method for biotin labeling ... 19
1.7 Method for TAMRA labeling ... 19
1.8 Method for FITC labeling ... 19
1.9 Cleavage of peptides ... 19
1.10 Pd Mediated Conjugation ... 20
1.11 Dehydroalanine formation ... 20
1.12 General procedure for Cross-linking MBP-16E6 with Covalent Peptides ... 20
1.13 Purification of the crude peptide. ... 20
1.14 Method for LC-MS characterization. ... 20
1.15 Direct binding measurement by BLI ... 21
1.16 Competitive binding assay by BLI ... 21
1.17 protein expression and purification method ... 22
1.17.1 E6AP ... 22
1.17.2 MBP-16E6 ... 23
1.17.3 SUMO-MDM2 ... 24
1.18 Complex disruption assay ... 25
1.19 Structural modeling of 16E6-bound Peptide-13 ... 25
2. LC-MS characterization of peptides ... 27
2.1 LC-MS methods ... 27
2.2 Peptides 1-13 ... 27
2.3 Peptide N1-N21 and N-FITC ... 42
2.4 Peptide C1-C13 and C-FITC ... 54
2.5 Peptide A1-A17 ... 65
2.6 Peptide E1-E11 ... 74
3. Binding characterization ... 82
4. Reference ... 93

### 1. Supplementary figures and tables

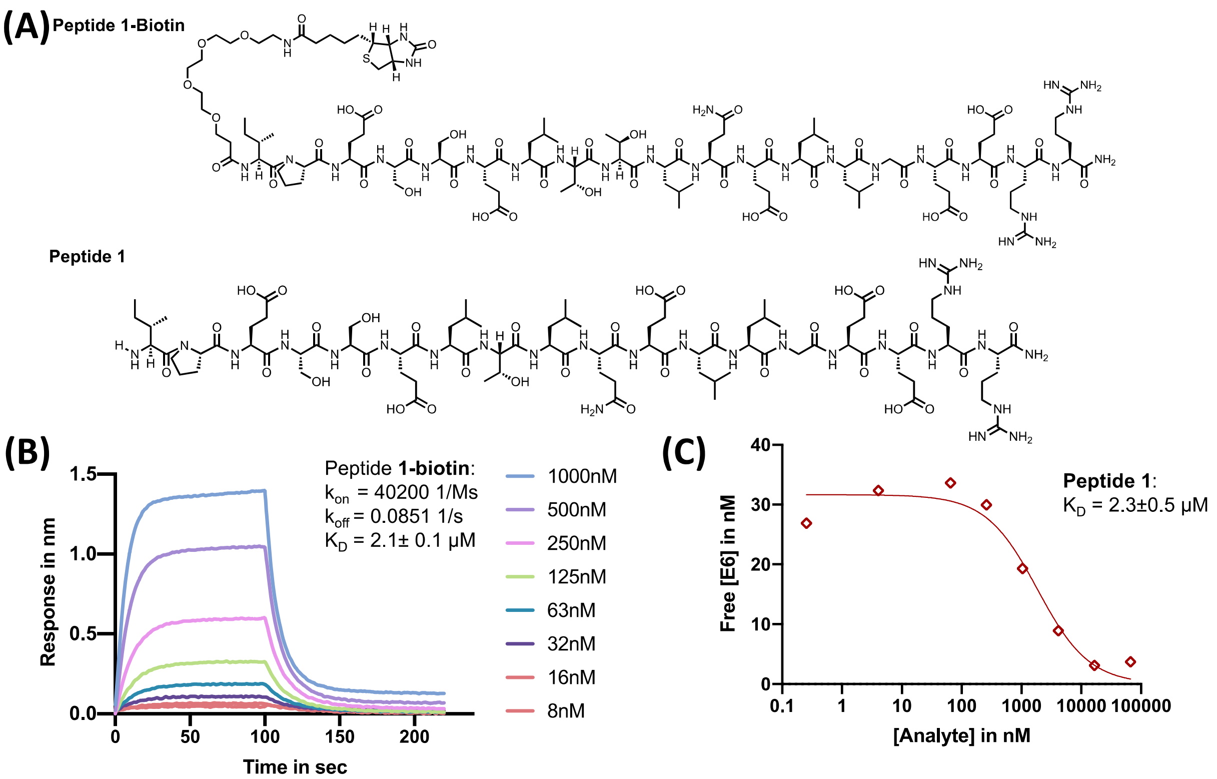


**Figure S1.** Determination of binding constant between peptide **1** and 16E6 by direct binding assay in BLI and competition BLI binding assay. (**A**) Structure of peptide **1** and peptide **1-biotin**. (**B**) BLI measurement of **1-biotin** and 16E6, with estimated parameters: $k_{on}$ = 40200 $M^{-1}s^{-1}$, $k_{off}$ = 0.0851 $s^{-1}$and $K_D$ = 2.1± 0.1 μM (N=3). MBP-16E6 protein concentration used in each channel is indicated on the right. $k_{on}$ is the on rate constant, $k_{off}$ is the off rate constant, $K_D$ is the kinetic apparent dissociation constant. **(C)** Peptide **1** binding was assessed by competition binding assay. Various concentrations of **1** were mixed with 30 nM MBP-16E6 protein. Peptide **1-biotin** was immobilized onto the streptavidin sensor tips to compete for MBP-16E6 protein with various concentrations of unlabeled peptide **1** in the solution (33333 nM, 8333 nM, 2083 nM, 520 nM, 130 nM, 32 nM, 2.0 nM or 0.1 nM). Dissociation binding constant estimated as $K_D$ = 2.3 ± 0.5 μM. Error is the quadratic curve fitting standard error of the mean (SEM) reported by the Prism 8 software (N=3).

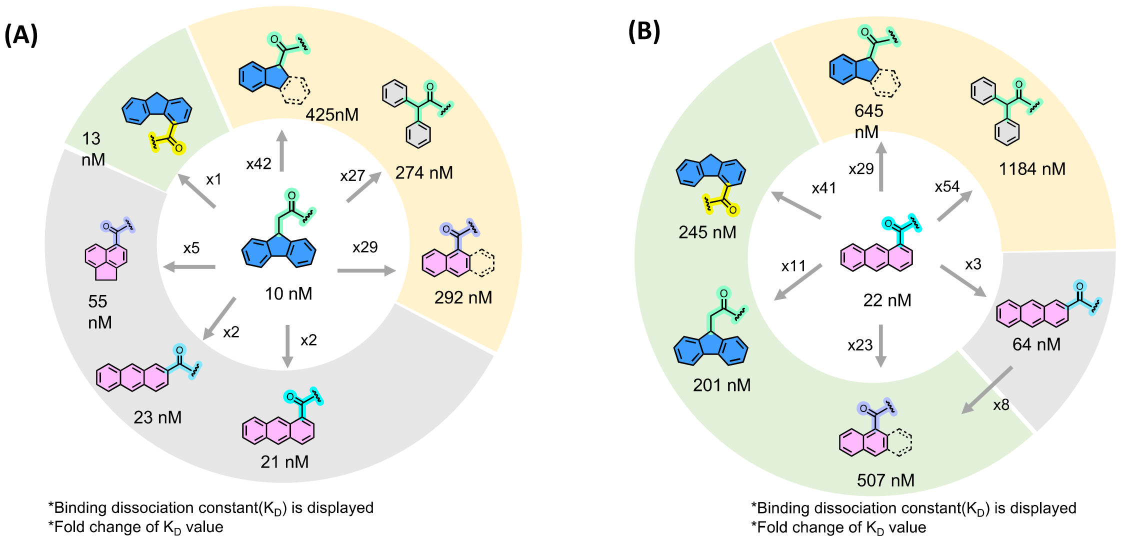


**Figure S2.** Structure-activity-relationship (SAR) study of small molecule modifications. (**A**) N-terminal modifications SAR. Green: modification is not sensitive to ring substitution position. Grey: tricyclic, planar, and aromatic molecules improve the affinity to the low nano-molar range. Yellow: Removing a phenyl ring from the structure or breaking the strain between two phenyl moieties will decrease the binding by 30-50-fold. (**B**) C-terminal modifications SAR. Green: binding improvement of anthracene modification is not sensitive to ring substitution position. Green: two phenyl rings are required for a mid-micromolar binding. Yellow: Removing a phenyl ring from the structure, breaking the strain between two phenyl moieties, will decrease the binding by 30-50-fold. $K_D$ values are measured by BLI binding assay in a competition mode.

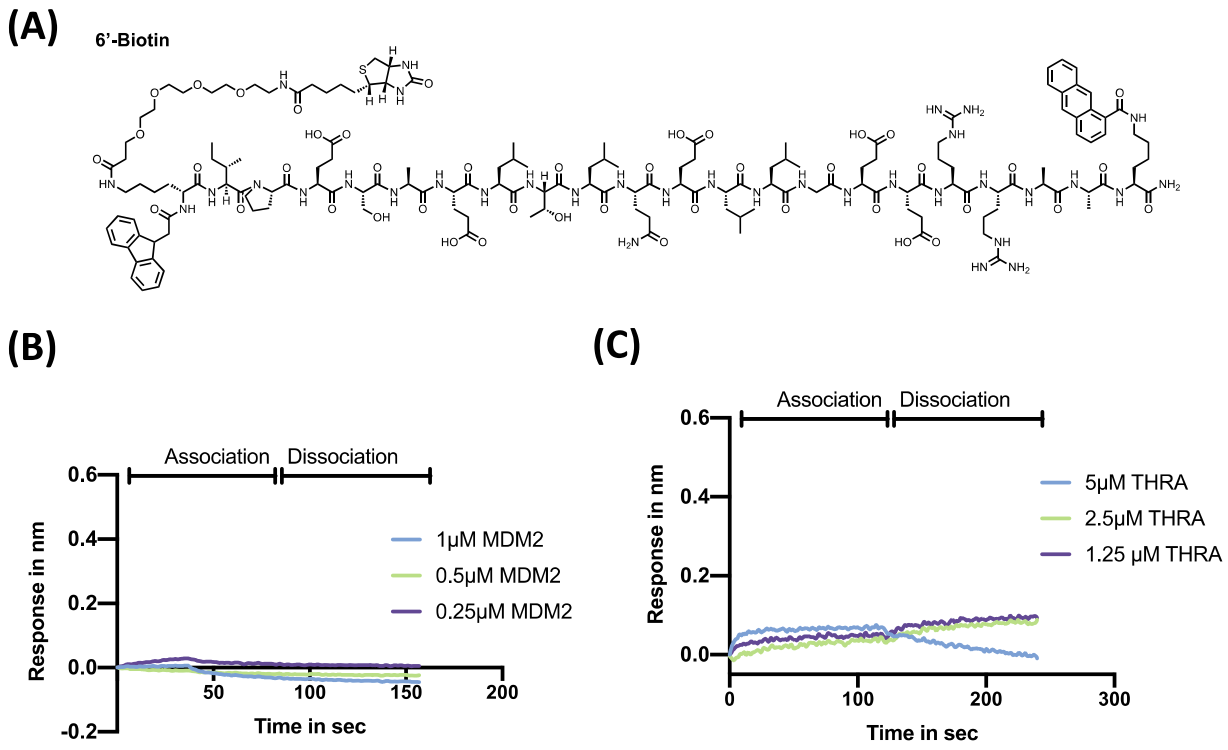


**Figure S3.** (**A**) Structure of **6'-biotin**. (**B**) Direct BLI binding assay of **6'-biotin** against SUMO-MDM2. No observable association and dissociation were observed (N=2). (**C**) Direct BLI binding assay of **6'-biotin** against THRA. No observable association and dissociation were observed (N = 2).

| Entry | Name | N term | Spacer | 1 | 2 | 3 | 4 | 5 | 6 | 7 | 8 | 9 | 10 | 11 | 12 | 13 | 14 | 15 | 16 | 17 | Binding affinity in nM | Ratio to WT |
|---|---|---|---|---|---|---|---|---|---|---|---|---|---|---|---|---|---|---|---|---|---|---|
| 1 | IPES | - | bAla | I | P | E | S | S | E | L | T | L | Q | E | L | L | G | E | E | R | 2589±1394 | - |
| 2 | N-FITC | FITC | bAla | I | P | E | S | S | E | L | T | L | Q | E | L | L | G | E | E | R | 186±32 | 1 |
| 3 | A1 | FITC | bAla | **A** | P | E | S | S | E | L | T | L | Q | E | L | L | G | E | E | R | 944±182 | 5 |
| 4 | A2 | FITC | bAla | I | **A** | E | S | S | E | L | T | L | Q | E | L | L | G | E | E | R | 691±157 | 4 |
| 5 | A3 | FITC | bAla | I | P | **A** | S | S | E | L | T | L | Q | E | L | L | G | E | E | R | 638±54 | 3 |
| 6 | A4 | FITC | bAla | I | P | E | **A** | S | E | L | T | L | Q | E | L | L | G | E | E | R | 739±113 | 4 |
| 7 | A5 | FITC | bAla | I | P | E | S | **A** | E | L | T | L | Q | E | L | L | G | E | E | R | 125±17 | 0.6 |
| 8 | A6 | FITC | bAla | I | P | E | S | S | **A** | L | T | L | Q | E | L | L | G | E | E | R | 1706±209 | 9 |
| 9 | A7 | FITC | bAla | I | P | E | S | S | E | **A** | T | L | Q | E | L | L | G | E | E | R | 1114±152 | 6 |
| 10 | A8 | FITC | bAla | I | P | E | S | S | E | L | **A** | L | Q | E | L | L | G | E | E | R | 3748±680 | 20 |
| 11 | A9 | FITC | bAla | I | P | E | S | S | E | L | T | **A** | Q | E | L | L | G | E | E | R | >10000 | - |
| 12 | A10 | FITC | bAla | I | P | E | S | S | E | L | T | L | **A** | E | L | L | G | E | E | R | 242±31 | 1 |
| 13 | A11 | FITC | bAla | I | P | E | S | S | E | L | T | L | Q | **A** | L | L | G | E | E | R | 3158±568 | 16 |
| 14 | A12 | FITC | bAla | I | P | E | S | S | E | L | T | L | Q | E | **A** | L | G | E | E | R | >10000 | - |
| 15 | A13 | FITC | bAla | I | P | E | S | S | E | L | T | L | Q | E | L | **A** | G | E | E | R | >10000 | - |
| 16 | A14 | FITC | bAla | I | P | E | S | S | E | L | T | L | Q | E | L | L | **A** | E | E | R | 4342±1177 | 23 |
| 17 | A15 | FITC | bAla | I | P | E | S | S | E | L | T | L | Q | E | L | L | G | **A** | E | R | 953±119 | 5 |
| 18 | A16 | FITC | bAla | I | P | E | S | S | E | L | T | L | Q | E | L | L | G | E | **A** | R | 274±21 | 1 |
| 19 | A17 | FITC | bAla | I | P | E | S | S | E | L | T | L | Q | E | L | L | G | E | E | **A** | 413±39 | 2 |

**Table S1**. Alanine scanning of the N-terminal modified E6AP peptide. Seventeen single alanine mutants of E6AP peptides were synthesized to investigate the critical residues for binding activity. Substituting residues Leu9, Leu12, and Leu13 to alanine or removing the N-terminal modification significantly decreases or disrupts binding. Four residues, Glu6, Thr8, Glu11, and Gly14, showed medium alanine tolerance. Binding affinity was measured by competition BLI binding assay (N = 3). *NB: no binding. *bAla= beta-Alanine.

| Name | N-Term | Sequence | $K_D$ in nM | Ratio to wild-type |
|---|---|---|---|---|
| **WT** | - | IPESSELTLQELLGEERRAAK | 2441 ± 1334 | 1 |
| **N-FITC** | Fluorescein-5-Isothiocyanate | IPESSELTLQELLGEERR | 95±17 | 25.7 |
| N1 | 9-fluorene acetamido | IPESSELTLQELLGEERR | 10±6.8 | 244.6 |
| N2 | 1-fluorene carboxamido | IPESSELTLQELLGEERR | 13±6.3 | 188.1 |
| N3 | 1-Indane acetamido | IPESSELTLQELLGEERR | 425±78 | 5.7 |
| N4 | 9-fluorenone-2-carboxamido | IPESSELTLQELLGEERR | 79±14 | 30.9 |
| N5 | 9-fluorenone-1-carboxamido | IPESSELTLQELLGEERR | 110±21 | 22.2 |
| N6 | 9-fluorenone-4-carboxamido | IPESSELTLQELLGEERR | 102±18 | 23.9 |
| N7 | anthraquinone-2-carboxamido | IPESSELTLQELLGEERR | 27±6.3 | 90.5 |
| N8 | xanthene-9-carboxamido | IPESSELTLQELLGEERR | 75±19 | 32.6 |
| N9 | 1-anthracene carboxamido | IPESSELTLQELLGEERR | 22±7.2 | 111.1 |
| N10 | 2-anthracene carboxamido | IPESSELTLQELLGEERR | 23±9.0 | 106.3 |
| N11 | 1-adamantane carboxamido | IPESSELTLQELLGEERR | 595±152 | 4.1 |
| N12 | Triphenyl acetamido | IPESSELTLQELLGEERR | 199±37 | 12.2 |
| N13 | Diphenyl acetamido | IPESSELTLQELLGEERR | 274±51 | 8.9 |
| N14 | 1-naphthyl carboxamido | IPESSELTLQELLGEERR | 292±54 | 8.3 |
| N15 | 1,6-dihydrophenyl carboxamido | IPESSELTLQELLGEERR | 767±295 | 3.1 |
| N16 | Pentafluorophenyl carboxamido | IPESSELTLQELLGEERR | 228±74 | 10.7 |
| N17 | 6-hydroxy-2-naphthyl carboxamido | IPESSELTLQELLGEERR | 159±47 | 15.3 |
| N18 | 1-pyrenebutyl carboxamido | IPESSELTLQELLGEERR | 106±27 | 23 |
| N19 | 5-Acenaphthene carboxamide | IPESSELTLQELLGEERR | 55±12 | 44.4 |
| N20 | 7-Methoxycoumarin-4-acetamido | IPESSELTLQELLGEERR | 213±47 | 11.4 |
| N21 | 4-phenyl-phenylalanine | IPESSELTLQELLGEERR | 1412±757 | 1.7 |
| N22 | Cyclohexyl-alanine | IPESSELTLQELLGEERR | 1658±705 | 1.4 |

**Table S2**. N-terminal modified E6AP peptides. The binding affinity was measured by BLI competition assay. Error distribution is reported as the standard error of the mean (SEM) of curve fitting reported by Prism 8 software. The wild-type peptide was used as a reference. $K_D$ values are measured by BLI binding assay in a competition mode (N = 2 or 3). The ratio is calculated by dividing $K_D$ to the WT $K_D$ showed in the first row.

| Name | Sequence | C-term | $K_D$ in nM | Ratio to wild-type |
|---|---|---|---|---|
| **WT** | IPESSELTLQELLGEERRAAK | - | 2432 ± 1302 | 1 |
| **C-FITC** | Ac-IPESSELTLQELLGEERRAA | Lys (Fluorescein-5-Isothiocyanate) | 640±82 | 3.8 |
| C1 | Ac-IPESSELTLQELLGEERRAA | Lys(fluorene-9-acetamido) | 165±35 | 9.9 |
| C2 | Ac-IPESSELTLQELLGEERRAA | Lys(fluorene-1-acetamido) | 245±59 | 9.9 |
| C3 | Ac-IPESSELTLQELLGEERRAA | Lys(1-Indane acetamido) | 645±92 | 3.7 |
| C4 | Ac-IPESSELTLQELLGEERRAA | Lys(9-fluorenone-2-carboxamido) | 71±14 | 34.4 |
| C5 | Ac-IPESSELTLQELLGEERRAA | Lys(9-fluorenone-1-carboxamido) | 81±13 | 30.1 |
| C6 | Ac-IPESSELTLQELLGEERRAA | Lys(9-fluorenone-4-carboxamido) | 84±13 | 29.1 |
| C7 | Ac-IPESSELTLQELLGEERRAA | Lys(anthraquinone-2-carboxamido) | 46±5.5 | 53.1 |
| C8 | Ac-IPESSELTLQELLGEERRAA | Lys(xanthene-9-carboxamido) | 78±18 | 31.3 |
| C9 | Ac-IPESSELTLQELLGEERRAA | Lys(1-anthracene carboxamido) | 22±5.8 | 111.1 |
| C10 | Ac-IPESSELTLQELLGEERRAA | Lys(2-anthracene carboxamido) | 64±10 | 38.2 |
| C11 | Ac-IPESSELTLQELLGEERRAA | Lys(1-adamantane carboxamido) | 1459±474 | 1.6 |
| C12 | Ac-IPESSELTLQELLGEERRAA | Lys(triphenylacetamido) | 878±125 | 2.7 |
| C13 | Ac-IPESSELTLQELLGEERRAA | Lys(diphenylacetamido) | 1184±264 | 2 |
| C14 | Ac-IPESSELTLQELLGEERRAA | Lys(1-naphthyl carboxamido) | 507±65 | 4.8 |
| C15 | Ac-IPESSELTLQELLGEERRAA | Lys(Exo-norbornene-carboxamido) | 756±128 | 3.2 |

**Table S3**. C-terminal modified E6AP peptides. The binding affinity was measured by BLI competition assay. Error distribution is reported as the standard error of the mean (SEM) of curve fitting reported by Prism 8 software. The wild-type peptide was used as a reference. $K_D$ values are measured by BLI binding assay in a competition mode (N = 2 or 3). The ratio is calculated by dividing $K_D$ to the WT $K_D$ showed in the first row.

**(A)**

| Name | Sequence | Cross-link yield (%) | Conditions[a] |
| --- | --- | --- | --- |
| E0 | AcE[1]L[2]T[3]L[4]Q[5]E[6]L[7]L[8]G[9]E[10]E[11]R[12] | N/A | N/A |
| E1 | AcELTLQELLC(Phacr)EER | 0 | protein 2μM; peptide 20 μM, x1 PBS, rt 24 h |
| E2 | AcELTLQELLGC(Phacr)ER | 0 | protein 2μM; peptide 20 μM, x1 PBS, rt, 24 h |
| E3 | AcELTLQELL(Dha)EER | 74 | protein 2 μM, peptide 4 μM, x1 PBS, rt, 2h |
| E4 | AcELTLQELLG(Dha)ER | 0 | protein 2 μM, peptide 20 μM, x1 PBS, rt, 1h |
| E5 | AcELTLQELLGE(Dha)R | 0 | protein 2 μM, peptide 20 μM, x1 PBS, rt, 2h |
| E6 | AcELTLQELL(dap-acr)EER | 0 | protein 2μM; peptide 20 μM, x1 PBS, rt 2 h |
| E7 | AcELTLQELLG(dap-acr)ER | 0 | protein 2μM; peptide 20 μM, x1 PBS, rt 2 h |
| E8 | AcELTLQELLGE(dap-acr)R | 0 (42%, 3 days) | protein 2μM; peptide 20 μM, x1 PBS, rt 3 d |
| E9 | AcELTLQELLG(dab-acr)ER | 0 | protein 2μM; peptide 20 μM, x1 PBS, rt 2 h |
| E10 | AcELTLQELLGE(dab-acr)R | 0 | protein 2μM; peptide 20 μM, x1 PBS, rt 2 h |
| E11 | AcELTLQELLGE(dab-ppa)R | 27 | protein 2μM; pep 5 μM, x1 PBS, rt 2 h |

**(B)**

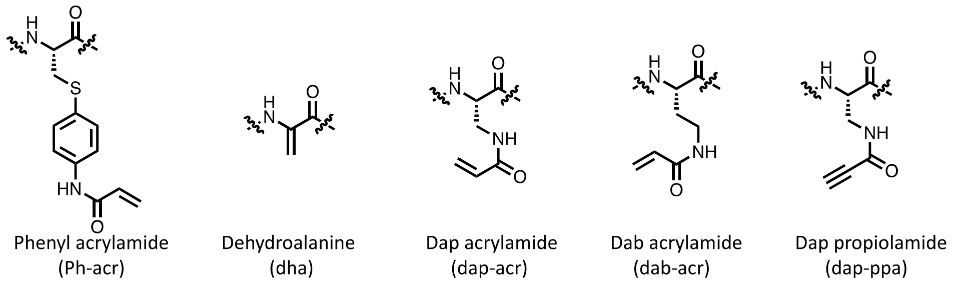


**Figure S4.** (**A**) Cross-linking experiments between MBP-16E6 and peptides E0 to E11 Reactions were performed in 1X PBS at pH = 7.4 under conditions listed for each experiment. Cross-link yield was measured by LC-MS protein deconvolution mass spectrum, and the percentage was calculated by dividing the peak area of cross-linked protein by the sum of uncross-linked and cross-linked protein peak area (N = 2). (**B**) Structures of electrophiles.

| Name | N-term | Sequence | C-term | Charge | Apparent $K_i$ in nM |
|---|---|---|---|---|---|
| **7** | 9-fluorenyl acetamido | IPESAELTLQELL(Dha)EERRAA | K(1-anthracenyl acetamido) | -3 | 23±8.1 |
| **8** | 9-fluorenyl acetamido | IPESAELTLQELL(Dha)EERRNKK | K(1-anthracenyl acetamido) | -1 | 39±6.1 |
| **9** | 9-fluorenyl acetamido | IPQSAELTLQELL(Dha)EARRNKK | K(1-anthracenyl acetamido) | 1 | 11±3.7 |
| **10** | 9-fluorenyl acetamido | IPQSAELTLQELL(Dha)QARRNKK | K(1-anthracenyl acetamido) | 2 | 16±2.6 |
| **11** | 9-fluorenyl acetamido | IPQSAELTLQELL(Dha)QARRKK | K(1-anthracenyl acetamido) | 2 | 11±4.9 |
| **12** | 9-fluorenyl acetamido | IPQSAELTLQELL(Dha)QRRKK | K(1-anthracenyl acetamido) | 2 | 20±5.4 |
| **13** | 9-fluorenyl acetamido | IPQSAELTLQELL(Dha)RRKK | K(1-anthracenyl acetamido) | 2 | 17±3.9 |
| **13-3L3A** | 9-fluorenyl acetamido | IPQSAELTAQEAA(Dha)RRKK | K(1-anthracenyl acetamido) | 2 | n.b. |

**Figure S5.** Sequence table of E6AP-mimicking peptides designed to increase positive charge. Apparent Ki is determined by BLI. Peptides were in competition with immobilized 1-Biotin following a 30 min incubation with E6. Dha: dehydroalanine.

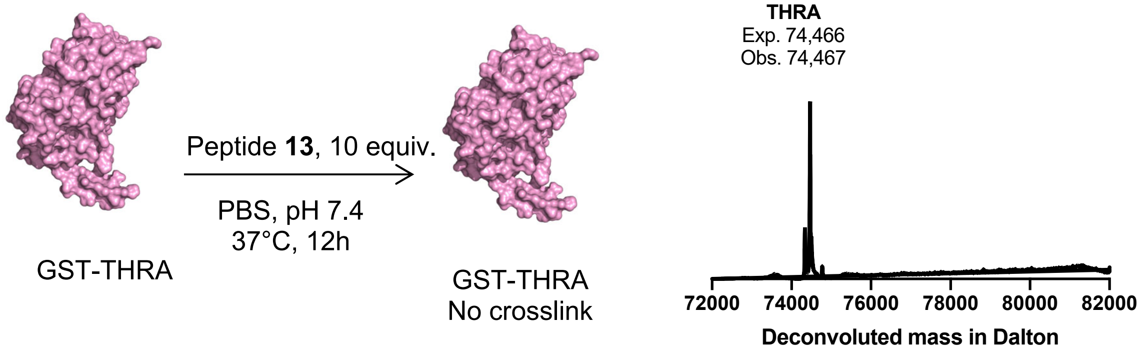


**Figure S6**. **13** showed no observable crosslink to THRA. The crosslinking reaction was performed with 1 μM THRA and 10 μM **13** at 37 °C for 12 h. PDB code of THRA is 1nav.

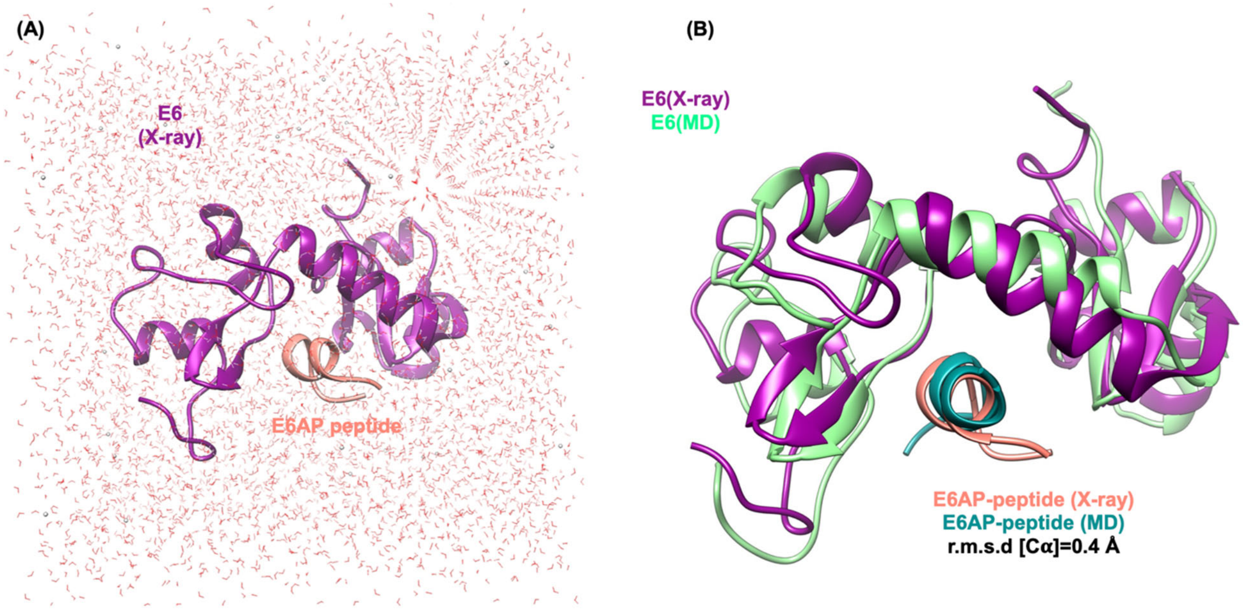


**Figure S7. The native E6AP LXXLL peptide forms a stable alpha helix when in complex with 16E6. (A)** E6AP peptide (salmon) bound to 16E6 protein (purple) with surrounding water molecules (red lines) and ions (white spheres). **(B)** Comparison of X-ray structure of 16E6 (green) bound to E6AP (salmon) (PDB ID: 4XR8) with relaxed 16E6(purple)-E6AP peptide (dark green) complex after 1.1 μs MD simulation. P53 and MBP were omitted for clarity. The RMSD was calculated for the Cα atoms.

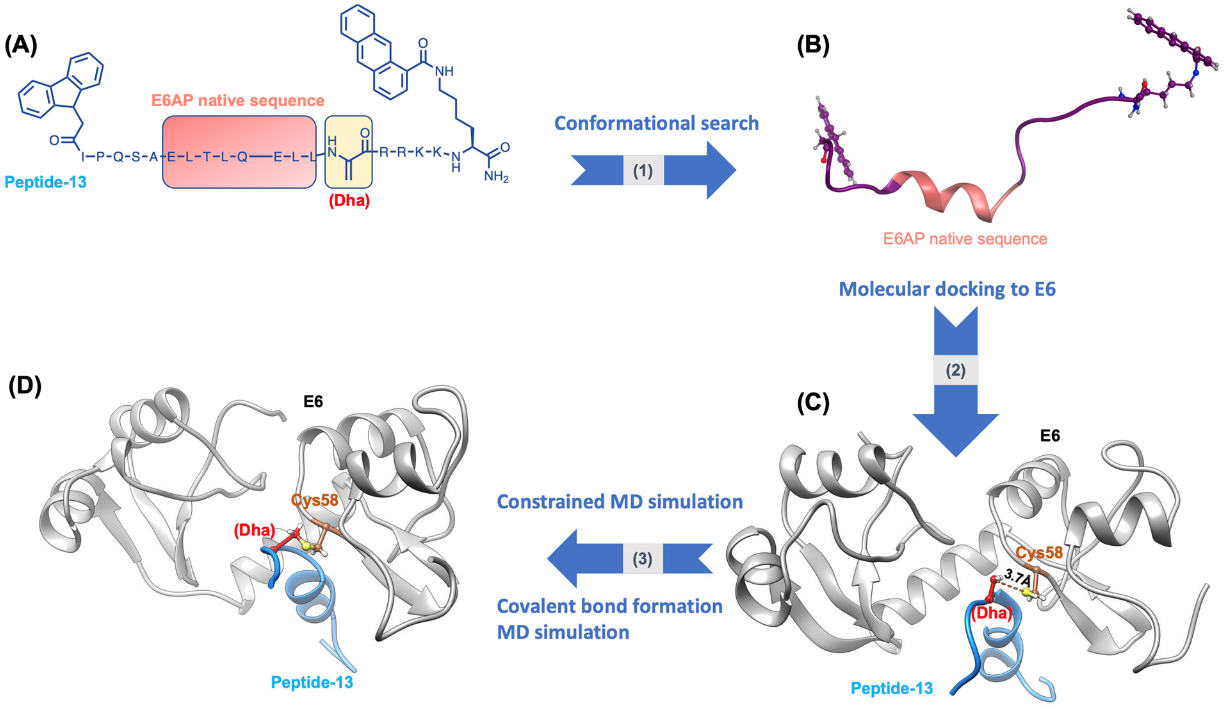


**Figure S8. Structural modeling of 16E6 bound to Peptide-13.** (**A**) Structure of Peptide-13. (**B**) E6AP LXXLL peptide native core adopts an alpha helical conformation in the peptide 13 structural model obtained by computational conformational sampling. (**C**) Molecular docking of Peptide-13 (blue) to 16E6 protein (gray). (**D**) Molecular modeling of 16E6 (gray) and peptide 13 (blue) in complex with the covalent thioester bond between the peptide Dha warhead of peptide 13 and Cys58 of 16E6 highlighted.

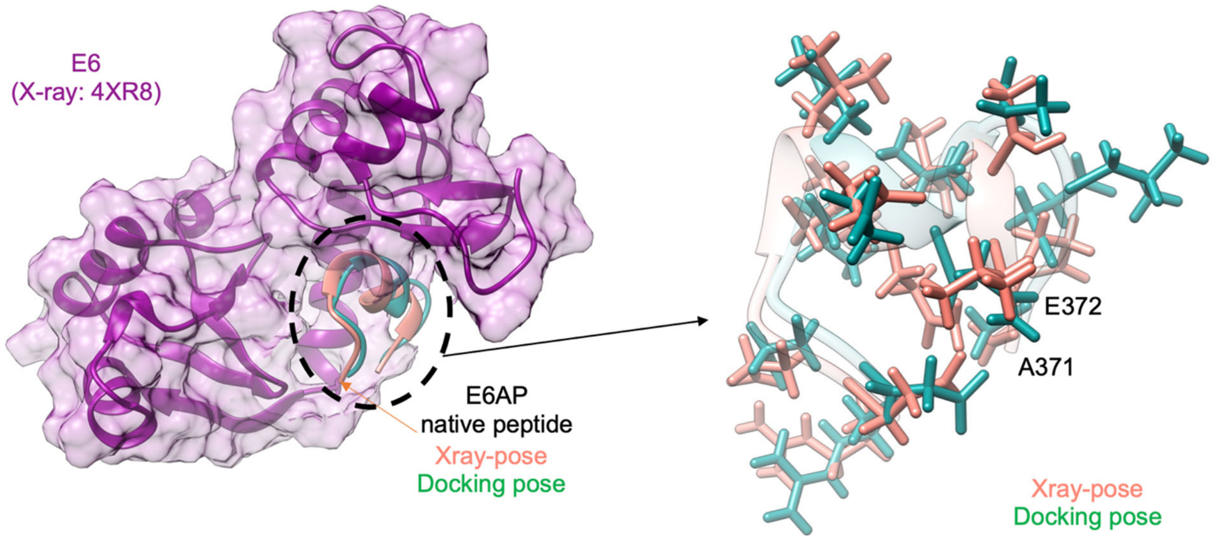


**Figure S9. Molecular docking of the native E6AP peptide in the 16E6 (left).** The MOE molecular docking procedure accurately describes the interactions between E6 and E6AP native peptide (right).

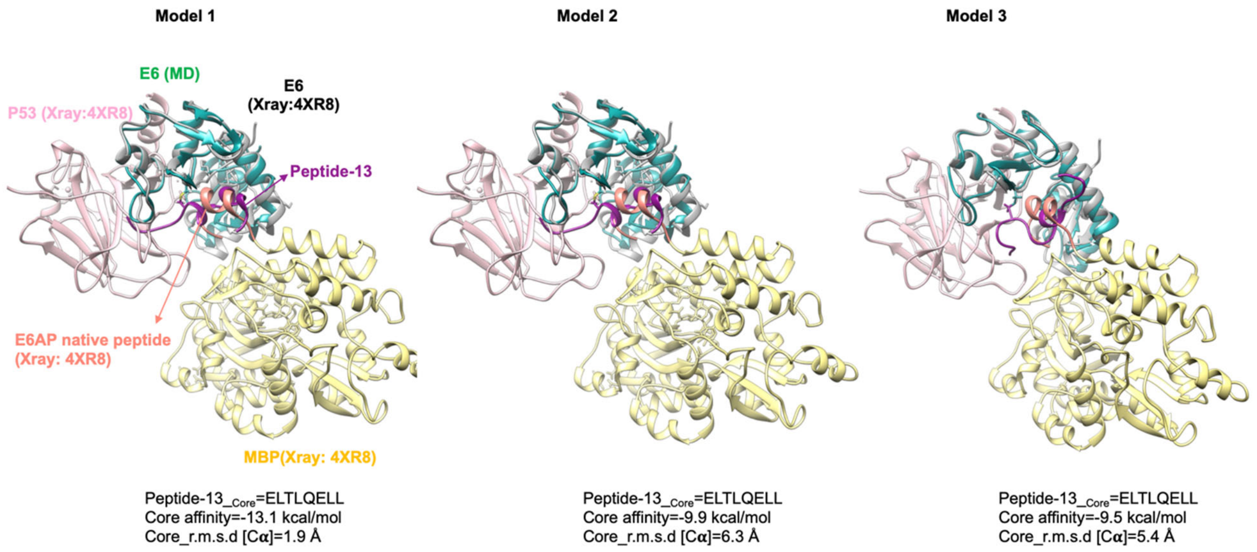


**Figure S10. Molecular modeling of 16E6 and peptide 13 complexes.** Complexes are ordered from highest binding affinity (mode 1) to lowest (mode 3) based on RMSD of peptide 13 core sequence compared to the native E6AP LXXLL peptide in PDB: 4XR8.

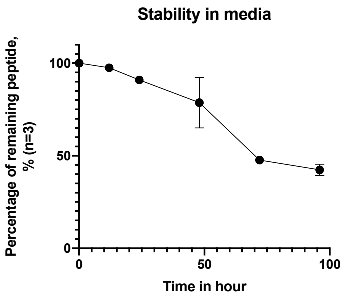


**Figure S11. Reactide 13 stability in media**. Ten μM peptide was incubated in RPMI with 10% FBS at 37 °C for the indicated time. Proteins were removed from the mixture and analyzed by LC-MS. Percentage of the remaining peptide was estimated by dividing the area under extract ion chromatogram (EIC) to the area of time zero (N = 3).

CLUSTAL O(1.2.4) multiple sequence alignment

```
                                                                                58
UNIPROT:VE6_HPV16       RTAMFQDPQERPRKLPQLCTELQTTIHDIILECVYCKQQLLRREVYDFAFRDLCIVYRDG 64
UNIPROT:VE6_HPV39       -MARFHNPAERPYKLPDLCTTLDTTLQDITIACVYCRRPLQQTEVYEFAFSDLYVVYRDG 59
UNIPROT:VE6_HPV18       -MARFEDPTRRPYKLPDLCTELNTSLQDIEITCVYCKTVLELTEVFEFAFKDLFVVYRDS 59
UNIPROT:VE6_HPV45       -MARFDDPKQRPYKLPDLCTELNTSLQDVSIACVYCKATLERTEVYQFAFKDLCIVYRDC 59
UNIPROT:VE6_HPV51       ---MFEDKRERPRTLHELCEALNVSMHNIQVVCVYCKKELCRADVYNVAFTEIKIVYRDN 57
UNIPROT:VE6_HPV56       MEPQFNNPQERPRSLHHLSEVLEIPLIDLRLSCVYCKKELTRAEVYNFACTELKLVYRDD 60
UNIPROT:VE6_HPV52       ---MFEDPATRPRTLHELCEVLEESVHEIRLQCVQCKKELQRREVYKFLFTDLRIVYRDN 57
UNIPROT:VE6_HPV33       ---MFQDTEEKPRTLHDLCQALETTIHNIELQCVECKKPLQRSEVYDFAFADLTVVYREG 57
UNIPROT:VE6_HPV58       ---MFQDAEEKPRTLHDLCQALETSVHEIELKCVECKKTLQRSEVYDFVFADLRIVYRDG 57
UNIPROT:VE6_HPV31       ---MFKNPAERPRKLHELSSALEIPYDELRLNCVYCKGQLTETEVLDFAFTDLTIVYRDD 57
UNIPROT:VE6_HPV35       ---MFQDPAERPYKLHDLCNEVEESIHEICLNCVYCKQELQRSEVYDFACYDLCIVYREG 57
UNIPROT:B9UPD9_HPV59    -MARFEDPTQRPYKLPDLSTTLNIPLHDIRINCVFCKEELQEREVFEFAFNDLFIVYRDC 59
                            *.:   :* .* .*.   ::   :: : ** *:  *   :* ..   :: :***:
```

* Conserved sequence(identical)
: Conservative mutation
. Semi-conservative mutation
()Non-conservative mutation
- Gap

**Figure S12**. Protein sequence alignment of High-risk type HPV E6 using CLUOSTAL O (1.2.4) sequence alignment tool. The alignment includes for the following high-risk subtypes: 18, 31, 33, 35, 39, 45, 51, 52, 56, 58, 59, 66, and 68. For example, in HPV16E6 (48-64) and HPV18E6 (43-59), the Phe53 of HPV18 E6 aligns with the Cys58 of HPV16 E6. This Phe52 residue made HPV18E6 insensitive to cysteine reactive bind-and-react inhibition strategy.

### Methods and Materials

#### 1.1 Materials

H-Rink Amide-ChemMatrix resin was purchased from PCAS BioMatrix Inc. Amino acids: Fmoc-Ala-OH, Fmoc-β-Ala-OH, Fmoc-Asn(Trt)-OH, Fmoc-Asp(*t*Bu)-OH, Fmoc-Cys(Trt)-OH, Fmoc-Glu(*t*Bu)-OH, Fmoc-Leu-OH, Fmoc-Lys(Boc)-OH, Fmoc-Phe-OH, Fmoc-Pro-OH, Fmoc-Ser(*t*Bu)-OH, Fmoc-Thr(*t*Bu)-OH, Fmoc-Trp(Boc)-OH, Fmoc-Tyr(*t*Bu)-OH, and Fmoc-Val-OH were purchased from Novabiochem (Billerica, MA). Other amino acids: Fmoc-Lys(ivdde)-OH, Fmoc-Lys(Alloc)-OH, Fmoc-Lys(Fmoc)-OH were purchased from Novabiochem (Billerica, MA). Palladium tetrakistriphenylphosphine(0) ($Pd(PPh_3)_4$) was purchased from Sigma Aldrich. Reagents used in solid phase peptide synthesis: Piperidine (ReagentPlus; 99%), formic acid (≥ 98%) were purchased from Sigma-Aldrich (St. Louis, MO). Diisopropylethylamine (DIEA; biotech. grade; 99.5%) was purchased from Millipore Sigma and purified by a Seca Solvent Purification system from Pure Process Technology (Nashua, NH). Reagents (cleavage): Trifluoroacetic acid (TFA; for HPLC, ≥99%), triisopropylsilane (TIPS; 98%) were purchased from Sigma-Aldrich (St. Louis, MO). Reagents used in peptide post-synthesis modifications: Acetic anhydride (≥98%) was purchased from Sigma-Aldrich (St. Louis, MO), 5-TAMRA (5-carboxytetramethylrhodamine) and Biotin-PEG4-carboxylic acid from ChemPep. Bovine serum albumin (BSA) from VWR, PA. Recombinant Human Thyroid Hormone Receptor alpha 1 protein (THRA) from Abcam, UK.

Media (RPMI-1640 ATCC modified, EMEM and MEM), fetal bovine serum (FBS), penicillin-streptomycin, 10× PBS, and 0.25% trypsin-EDTA were obtained from Gibco. Culture flasks, plates, and serological pipettes were obtained from Fisher Scientific. Fixing reagent: 37% formaldehyde, dilute 10 times in 1x PBS (Sigma). Dye: Wheat Germ Agglutinin (WGA) Alexa Fluor™ 350 Conjugate (W11263, Thermo).

#### 1.2 Fast flow synthesis of peptides

H-Rink Amide-ChemMatrix resin (200 mg, 0.49 mmol/g, 0.10 mmol) was used to prepare peptide-α-carboxamides. Peptides containing noncanonical amino acids were prepared by manual SPPS. Peptides without noncanonical amino acids were prepared by fully automated SPPS[1]. Upon completion, resins were washed with dichloromethane (DCM) three times and dried under reduced pressure.

#### 1.3 Automated flow peptide synthesis (AFPS) set-up

All peptides were synthesized on automated-flow systems built in the Pentelute lab (“Amidator” and “Peptidator”), which are similar to the published AFPS system. The synthesis conditions were published previously[2]:

The following settings were used for protein synthesis: flowrate = 40 mL/min, temperature = 90°C (loop) and 85–90°C (reactor). The 50 ml/min pump head pumps 400 μL of liquid per pump stroke; the 5 mL/min pump head pumps 40 μL of liquid per pump stroke. The standard synthetic cycle involves a first step of prewashing the resin at elevated temperatures for 60 s at 40 mL/min. During the coupling step, three HPLC pumps are used: a 50 mL/min pump head pumps the activating agent, a second 50 ml/min pump head pumps the amino acid and a 5 mL/min pump head pumps DIEA. The first two pumps are activated for 8 pumping strokes to prime the coupling agent and amino acid before the DIEA pump is activated. The three pumps are then actuated together for a period of 7 pumping strokes, after which the activating agent pump and amino acid pump are switched using a rotary valve to select DMF. The three pumps are actuated together for a final 8 pumping strokes, after which the DIEA pump is shut off, and the other two pumps continue to wash the resin for another 40 pump strokes. During the deprotection step, two HPLC pumps are used. Using a rotary valve, one HPLC pump selects deprotection stock solution and DMF. The pumps are activated for 13 pump strokes. Both solutions are mixed in a 1:1 ratio. Next, the rotary valves select DMF for both HPLC pumps, and the resin is washed for an additional 40 pump strokes. The coupling–deprotection cycle is repeated for all additional monomers.

**1.4 Method for peptide acetylation**

A 100 mg portion of peptidyl resin from section 1.3 was placed into a 5 mL Torviq fritted syringe and subsequently swelled in DMF. After removing DMF, a solution of $Ac_2O$, DIEA, and DMF (2 mL, 85:315:1600, v/v) were added to the peptidyl resin. The resulting mixture was occasionally agitated for 45 min. After draining the solution, the remaining resin was washed with DMF three times, DCM three times, and dried under reduced pressure.

**1.5 Method for Alloc deprotection**

Peptidyl resin (~10 µmol theoretical loading) was washed with DCM (3 × 5 mL) and then treated with $Pd(PPh_3)_4$ (11.0 mg, 10 µmol, 1 equiv) in DCM/piperidine (8:2, 1 mL) for 30 minutes at room temperature under exclusion of light. The resin was then drained and washed with DCM (3 × 5 mL).

**1.6 Method for biotin labeling**

Peptidyl resin (~10 µmol theoretical loading) was loaded into a fritted syringe (6 mL), swollen in DMF (4 mL) for 5 minutes, and then drained. Biotin-PEG4-propionic acid (Biotin-PEG4-OH, 22 mg, 50 µmol, 5 equivalents) and HATU (17 mg, 45 µmol, 4.5 equivalents) were dissolved in DMF (500 µL), activated with DIEA (19 mg, 26 µL, 150 µmol), added to the peptidyl resin and incubated for 30 minutes under exclusion of light. After this time, the resin was drained, washed with DMF (3 × 5 mL), and stored until cleavage.

**1.7 Method for TAMRA labeling**

Peptidyl resin (~ 10 µmol theoretical loading) was loaded into a fritted syringe (6 mL), swollen in DMF (4 mL) for 5 minutes, and then drained. 5-Carboxytetramethylrhodamine (5-TAMRA, 22 mg, 50 µmol, 5 equivalents) and HATU (17 mg, 45 µmol, 4.5 equivalents) were dissolved in DMF (500 µL), activated with DIEA (19 mg, 26 µL, 150 µmol), added to the peptidyl resin and incubated for 30 minutes under exclusion of light. After this time, the resin was drained, washed with DMF (3 × 5 mL), and stored until cleavage.

**1.8 Method for FITC labeling**

Peptidyl resin (~ 10 µmol theoretical loading) was loaded into a fritted syringe (6 mL), swollen in DMF (4 mL) for 5 minutes, and then drained. 5-Carboxytetramethylrhodamine (Fluorescein isomer I, 22 mg, 50 µmol, 5 equivalents) was dissolved in DMF (500 µL), activated with DIEA (19 mg, 26 µL, 150 µmol), added to the peptidyl resin and incubated for 30 minutes under exclusion of light. After this time, the resin was drained, washed with DMF (3 × 5 mL) and stored until cleavage.

**1.9 Cleavage of peptides**

The synthesized peptide was cleaved from the resin and globally deprotected by treating the peptidyl resin with a cleavage cocktail containing 94% TFA, 2.5% water, and 2.5% TIPS (*v/v*), for 2 h at room temperature. TFA was removed under a gentle stream of nitrogen gas, and the crude peptide was precipitated by adding cold $Et_2O$ (-80°C). After centrifugation at 3220 rcf for 3 min, the supernatant was removed, and the precipitated peptide was triturated three times with cold $Et_2O$. The resulting material was dissolved in 50% MeCN in water with 0.1% TFA and lyophilized as crude.

### 1.10 Pd Mediated Conjugation

In a 50 mL falcon tube: Dissolve peptide (5.0 mg, 2.75 mmol, 1.0 equiv) in water (8.5 mL) and 500 mM HEPES (1.5 mL, pH = 8.0). In a 20 mL glass vial: Dissolve Pd OAC (5.0 mg, 6.19 mmol, 2.25 equiv) in MeCN and add to peptide solution over 30 seconds (final volume = 15 mL, final peptide concentration = 184 μM). Mix by vortexing and let stand for 20 minutes. Add 10 mL AcOH and 30 mL $H_2O$ and mix and then purify by reversed phase flash chromatography using a Sfär Bio C18 D (300 Å 20 μm, 6 g) column (mobile phase 5% MeCN/$H_2O$ to 55% MeCN in $H_2O$ + 0.1% TFA).

### 1.11 Dehydroalanine formation

In a 1.5 mL microcentrifuge tube, cysteine-containing peptide (7 mg) was dissolved in DMF (0.5 mL), and to this, a 10 mg/mL potassium carbonate solution (3.1 mg, 22.3 μmol, 5 equiv) was added. To this mixture, a 10 mg/mL solution of Diethylmeso-2,5-dibromoadipate (151 μL, 1.1 equiv) was added. The mixture was mixed by vortexing for 5 seconds and allowed to react for 4 h. The reaction mixture was diluted with 5% MeCN in $H_2O$ + 0.1% TFA and purified by reversed-phase HPLC (Zorbax 300SB-C3, 300Å, 5 μm, 9.4 mm x 250 mm) Mobile phase 5% to 55% MeCN in $H_2O$ + 0.1% TFA.

### 1.12 General procedure for Cross-linking MBP-16E6 with Covalent Peptides

A solution of cross-linker (250 μM solution in 10% DMSO/$H_2O$, 2 to 10 equiv, 0.66 μL) was diluted with $H_2O$ (30 μL) and 10× PBS (4.1 μL, pH = 7.4). A 30 μM solution of MBP-6 (2.4 μL, 1 equiv) and then incubated at room temperature for 2 h. An 8 μL aliquot of the reaction mixture was removed and quenched with 92 μL of 50:50 MeCN/$H_2O$ + 0.1% TFA and analyzed by LC/MS. Yields were obtained by extracting all protein-containing species' total ion current (TIC) spectra in the chromatogram utilizing Agilent MassHunter Bioconfirm Software 10.0. The extracted chromatograms were deconvoluted utilizing a maximum entropy algorithm, and the abundance of each species was determined using total ion count. $\%\text{yield} = \frac{P_c}{P_c+P_0} \times 100$ where $P_c$ is the peak area of the peptide-protein conjugate, and $P_0$ is the peak area of the unmodified protein

### 1.13 Purification of the crude peptide.

Crude peptides were purified by a Biotage Selekt flash purification system. Water with 0.1% TFA (solvent A) and MeCN with 0.1% TFA (solvent B) was utilized as mobile phases for purifications. The crude peptide was dissolved in a minimal amount of 10% MeCN in water with 0.1% TFA and then loaded onto a 10 g Biotage SNAP Bio C4 20 μm column. The purification was performed using a gradient as follows: 10% B for 2 column volume (CV), the linear ramp from 30% B to 50% B for 20 CV, 25 mL/min flow rate.

### 1.14 Method for LC-MS characterization.

LC-MS characterizations were carried out using an Agilent 6550 quadrupole time-of-flight LC-MS. Total ion current (TIC) chromatograms were plotted. Mass spectra were integrated over the principal TIC peaks. High-performance liquid chromatography was done by the following methods: (solvent A: water with 0.1% formic acid; solvent B: MeCN with 0.1% formic acid).

**Method A:** Column: Phenomenex Jupiter C4 column (1.0 × 150 mm, 5 μm particle size, 300 Å pore size) Gradient: 1% B (0-2 min), linearly ramp from 1% B to 91% B (2-8 min). The flow rate is 100 μL/min. MS acquisition is from 2 to 8 min.

**Method B:** Column: Phenomenex Jupiter C4 column (1.0 × 150 mm, 5 μm particle size, 300 Å pore size) Gradient: 1% B (0-2 min), linearly ramp from 1% B to 61% B (2-12 min), 61% B to 95% B (11-16 min). The flow rate is 100 μL/min. MS acquisition is from 4 to 12 min.

**Method C:** Column: Agilent Zorbax 300SB C3 column (2.1 × 150 mm, 5 μm particle size, 300 Å pore size) Gradient: 1% B (0-2 min), linearly ramp from 1% B to 91% B (2-12 min), 91% B to 91%

B (12-13 min). The flow rate is 500 µL/min. MS acquisition is from 4 to 12 min.

#### 1.15 Direct binding measurement by BLI

Streptavidin sensors were soaked in blocking buffer (1× PBS supplemented with 0.05% Tween-20 and 1 mg/mL bovine serum albumin) for 5 min. After immobilizing the **1-Biotin** peptide (200 nM onto streptavidin sensors, serial dilutions of E6 in blocking buffer were analyzed for binding. The response was recorded at equilibrium after 2 min. Association lasted for 100 seconds, and dissociation lasted for another 120 seconds. The curve was reported by GatorPlus software and replotted by Prism 8 software.

#### 1.16 Competitive binding assay by BLI

A competition binding assay was performed as described below using GatorPlus bio-layer interferometry(GatorBio) to estimate the binding affinity of peptides.

*Calibration curve:* Streptavidin sensors were soaked in blocking buffer (PBS supplemented with 0.05% Tween-20 and 1 mg/mL bovine serum albumin) for 5 min. After immobilizing the PEG4-Biotinylated **1** peptide (200 nM of Biotin-PEG4-IPESSELTLQELLGEER) onto streptavidin sensors, 1:1 serial dilutions from 1000 nM of E6 in blocking buffer were analyzed for binding (All concentrations: 1000nM, 500 nM, 250 nM, 125 nM, 62 nM, 31nM, 15nM, and 7.8nM). The response was recorded at equilibrium after 2 min. A curve of sensor response (nm) vs. E6 concentration (nM) was generated to calibrate the free E6 concentration in the solution observed in the competition assay. The curve was generated using Prism 8 software.

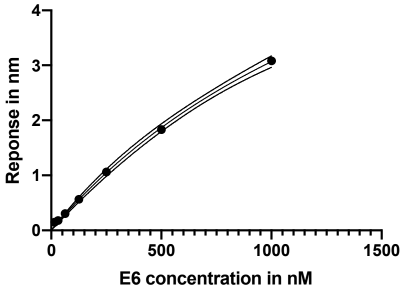


*Competition assay.* Various concentrations of peptides were incubated in wells with E6 protein in the blocking buffer for 30 min. The PEG4-Biotinylated **1** peptide was immobilized onto streptavidin sensors and dipped into preincubated sample wells. The association events were measured at 30°C, 1,000 rpm. Response at equilibrium after 2 min was recorded. Based on the binding response (nm) values, the concentration of 'free' E6 was interpolated for each sample using the calibration curve. The apparent dissociation constant, $K_D$, can be obtained from the non-linear regression analysis using the equation:

$[\mathrm{Y}] = 0.5 \times [b - \mathrm{K}_d - [\mathrm{X}] + \sqrt{([\mathrm{X}] + \mathrm{K}_d - b)^2 + 4b \times \mathrm{K}_d}\,]$ , where [Y] is the free [E6] in nM, [X] is the total [peptide] in nM, $K_D$ is the binding dissociation constant to be fitted by the equation, and b is the maximal possible E6 concentration to be fitted by the equation. By fitting the free [E6] and [peptide] to the equation, a binding constant with a fitting error was generated by Prism 8 software.

### 1.17 Protein expression and purification methods

### 1.17.1 E6AP

***Expression and Purification Method:***
E6AP(1-875) protein with a C-terminal TEV-6xHis-Avi sequence was subcloned into a pFastBac1 vector (Genscript). Bacmid and viruses of E6AP prepared as described by vendor's instructions (Invitrogen Version A, A10606) were amplified in *Sf9* cells (ThermoFisher, cat no. 11496-015). P2 viruses at 2 uL/mL virus to media were used to infect *Sf21* cells for protein expression. Cells were harvested 48 hours post-infection.

*Sf21* cells were lysed by French Press in 50 mM HEPES pH 7.5, 500 mM NaCl, 5 mM Imidazole, 5% glycerol, 1 mM PMSF, and cOmplete Protease Inhibitor (Roche, cat no. 53002800). The supernatant was collected after centrifugation at 39,800 RCF for 30 minutes and loaded onto Ni Resin (Bestchrom, cat no. AA0053) and washed with 10 CVs of 50 mM HEPES pH 7.5, 500 mM NaCl, 5% glycerol, 1mM PMSF, and 20 mM imidazole before eluting with the same buffer supplemented with 500mM imidazole. The elution was diluted five-fold with 50 mM Tris-HCl pH 7.5 before loading onto a Mono Q 10/100 GL column (Cytiva, Cat no. 17516701) and eluted with a 20 CV linear gradient (Buffer A: 50 mM Tris-HCl (pH 7.5), 100 mM NaCl, 5% glycerol, 1 mM PMSF; Buffer B: 50 mM Tris-HCl (pH 7.5), 1 M NaCl, 5% glycerol, 1mM PMSF). Fractions containing E6AP as determined by SDS-PAGE and Coomassie staining were pooled and concentrated using Amicon Centrifugal Filters (cat no. R1SB42368) and loaded onto a HiLoad 16/600 Superdex 200pg column (Fisher Scientific, cat no. 28989335) equilibrated in 25 mM HEPES pH 7.5, 150 mM NaCl, 1 mM TCEP.

***Sequence:***
MEKLHQCYWKSGEPQSDDIEASRMKRAAAKHLIERYYHQLTEGCGNEACTNEFCASCPTFLR
MDNNAAAIKALELYKINAKLCDPHPSKKGASSAYLENSKGAPNNSCSEIKMNKKGARIDFKDVT
YLTEEKVYEILELCREREDYSPLIRVIGRVFSSAEALVQSFRKVKQHTKEELKSLQAKDEDKDED
EKEKAACSAAAMEEDSEASSSRIGDSSQGDNNLQKLGPDDVSVDIDAIRRVYTRLLSNEKIETA
FLNALVYLSPNVECDLTYHNVYSRDPNYLNLFIIVMENRNLHSPEYLEMALPLFCKAMSKLPLAA
QGKLIRLWSKYNADQIRRMMETFQQLITYKVISNEFNSRNLVNDDDAIVAASKCLKMVYYANVV
GGEVDTNHNEEDDEEPIPESSELTLQELLGEERRNKKGPRVDPLETELGVKTLDCRKPLIPFEE
FINEPLNEVLEMDKDYTFFKVETENKFSFMTCPFILNAVTKNLGLYYDNRIRMYSERRITVLYSLV
QGQQLNPYLRLKVRRDHIIDDALVRLEMIAMENPADLKKQLYVEFEGEQGVDEGGVSKEFFQL
VVEEIFNPDIGMFTYDESTKLFWFNPSSFETEGQFTLIGIVLGLAIYNNCILDVHFPMVVYRKLMG
KKGTFRDLGDSHPVLYQSLKDLLEYEGNVEDDMMITFQISQTDLFGNPMMYDLKENGDKIPITN
ENRKEFVNLYSDYILNKSVEKQFKAFRRGFHMVTNESPLKYLFRPEEIELLICGSRNLDFQALEE
TTEYDGGYTRDSVLIREFWEIVHSFTDEQKRLFLQFTTGTDRAPVGGLGKLKMIIAKNGPDTER
LPTSHTCFNVLLLPEYSSKEKLKERLLKAITYAKGFGMLENLYFQGHHHHHHGLNDIFEAQKIEW
HE*

***QC: LC-MS, MALS and Gel***

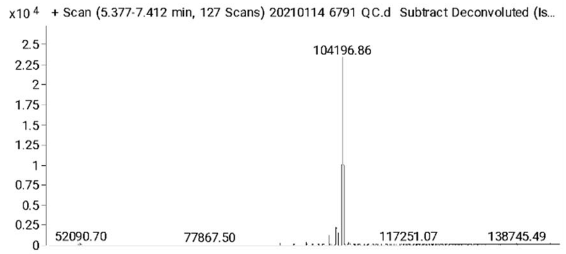


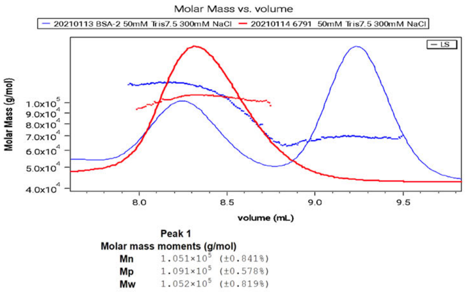


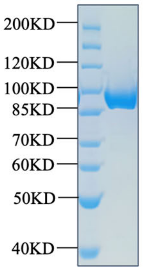


1.17.2 **MBP-16E6**

***Expression and Purification Method:***

HPV16E6(8-158) with a N-terminal HisMBP and four cysteine to serine point mutations to increase solubility was subcloned into a pET45b vector (Genscript). The plasmid was transformed into BL21(DE3) competent cells (NEB, Cat no. C2527H). The cells were grown in LB media supplemented with 50 ug/mL carbenicillin shaking at 190 RPM in a 37°C incubator to an optical density of 0.6 at 600 nm before induction with IPTG at a final concentration of 0.5 mM. Protein expression was allowed to proceed for 16 hours shaking at 190 RPM in an 18°C incubator before harvest.

BL21(DE3) cells were resuspended in 50 mM Tris-HCl pH 7.5, 500 mM NaCl, 2 mM DTT, and cOmplete Protease Inhibitor before lysis by sonification. The supernatant was collected after centrifugation at 30,000 RCF for 45 minutes and loaded onto a MBPTrap HP column (Cytiva, cat no. 28918779) equilibrated in the same resuspension buffer and washed for 30 CV. The protein was eluted with 50 mM Tris-HCl pH 7.5, 500 mM NaCl, 2 mM DTT, and 15 mM maltose. The elution was further purified by size exclusion chromatography on a HiLoad 26/600 Superdex 200pg column (Cytiva, cat no. 28989336) equilibrated in PBS supplemented with 1 mM DTT.

***Sequence:***
MAHHHHHHPMKIEEGKLVIWINGDKGYNGLAEVGKKFEKDTGIKVTVEHPDKLEEKFPQVAAT
GDGPDIIFWAHDRFGGYAQSGLLAEITPDKAFQDKLYPFTWDAVRYNGKLIAYPIAVEALSLIYN
KDLLPNPPKTWEEIPALDKELKAKGKSALMFNLQEPYFTWPLIAADGGYAFKYENGKYDIKDVG
VDNAGAKAGLTFLVDLIKNKHMNADTDYSIAEAAFNKGETAMTINGPWAWSNIDTSKVNYGVTV
LPTFKGQPSKPFVGVLSAGINAASPNKELAKEFLENYLLTDEGLEAVNKDKPLGAVALKSYEEE
LAKDPRIAATMENAQKGEIMPNIPQMSAFWYAVRTAVINAASGRQTVDEALKDAQTNSSSNNN
NNNNNNNPMSENLYFQGAMFQDPQERPRKLPQLCTELQTTIHDIILECVYCKQQLLRREVYDF
AFRDLCIVYRDGNPYAVCDKCLKFYSKISEYRHYSYSLYGTTLEQQYNKPLSDLLIRCINCQKPL
SPEEKQRHLDKKQRFHNIRGRWTGRCMSCSRSSRTRRETQL*

***QC: LC-MS and Gel of MBP 16E6 4C4S***

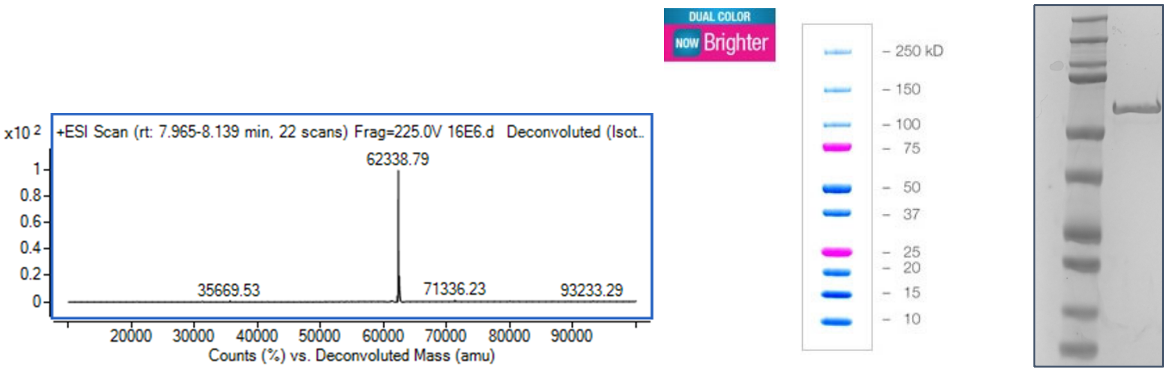


***QC: LC-MS and Gel of MBP 16E6 4C4S C58S***

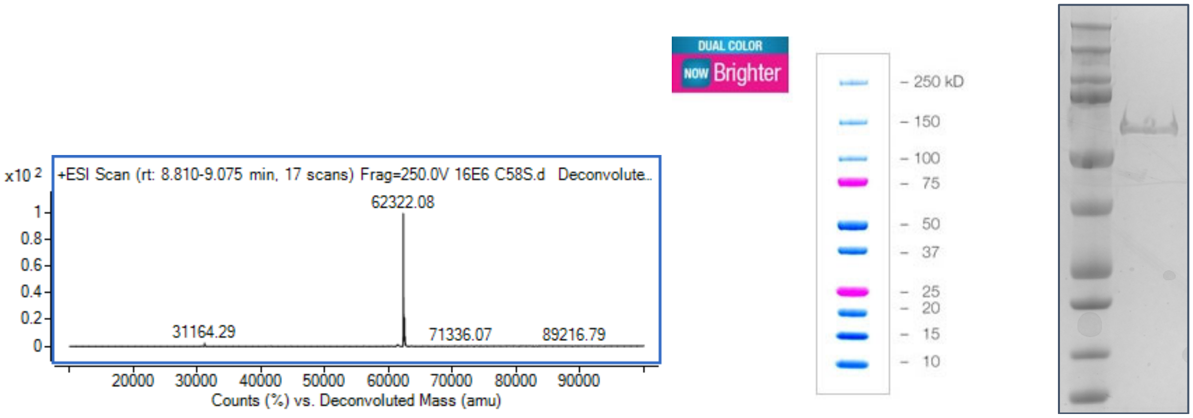


#### 1.17.3 **SUMO-MDM2**

Recombinant SUMO-MDM2 was expressed as previously reported[3].

In brief, the 25-109MDM2 gene was purchased from Addgene (pGEX-4T MDM2 wild type (WT), 16237). The SUMO tag was incorporated using the Champion™ pET SUMO Expression System (Invitrogen, CA). SUMO-25-109MDM2 was expressed in Rosetta (DE3) pLysS cells. The bacteria were inoculated to reach OD600 = 0.5 at 37°C, induced with 0.4 mM IPTG for 4 hours, and pelleted. Approximately 1 L broth produced 10 g cell pellet, which was resuspended in 30 mL of 50 mM Tris-HCl, 150 mM NaCl, pH 7.5 buffer containing 40 mg lysozyme, 1 mg Roche DNAase I, and one tablet of Roche protease inhibitor cocktail, and then sonicated for three times for 20 s. The suspension was then centrifuged at 30,000 rcf for 30 min to clarify the lysate. The supernatant was loaded onto a 5 mL HisTrap FF crude Ni-NTA columns (GE Healthcare, UK), and washed sequentially with 30 mL of 20 mM Tris-HCl, 150 mM NaCl, pH 8.5 and then 30 mL 20 mM Tris-HCl 150 mM NaCl, 80 mM imidazole pH 8.5. The protein was eluted with 10 mL 20 mM Tris-HCl, 500 mM imidazole, 500 mM NaCl, pH 8.5. The eluted protein was buffer exchanged into 20 mM Tris-HCl, 50 mM NaCl, pH 8.5 using a HiPrep 26/10 Desalting column (GE Healthcare, UK). The resulting protein solution was purified the same day using a 5 mL HiTrap Q HP (GE Healthcare, UK) anion exchange columns with a linear NaCl gradient (B% graded from 5% to 25%, B = 20 mM Tris-HCl, 500 mM NaCl, pH 8.5). Fractions containing pure SUMO-25-109MDM2, as determined by SDS-PAGE, were collected, and concentrated to 0.5 mg/mL using a 10,000 Da nominal molecular weight limit Amicon Ultra-15 Centrifuge Filter Unit (EMD Millipore), and immediately flash frozen and stored at −80 °C.

### 1.18 Complex disruption assay

E6AP protein was biotinylated by the commercial BirA kit (AVIDITY, CA). In brief, 40 µL of E6AP was added directly into a tube containing 20 µl of 1 mg/ml BirA biotin-protein ligase. Then the solution was added with 5 µL of BiomixB (10× concentration: 100 mM Adenosine 5′-triphosphate, 100 mM MgOAc, 500 µM d-biotin), followed by the addition of 5 µL of BIO200 (d-biotin; 10× concentration: 500 µM). The whole mixture was incubated at 4°C for 16 hrs, followed by a spinning filter through 0.2 µm. Excess biotin was purified away on Superdex200 or AdvanceBio 150mm (Agilent, MA) equilibrated with PBS + 1mM DTT. Biotin-E6AP fractions were collected and frozen at -80°C before use. 50 µL of 12 µM MBP-16E6 in PBS was added with 0.5 µL of 10 mM reactive peptide of interest (final concentration 100 µM). The mixture was incubated at 4°C for 16 hrs. One µM of Biotin-E6AP protein was immobilized onto streptavidin sensors soaked in a blocking buffer. The sensors were then dipped into the mixture of MBP-16E6 and reactive peptide. The signal response was recorded.

### 1.19 Structural modeling of 16E6-bound Peptide-13

Our Peptide-13 consists of a flexible N-terminal region {*9-fluorenyl acetamido*-IPQSA}, the native E6AP core {ELTLQEELL}, and flexible C-terminal {Dha-RRKK-K *(1-anthracenyl acetamido)*} segments, as shown by its sequencing (**Figure 3B**). The E6AP peptide, which interacts with the E6 protein and functions as a ubiquitin ligase, has an alpha-helical conformation according to the x-ray 16E6-MBP(E6AP)-P53 ternary complex [PDB ID: 4XR8]. To investigate whether the native core sequence of E6AP in Peptide-13 also adopts an alpha helix, we used the x-ray complex as a starting point, removed the MBP and P53 proteins, and kept the E6-bound E6AP peptide. We then conducted a 1.1 $\mu$s MD simulation in a box (60.0Å * 60.0Å * 60.0Å) of water (~6K water molecules) and ions and found that the E6AP peptide remained stable (**Supplementary Figure 10**), with an RMSD of 0.4 Å (measured using the C$\alpha$ atoms) compared to the E6AP peptide in the ternary complex.

Using the Molecular Operating Environment (MOE) software **(**Molecular Operating Environment (MOE), Canada**)**, we expanded upon the alpha helix E6AP core by incorporating the flexible N-terminal and C-terminal sequences to construct our Peptide-13 molecule (**Supplementary Figure 11B**). To optimize the Peptide-13 structure, we employed the LowModeMD algorithm**[4]**, which is integrated into MOE. We limited the sampling by imposing the following parameters defined in the MOE package: {Rejection Limit=100; Iteration Limit=10000; RMS Gradient=0.005, MM Iteration Limit=500; RMSD limit=0.75 Å; Energy Window =10.0 kcal/mol; Conformational Limit=10000}. Additionally, we treated the backbone atoms of the E6AP peptide as a rigid segment to maintain the alpha-helix conformation during the calculations.

To determine how Peptide-13 binds to the E6 protein (**Supplementary Figure 11C** and **11D**), we used the MOE docking module for molecular docking, which is known to be reliable for capturing peptide/receptor interactions[5,6]. To ensure the accuracy of our approach, we first performed molecular docking to determine the binding mode of the native E6AP peptide to the crystallographic E6 protein, using the X-ray E6-bound native E6AP peptide (PDB ID: 4XR8) as the starting point after eliminating the MBP and P53 from the ternary complex. We employed a docking protocol that started with the triangular matcher algorithm to quickly generate 1000 poses of docked peptides based on the receptor shape. From the resulting docked poses, we used the London dG scoring function to screen among the resulting poses to keep the top 100 structures for further refinement. We then used the rigid receptor replacement method combined with the GBVI/WSA dG scoring function to refine the resulting poses, obtaining an E6-bound E6AP peptide that closely matched the crystallographic binding mode of E6AP (**Supplementary Figure 12**). Similarly, we followed the same docking procedure to obtain top 3 docking poses of E6-Peptide-

13 complexes using our pre-generated conformations of peptide-13 (described above) to dock to the binding pocket of the native E6AP peptide in the E6.

Next, we performed ~60ns of constrained MD simulations to further refine the E6-Peptide-13 complexes in the presence of water and ions. Our goal was to push the peptide-13 warhead (Dha) toward the E6 Cys58 hotspot and create a covalent link between the two. To achieve this, we placed a harmonic restraint at 3.0 Å between Cys58 (S atom) and Dha (C atom), with a force constant of 0.36 kcal $mol^{-1}$ $Å^{-2}$. These simulations allowed us to prepare the E6-Peptide-13 constructs for the formation of the covalent link between Dha and Cys58.

Subsequently, we covalently linked Cys58 and Dha using MOE followed by energetically minimizing the covalent link and the E6-Pepitde-13 complexes. We then subjected each of three complexes to a ~120 ns classical MD simulation to characterize the binding mode of Peptide-13 covalently linked to the E6 protein. The resulting final complexes closely match the crystallographic E6AP binding peptide (**Supplementary Figure 13**). For further analysis, we selected the structure featuring the largest binding affinity between the peptide-13 core sequence and E6 while exhibiting the lowest RMSD of the peptide-13 core sequence compared to the crystallographic native E6AP peptide. For the results presented in **Figure 6**, we used **GROMOS** algorithm[7] with RMSD cutoff=0.18Å to cluster the entire MD trajectory of the best E6-Pepitde-13 complex.

#### *MD Simulation protocol*

Before running molecular docking and MD simulations, we used the MOE preparation module to refine the crystallographic E6 protein and added the missing sidechains/residues. The E6-pepitde complexes were then immersed in a water/ion (150 mM excess ions) box and underwent 500 steps of energy minimization using the steepest descents algorithm in GROMACS [8]. The refinement was followed by an MD simulation in a canonical ensemble, where the system was heated gradually from 0 K to 310 K in 20 ps, followed by MD simulations in an isobaric-isothermal ensemble for an aggregated 80 ps, during which the pressure was maintained at 1 bar to relax the simulation box. Throughout these pre-equilibration steps, the positional restraints were placed on all heavy atoms, gradually reduced to 0 kcal.$mol^{-1}Å^{2}$ for the final equilibration step. Finally, we optimized the E6-pepitde constructs by removing the positional restraints.

In all simulations, the AMBER-14 force field parameter set[9] was used to describe E6, Peptide-13, E6AP peptide, and ions. Non-canonical amino acid residues were parameterized with Antechamber program[10] to ensure compatibility with AMBER force field, while the force field parameters for two tagged molecules were borrowed from the Generalized AMBER force field (GAFF)[11] . The TIP3P model was utilized to describe water.

The temperature was maintained at 310 K using a velocity-rescale[12] thermostat with a damping constant of 1.0 ps for temperature coupling and the pressure was controlled at 1 bar using a Parrinello-Rahman barostat algorithm[13] with a 5.0 ps damping constant for the pressure coupling. Isotropic pressure coupling was used during this calculation. The Lennard-Jones cutoff radius was 12 Å, where the interaction was smoothly shifted to 0 after 10 Å. Periodic boundary conditions were applied to all three directions. The Particle Mesh Ewald algorithm [14] was used to calculate long-range coulombic interactions with a real cutoff radius of 10 Å and a grid spacing of 1.2 Å. A compressibility of $4.5 \times 10^{-5}$ $bar^{-1}$ was used to relax the box volume. In all the above simulations, water OH bonds were constrained by the SETTLE algorithm[15]. The remaining H-bonds were constrained using the P-LINCS algorithm[16]. All MD simulations were carried out using GROMACS, with constrained MD simulations aided by PULMED[17].

### 2. LC-MS characterization of peptides

#### 2.1 LC-MS methods

LC-MS characterizations were carried out using an Agilent 6550 quadrupole time-of-flight LC-MS. Total ion current (TIC) chromatograms were plotted. Mass spectra were integrated over the principal TIC peaks. High-performance liquid chromatography was done by the following methods: (solvent A: water with 0.1% formic acid; solvent B: MeCN with 0.1% formic acid).
**Method A:** Column: Phenomenex Aeris C4 column (1.0 × 150 mm, 5 μm particle size, 300 Å pore size) Gradient: 1% B (0-2 min), linear ramp from 1% B to 91% B (2-10 min). The flow rate is 100 μL/min. MS acquisition is from 2 to 10 min.
**Method B:** Column: Phenomenex Aeris C4 column (1.0 × 150 mm, 5 μm particle size, 300 Å pore size) Gradient: 1% B (0-2 min), linear ramp from 1% B to 91% B (2-8 min), 61% B to 95% B (8-10 min). The flow rate is 100 μL/min. MS acquisition is from 2 to 8 min.
**Method C:** Column: Agilent Zorbax 300SB C3 column (2.1 × 150 mm, 5 μm particle size, 300 Å pore size) Gradient: 1% B (0-2 min), linear ramp from 1% B to 91% B (2-12 min), 91% B to 91% B (12-13 min). The flow rate is 500 μL/min. MS acquisition is from 4 to 12 min.
**Method D:**
Column: Phenomenex Jupiter C4 column (1.0 × 150 mm, 5 μm particle size, 300 Å pore size) Gradient: 1% B (0-2 min), linear ramp from 1% B to 91% B (2-18 min), 91% B to 91% B (18-21 min). The flow rate is 100 μL/min. MS acquisition is from 4 to 18 min.
**Method E:**
Column: Agilent Zorbax 300SB C3 column (2.1 × 150 mm, 5 μm particle size, 300 Å pore size) Gradient: 1% B (0-1 min), linear ramp from 1% B to 91% B (1-11 min), 91% B to 91% B (11-15 min). The flow rate is 500 μL/min. MS acquisition is from 0 to 11 min.

#### 2.2 Peptides 1-13

Name: **Peptide 1**
Sequence**:** IPESSELTLQELLGEERR-$NH_2$
HPLC: method A
HRMS (ESI-QTOF): Calcd. for (M +2H)2+: 2097.10, found: 2097.15.
**Total ion chromatogram:**

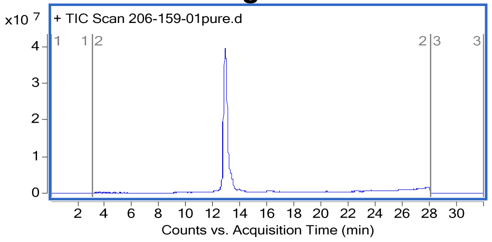


**Mass spectrum of major compound:**

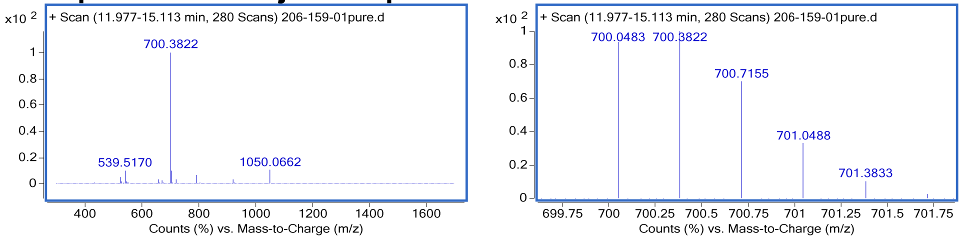

Name: Peptide **1-biotin**

Sequence**:** Biotin-(PEG)4-IPESSELTLQELLGEERR

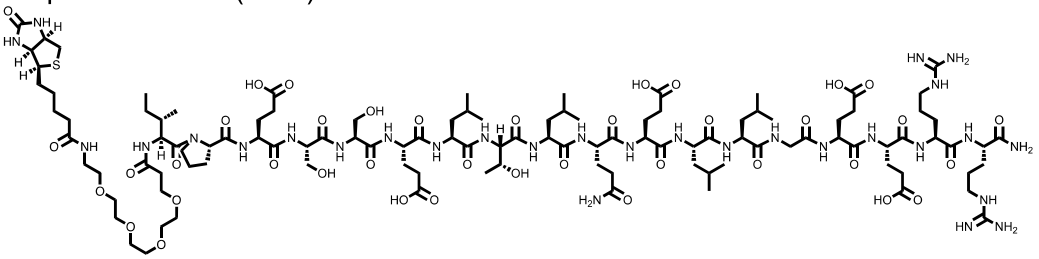

HPLC: method A

HRMS (ESI-QTOF): Calcd. for (M +2H)2+: 2570.65, found: 2570.64.

**Total ion chromatogram:**

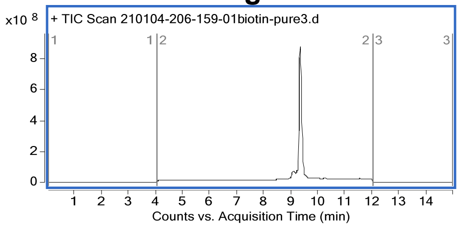


**Mass spectrum of major compound:**

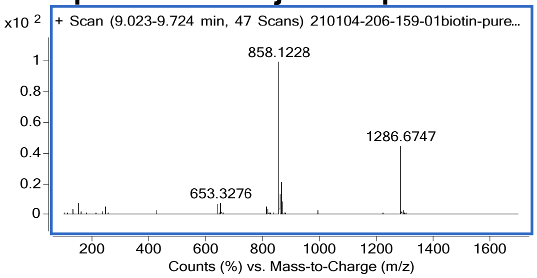


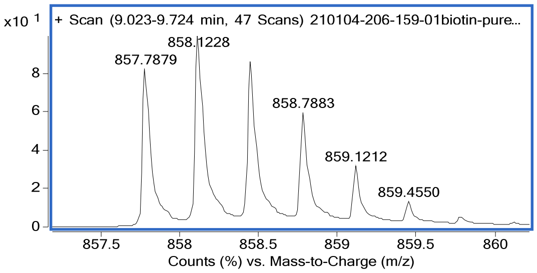


Name: **Peptide 2**

Sequence**:**

| N-terminal | Sequence | C-terminal |
|---|---|---|
| 9-fluorene acetamido-PEG1 | IPESSELTLQELLGEERRAA | Lys(9-fluorene acetamido) |

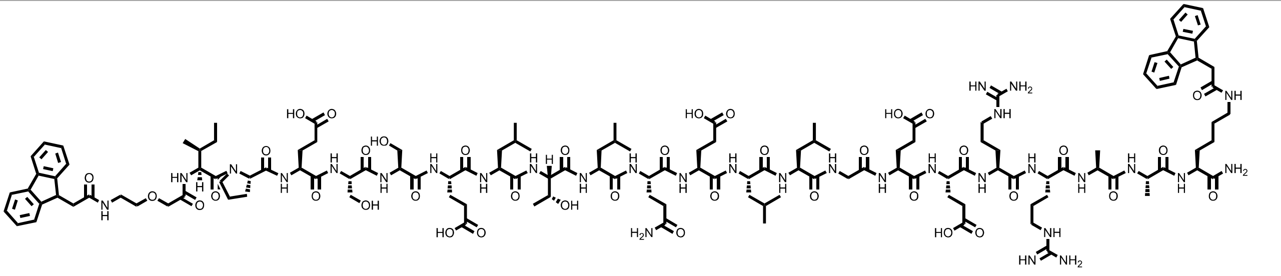

HPLC: method A

HRMS (ESI-QTOF): Calcd. for $(M + H)^+$: 2880.50, found: 2880.50.

**Total ion chromatogram:**

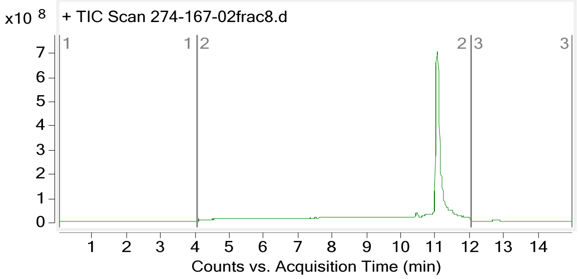


**Mass spectrum of major compound:**

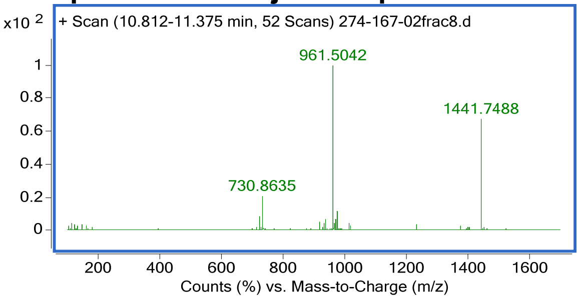


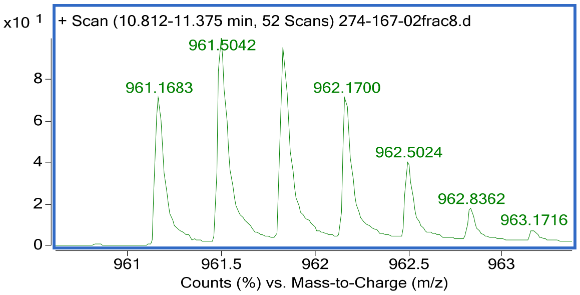


Name: **Peptide 3**
Sequence**:**

| N-terminal | Sequence | C-terminal |
|---|---|---|
| 9-fluorene acetamido | IPESSELTLQELLGEERRAA | Lys(5-Acenaphthyl carboxamido) |

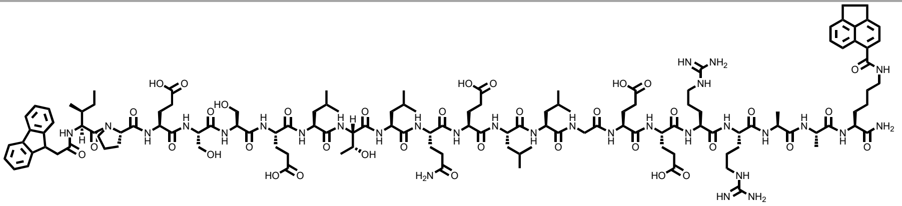

HPLC: method A
HRMS (ESI-QTOF): Calcd. for $(M + H)^+$: 2760.46, found: 2760.46

**Total ion chromatogram:**

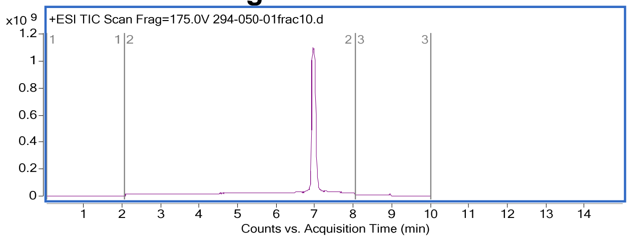


**Mass spectrum of major compound:**

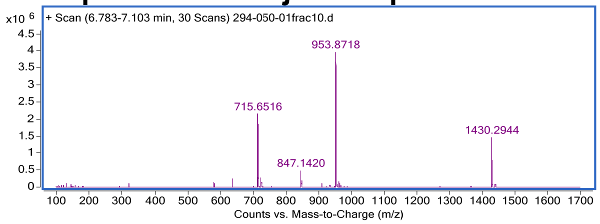


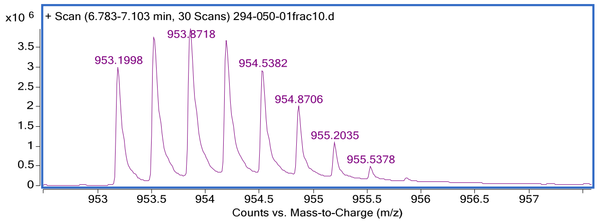

Name: **Peptide 4**
Sequence**:**

| N-terminal | Sequence | C-terminal |
|---|---|---|
| 9-fluorene acetamido | IPESSELTLQELLGEERRAA | Lys(1-naphthylcarboxamido) |

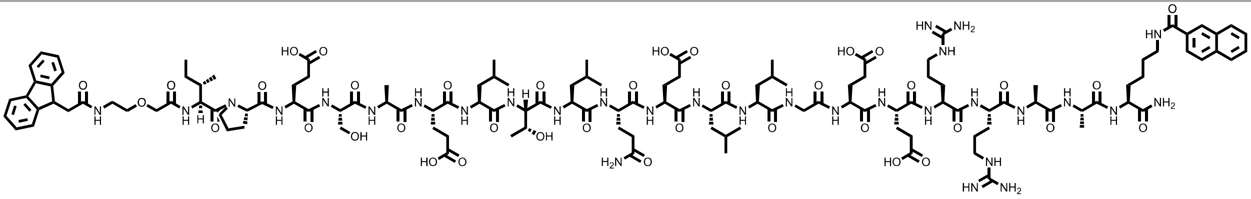

HPLC: method A
HRMS (ESI-QTOF): Calcd. for $(M + 3H)^{3+}$: 938.49, found:938.49.

**Total ion chromatogram:**

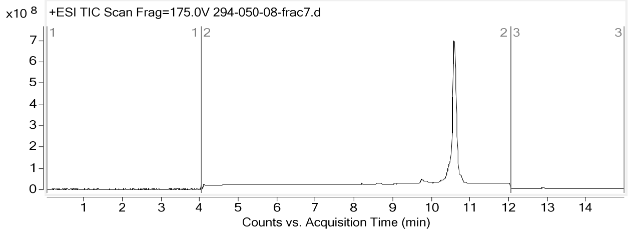


**Mass spectrum of major compound:**

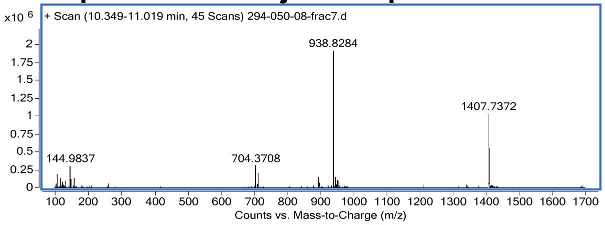


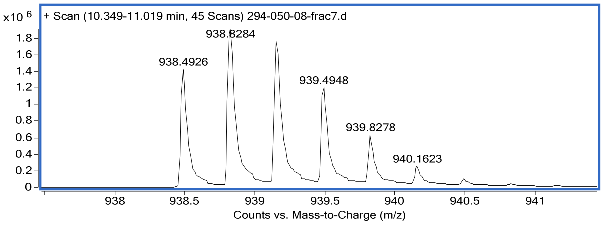


Name: **peptide 5**
Sequence**:**

| N-terminal | Sequence | C-terminal |
|---|---|---|
| 9-fluorene acetamido-PEG$_1$ | IPESSELTLQELLGEERRAA | Lys(anthraquinone-2-carboxamido) |

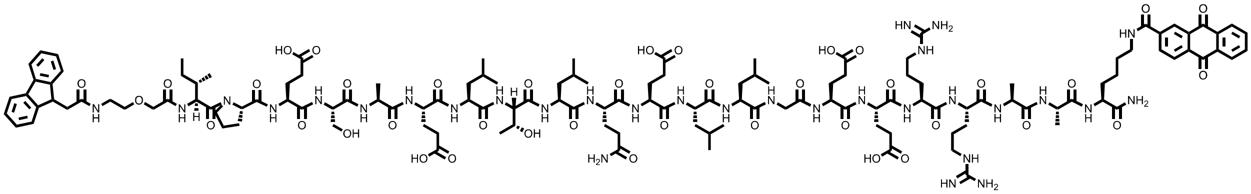

HPLC: method A
HRMS (ESI-QTOF): Calcd. for $(M + 2H)^{2+}$: 1447.18, found: 1447.18

### Total ion chromatogram:

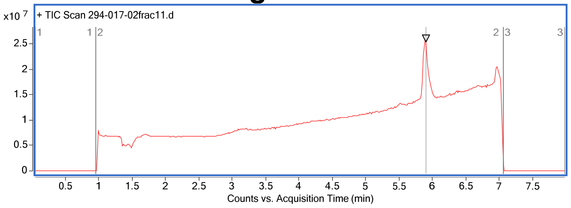


### Mass spectrum of major compound:

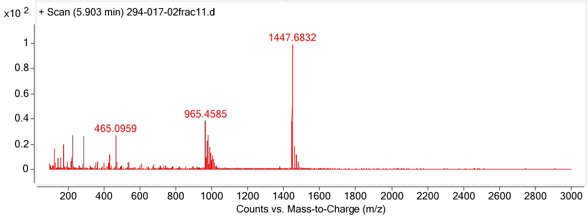


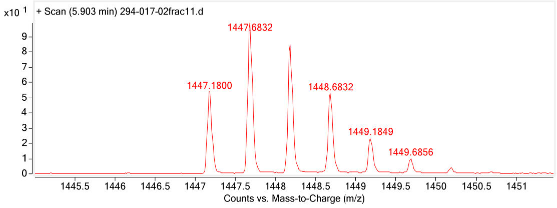


Name: **6’-biotin**
Structure**:**

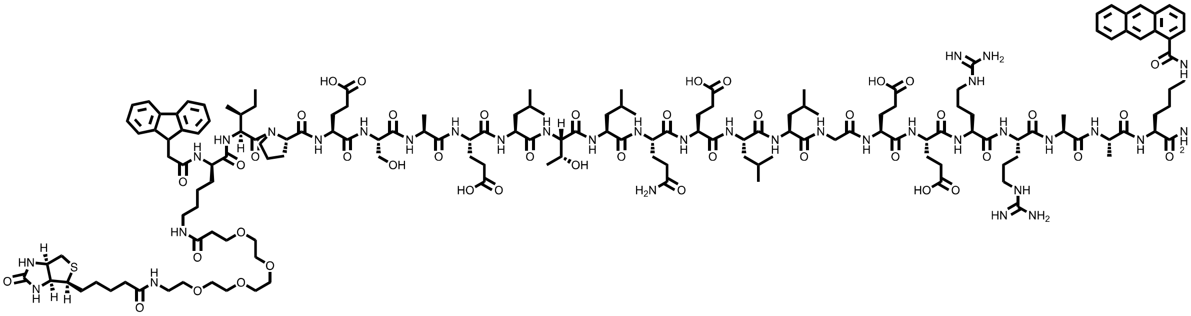

HPLC: method A
HRMS (ESI-QTOF): Calcd. for $(M + 3H)^{3+}$: 1121.92, found: 1121.92

### Total ion chromatogram:

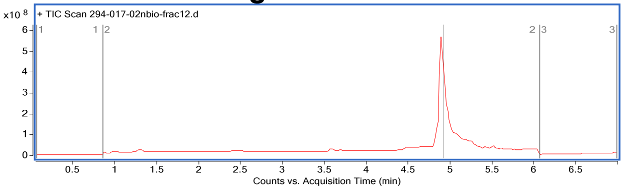


### Mass spectrum of major compound:

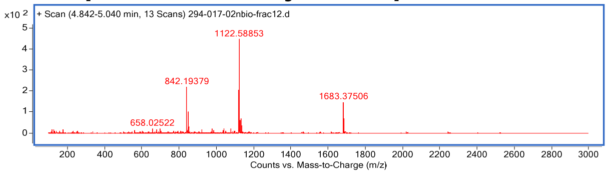


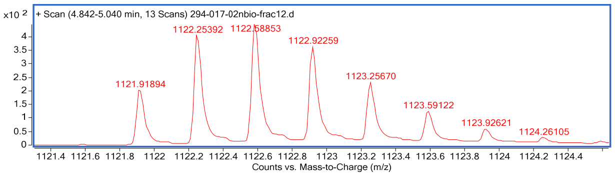

Name: **peptide 6’**
Sequence**:**

| N-terminal | Sequence | C-terminal |
|---|---|---|
| 9-fluorene acetamido-PEG$_1$ | IPESAELTLQELLGEERRAA | Lys(1-anthracene carboxamido) |

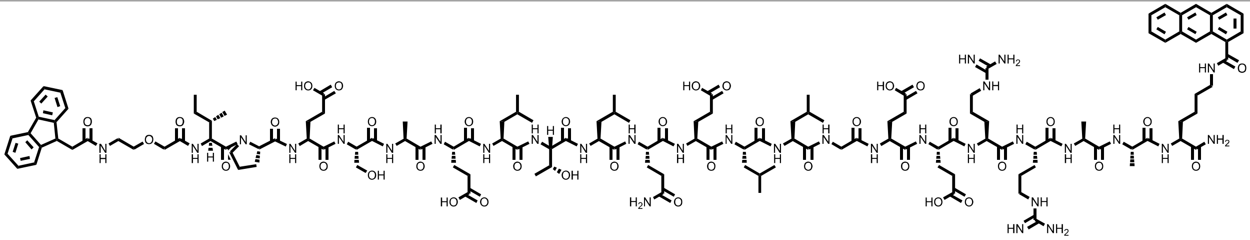

HPLC: method A
HRMS (ESI-QTOF): Calcd. for $(M + 2H)^{2+}$: 1432.20, found: 1432.21.

**Total ion chromatogram:**

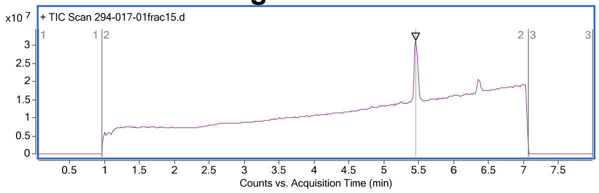


**Mass spectrum of major compound:**

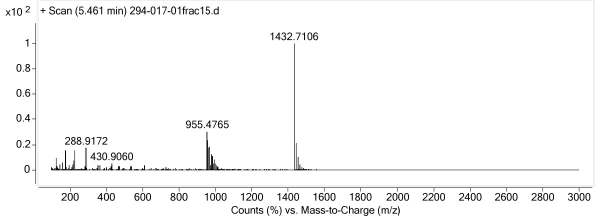


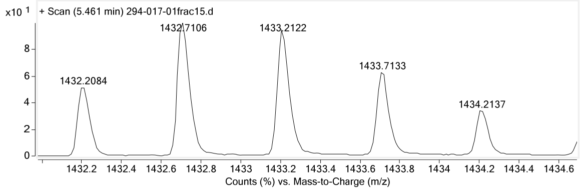


Name: **peptide 6**
Sequence**:**

| N-terminal | Sequence | C-terminal |
|---|---|---|
| 9-fluorene acetamido-PEG$_1$ | IPES**S**ELTLQELLGEERRAA | Lys(1-anthracene carboxamido) |

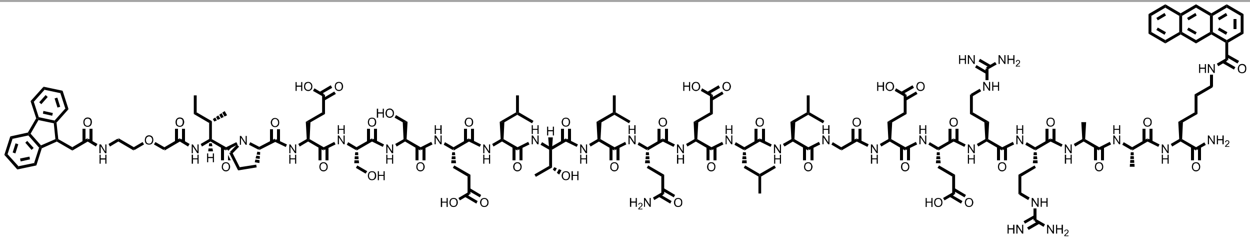

HPLC: method A
HRMS (ESI-QTOF): Calcd. for $(M + 3H)^{3+}$: 960.50, found: 960.497,

### Total ion chromatogram:

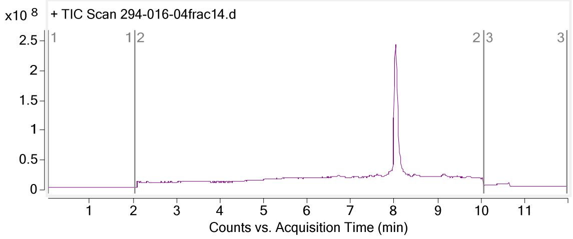


### Mass spectrum of major compound:

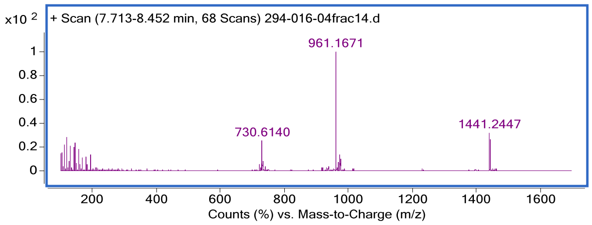


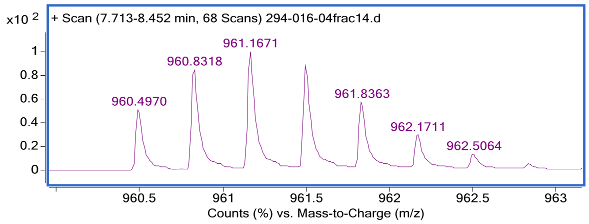


Name: **peptide 6’-3L3A**
Sequence**:**

| N-terminal | Sequence | C-terminal |
|---|---|---|
| 9-fluorene acetamido-PEG$_1$ | IPESAELTAQEAAGEERRAA | Lys(1-anthracene carboxamido) |

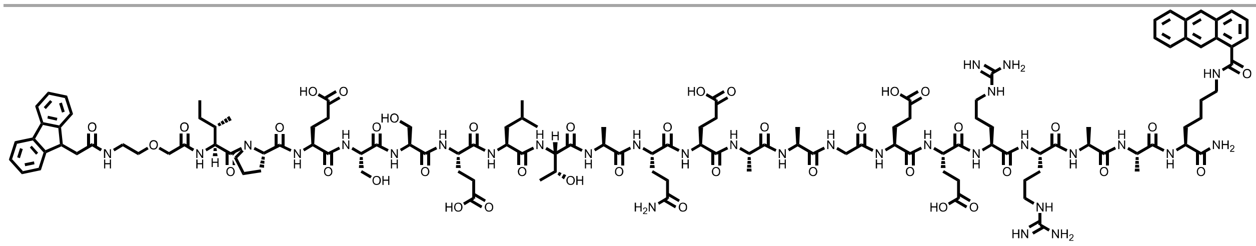

HRMS (ESI-QTOF): Calcd. for (M + H)$^+$: 2754.3548, found: 2754.3548.

### Total ion chromatogram:

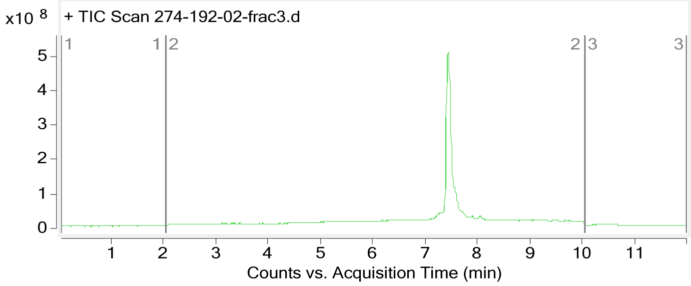


### Mass spectrum of major compound:

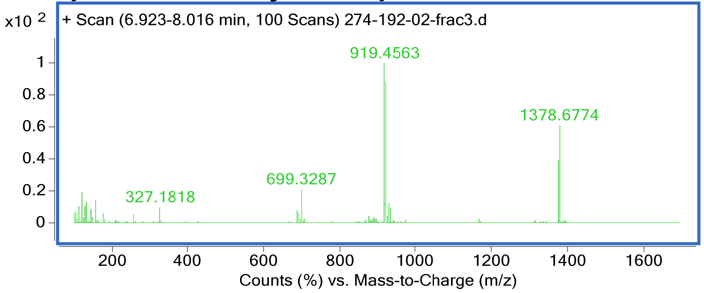

Name: **Peptide 7**
Sequence**:**

| N-terminal | Sequence | C-terminal |
|---|---|---|
| 9-fluorene acetamido | IPESAELTLQELL(DHA)EERRAA | Lys(1-anthracene carboxamido) |

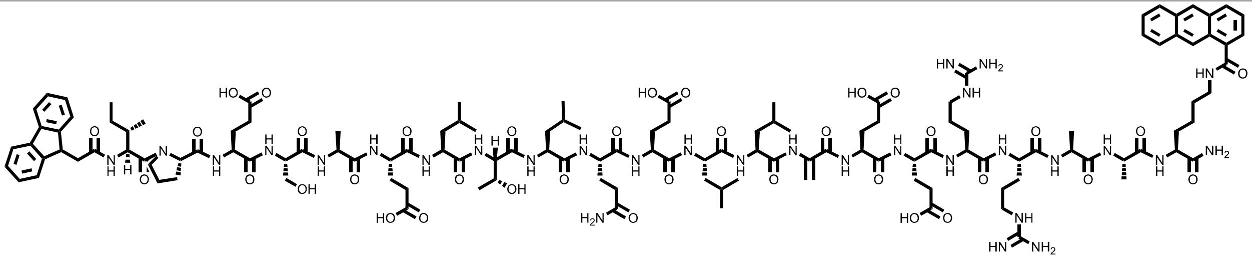

HPLC: method A
HRMS (ESI-QTOF): Calcd. for $(M + 2H)^{2+}$: 1387.73, found: 1387.73

**Total ion chromatogram:**

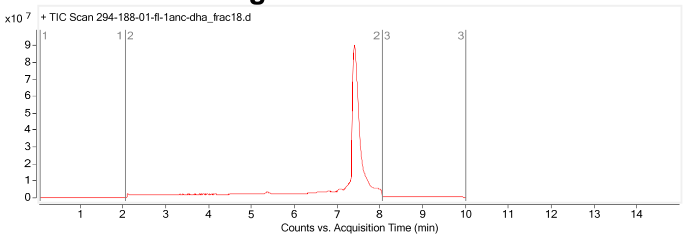


**Mass spectrum of major compound:**

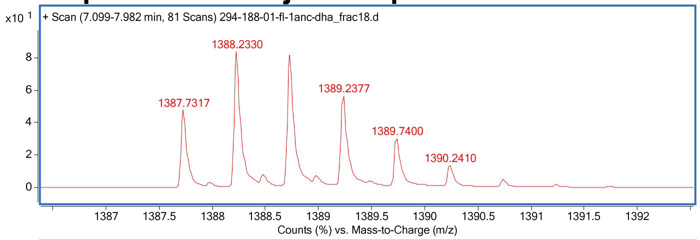


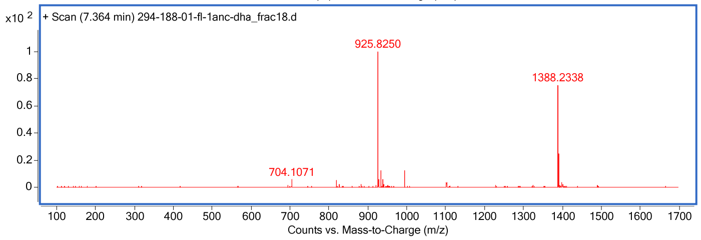

Name: **peptide 8**
Structure:

| N-terminal | Sequence | C-terminal |
|---|---|---|
| 9-fluorene acetamido | IPESSELTLQELL(DHA)EERRNK | Lys(1-anthracene carboxamido) |

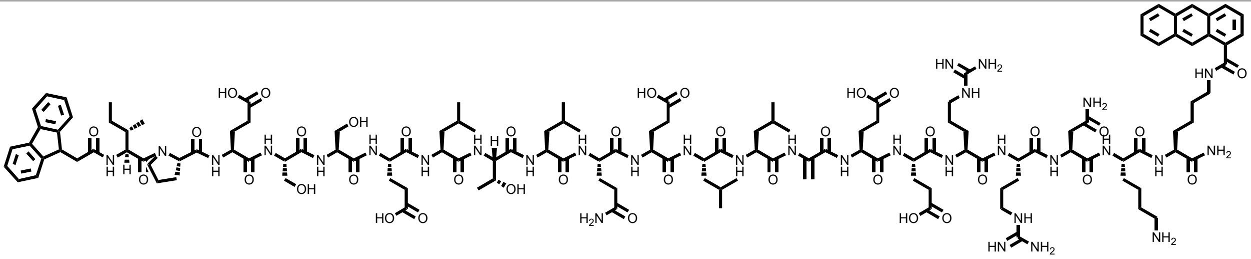

HPLC: method A
HRMS (ESI-QTOF): Calcd. for $(M + 3H)^{3+}$:962.20, found: 962.20.

**Total ion chromatogram:**

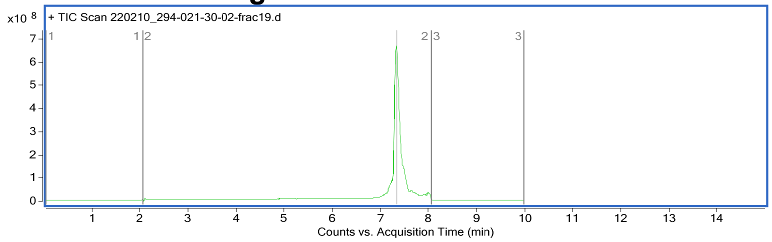


**Mass spectrum of major compound:**

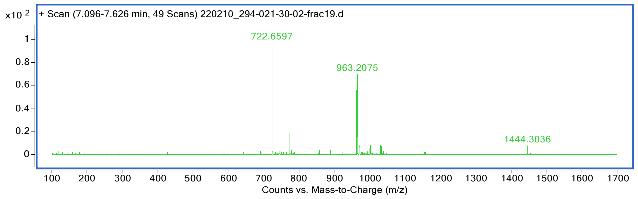


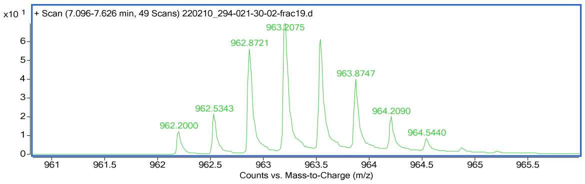


Name: **Peptide 9**
Sequence**:**

| N-terminal | Sequence | C-terminal |
|---|---|---|
| 9-fluorene acetamido | IPQSAELTLQELL(DHA)EARRNKK | Lys(1-anthracene carboxamido) |

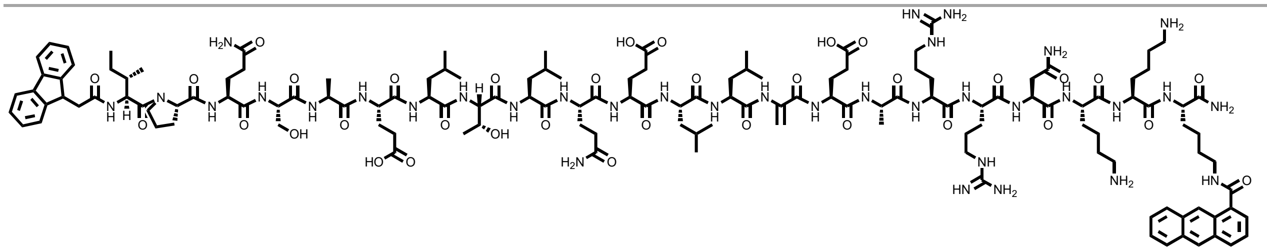

HPLC: method A
HRMS (ESI-QTOF): Calcd. for $(M + 4H)^{4+}$: 705.65, found: 705.65.

**Total ion chromatogram:**

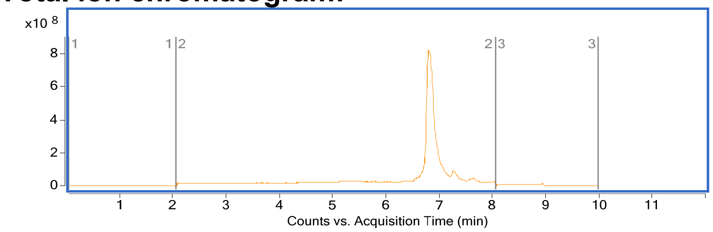


**Mass spectrum of major compound:**

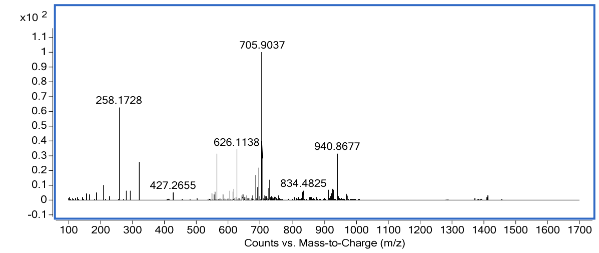


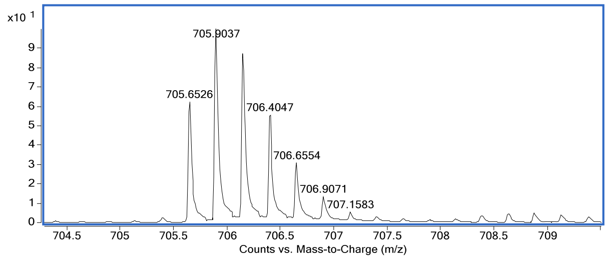


Name: **Peptide 10**
Sequence**:**

| N-terminal | Sequence | C-terminal |
|---|---|---|
| 9-fluorene acetamido | IPQSAELTLQELL(DHA)QARRNKK | Lys(1-anthracene carboxamido) |

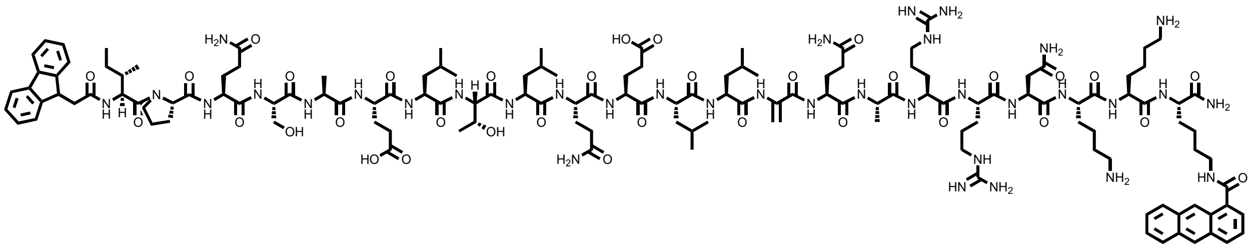

HPLC: method A
HRMS (ESI-QTOF): Calcd. for $(M + 2H)^{2+}$: 1472.89, found: 1472.78.

**Total ion chromatogram:**

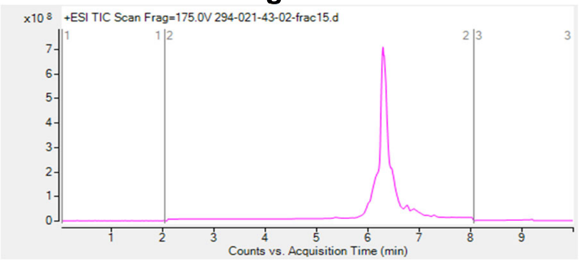


**Mass spectrum of major compound:**

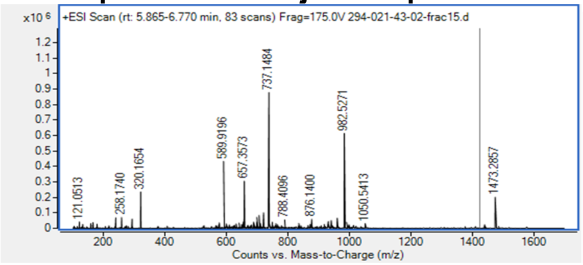


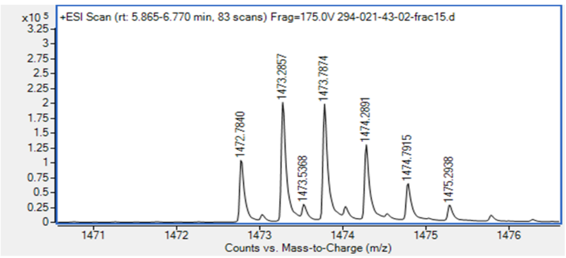

Name: **peptide 11**
Sequence**:**

| N-terminal | Sequence | C-terminal |
|---|---|---|
| 9-fluorene acetamido | IPQSAELTLQELL(DHA)QARRKK | Lys(1-anthracene carboxamido) |

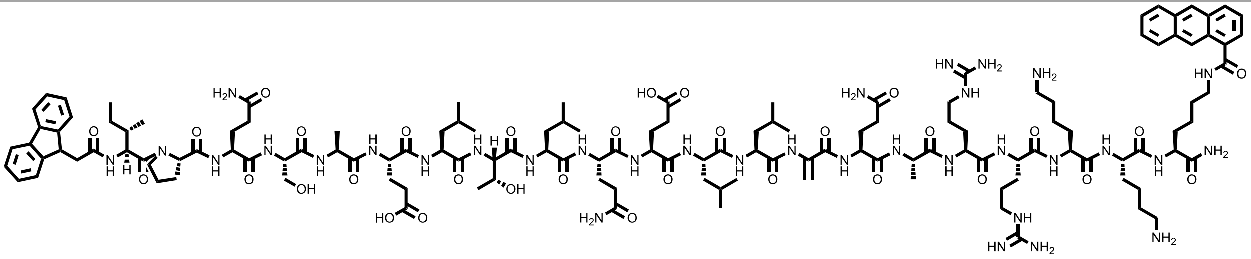

HPLC: method A
HRMS (ESI-QTOF): Calcd. for $(M + 3H)^{3+}$: 943.52, found: 943.52.

**Total ion chromatogram:**

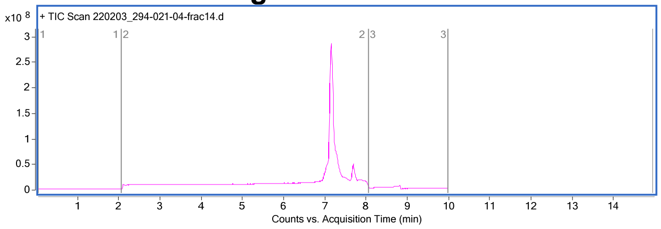


**Mass spectrum of major compound:**

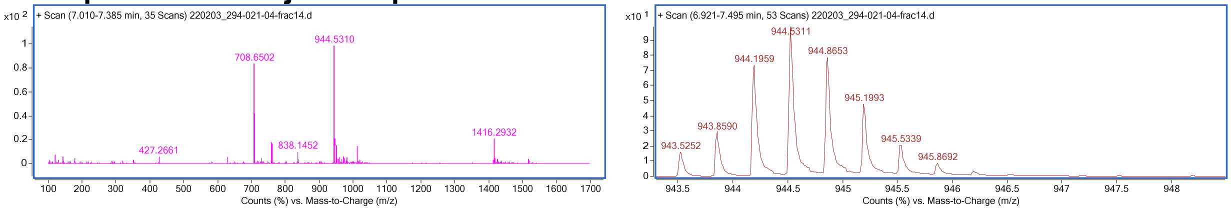


Name: **peptide 12**
Sequence**:**

| N-terminal | Sequence | C-terminal |
|---|---|---|
| 9-fluorene acetamido | IPQSAELTLQELL(DHA)QRRKK | Lys(1-anthracene carboxamido) |

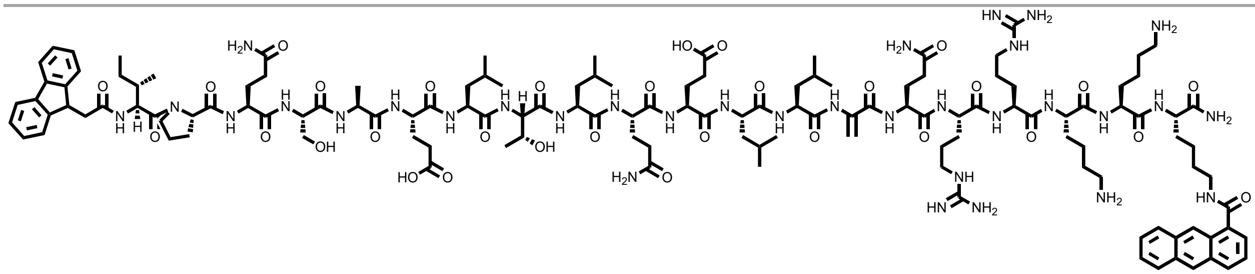

HPLC: method A
HRMS (ESI-QTOF): Calcd. for $(M + 3H)^{3+}$: 919.87, found: 919.85.

### Total ion chromatogram:

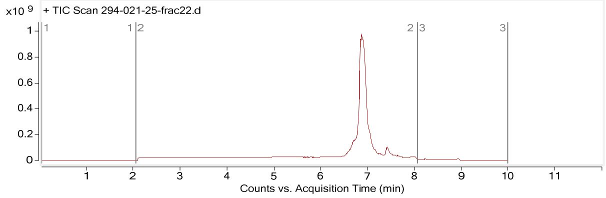


### Mass spectrum of major compound:

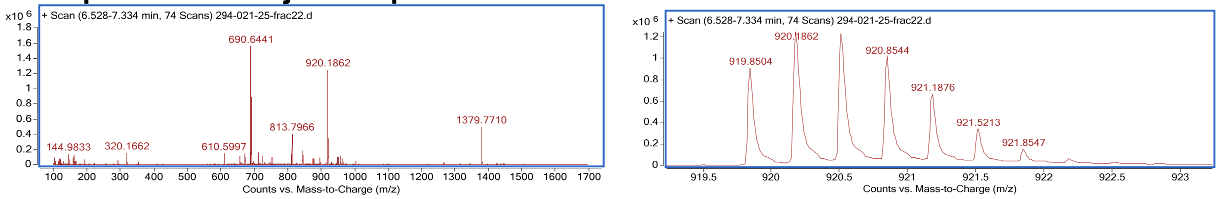


Name: **peptide 13**
Sequence**:**

| N-terminal | Sequence | C-terminal |
|---|---|---|
| 9-fluorene acetamido | IPQSAELTLQELL(DHA)RRKK | Lys(1-anthracene carboxamido) |

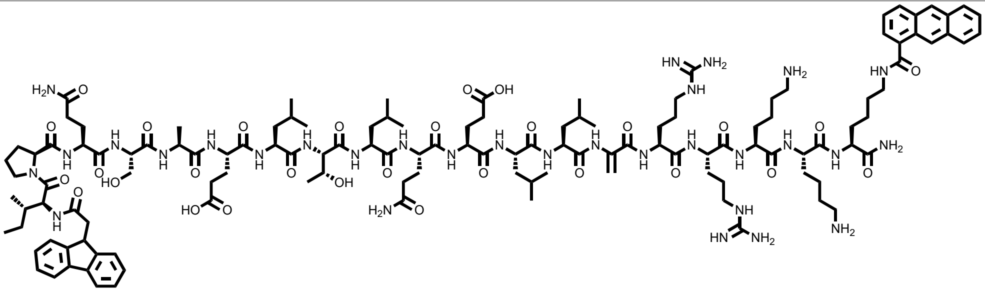

HPLC: method A
HRMS (ESI-QTOF): Calcd. for $(M + 2H)^{2+}$: 1315.23, found: 1315.24.

### Total ion chromatogram:

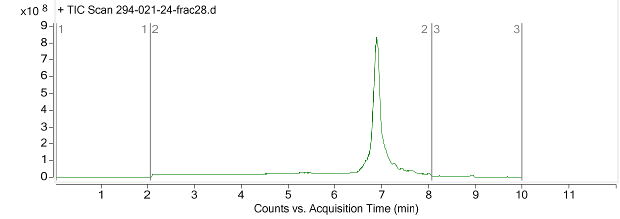


### Mass spectrum of major compound:

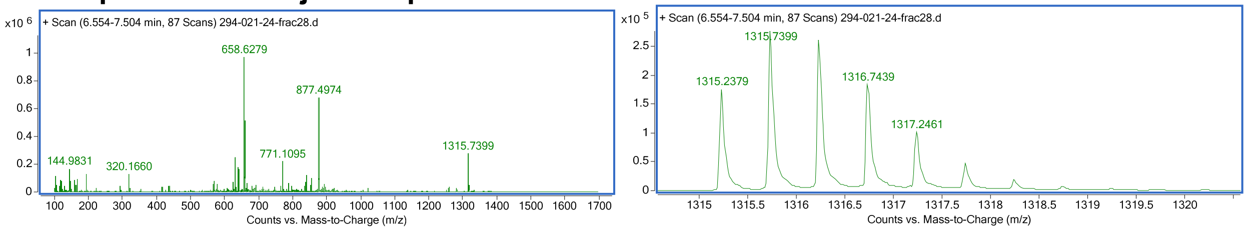

Name **peptide 13-3L3A**
Sequence**:**

| N-terminal | Sequence | C-terminal |
|---|---|---|
| 9-fluorene acetamido | IPQSAELTAQEAA(DHA)RRKK | Lys(1-anthracene carboxamido) |

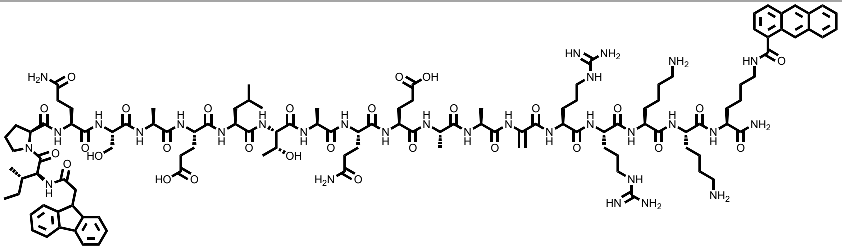

HPLC: method A
HRMS (ESI-QTOF): Calcd. for $(M + H)^+$: 2502.32, found:2502.32.

**Total ion chromatogram:**

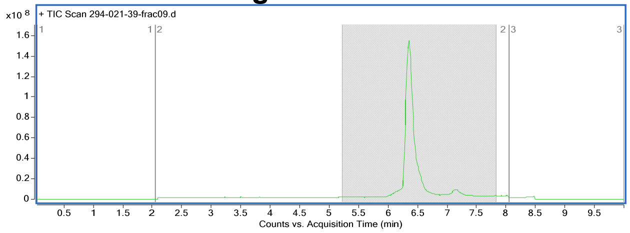


**Mass spectrum of major compound:**

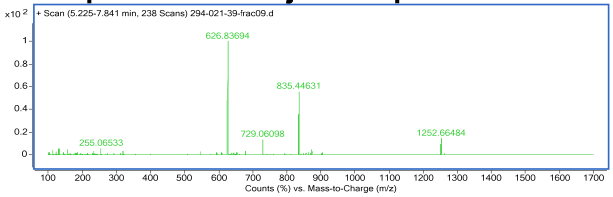


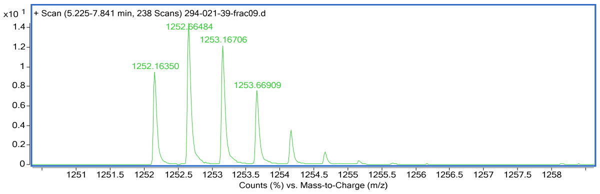


Name: **peptide 13-biotin**
Structure**:**

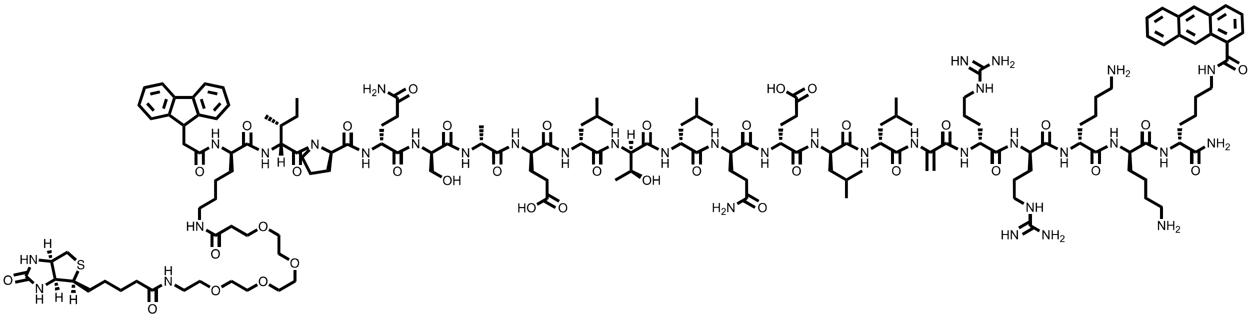

HPLC: method A
HRMS (ESI-QTOF): Calcd. for $(M + H)^+$: 3229.77, found: 3229.78
**Total ion chromatogram:**

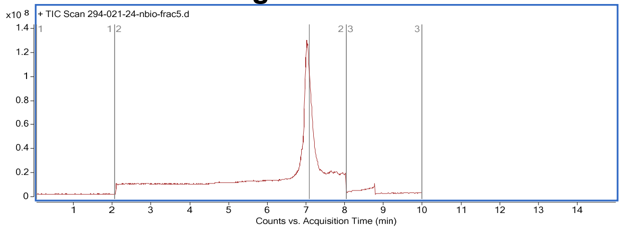

**Mass spectrum of major compound:**

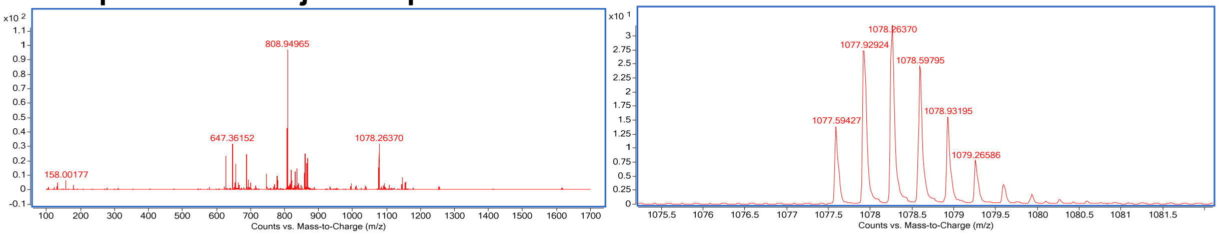


Name **peptide 13-3L3A-biotin**

Structure**:**

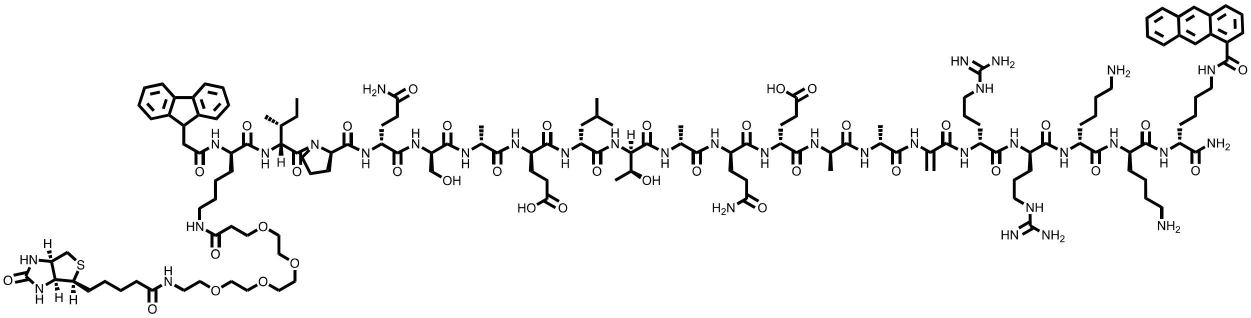

HPLC: method A

HRMS (ESI-QTOF): Calcd. for $(M + H)^{+}$: 3103.63, found: 3103.69.

**Total ion chromatogram:**

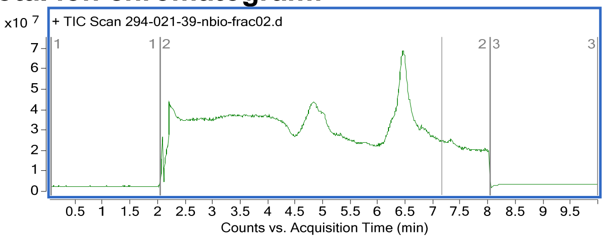


**Mass spectrum of major compound:**

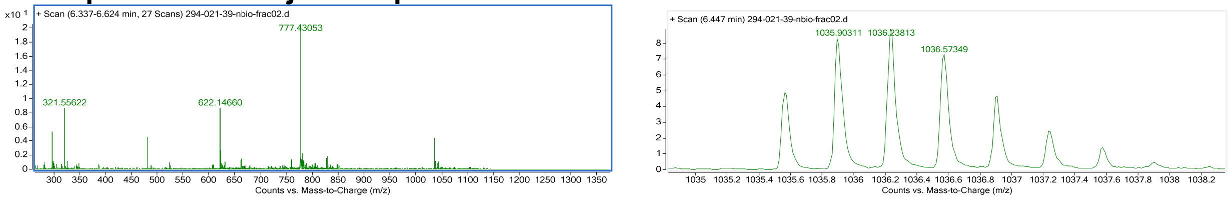


Name: **peptide 6'-TAMRA**

Structure**:**

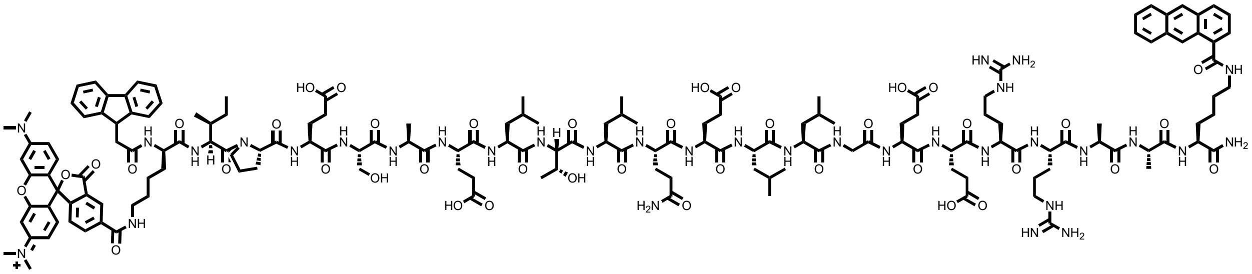

HPLC: method A

HRMS (ESI-QTOF): Calcd. for $(M + H)^{+}$: 3301.67, found: 3301.72.

**Total ion chromatogram:**

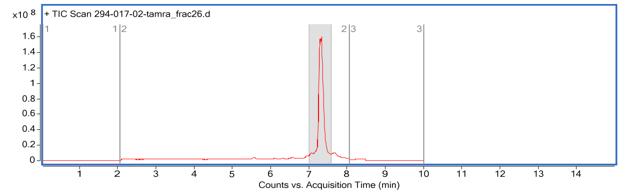


### Mass spectrum of major compound:

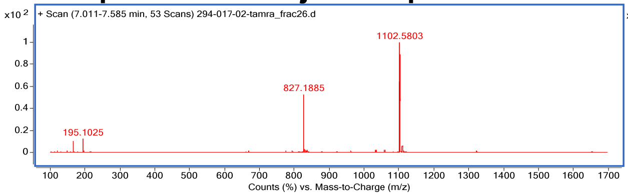


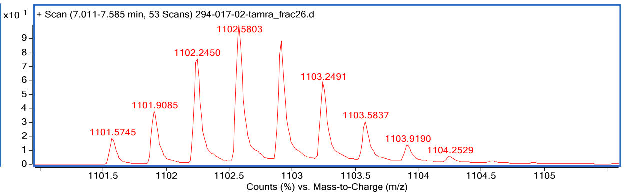


Name: **peptide 13-TAMRA**

Structure**:**

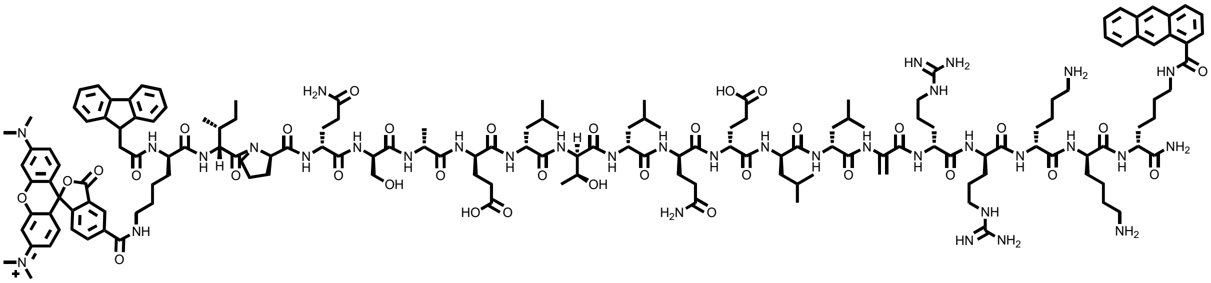

HPLC: method A

HRMS (ESI-QTOF): Calcd. for $(M + H)^+$: 3168.70, found: 3168.73

### Total ion chromatogram:

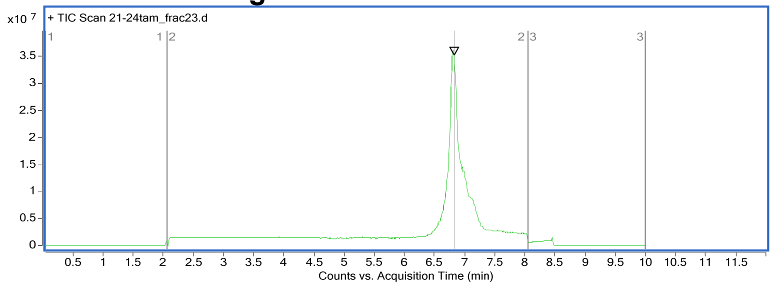


### Mass spectrum of major compound:

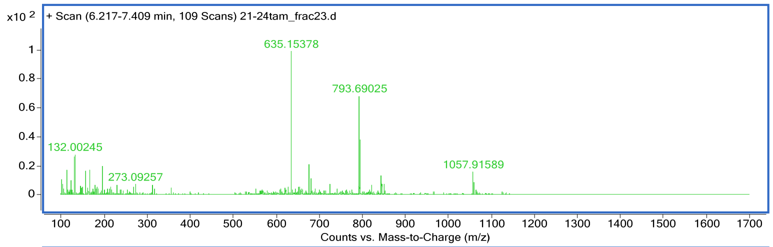


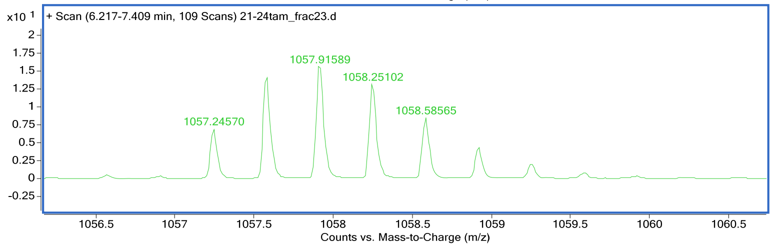

### 2.3 Peptide N1-N22 and N-FITC

Name: peptide **N1**

HPLC: method A

HRMS (ESI-QTOF): Calcd. for (M +2H)2+: 1153.53, found: 1153.5297.

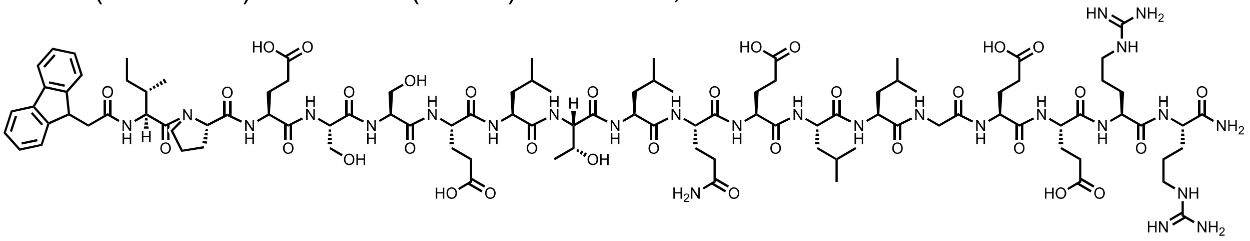

**Total ion chromatogram:**

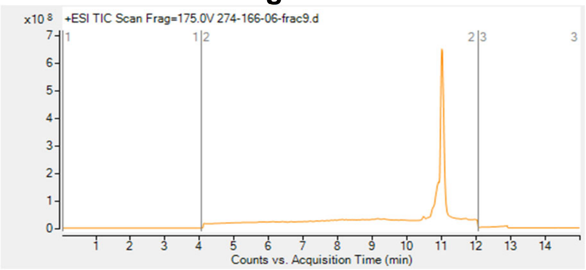


**Mass spectrum of major compound:**

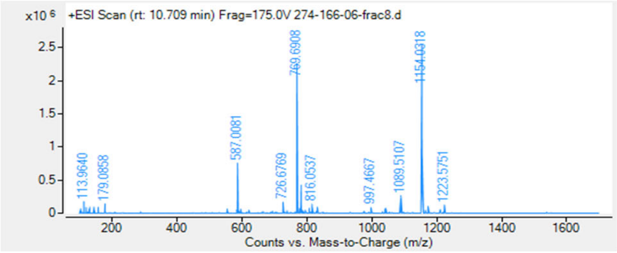


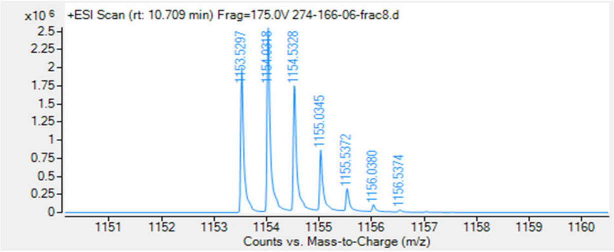


Name: peptide **N2**

HPLC: method A

HRMS (ESI-QTOF): Calcd. for (M +2H)2+: 1121.57, found: 1121.59.

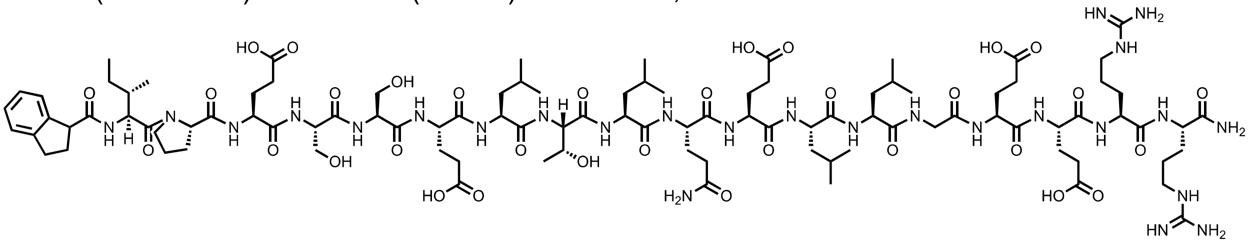

**Total ion chromatogram:**

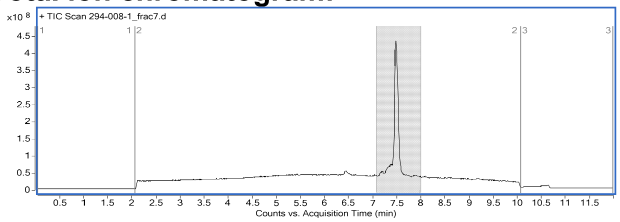


**Mass spectrum of major compound:**

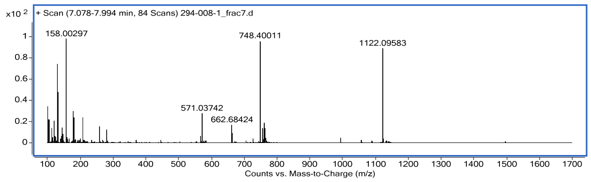


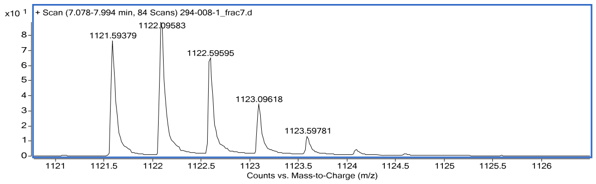


Name: peptide **N3**
HPLC: method A
HRMS (ESI-QTOF): Calcd. for (M +2H)2+:1145.57, found: 1145.57.

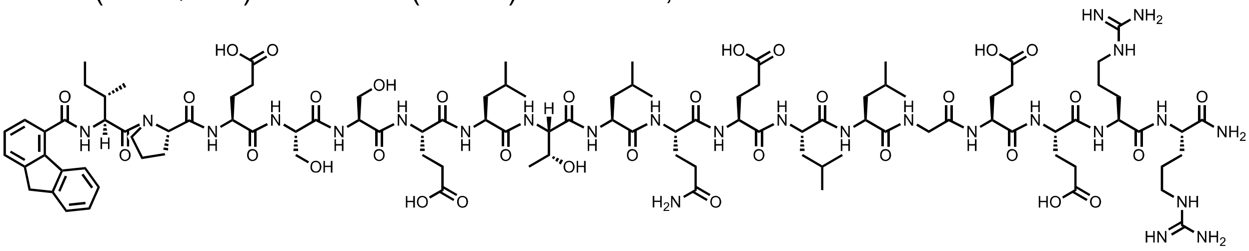

**Total ion chromatogram:**

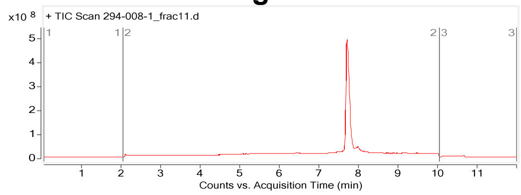


**Mass spectrum of major compound:**

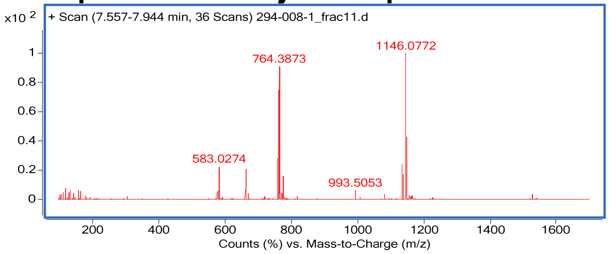


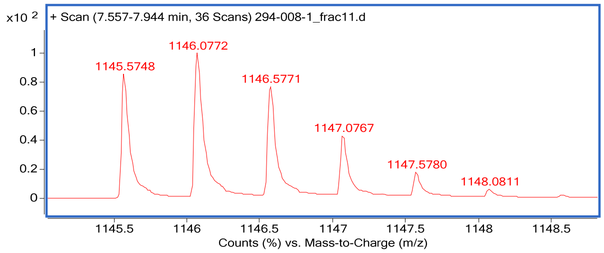


Name: peptide **N4**
HPLC: method A
HRMS (ESI-QTOF): Calcd. for (M +2H)2+: 1152.58, found: 1152.58.

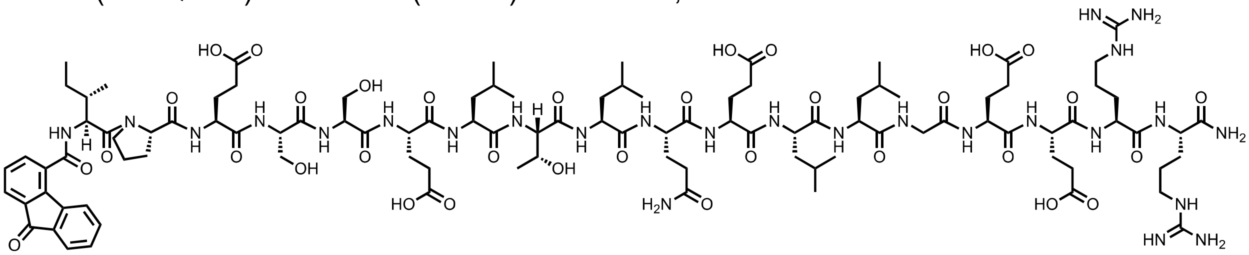

**Total ion chromatogram:**

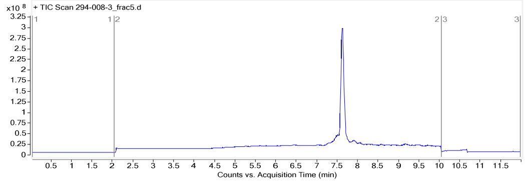


**Mass spectrum of major compound:**

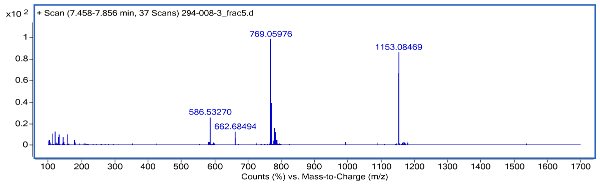


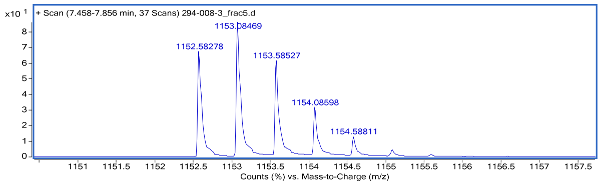


Name: peptide **N5**
HPLC: method A
HRMS (ESI-QTOF): Calcd. for (M +2H)2+: 1153.59, found: 1153.59

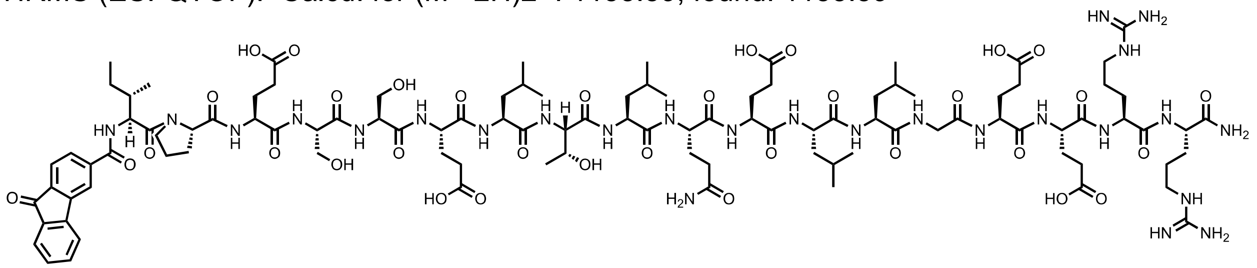

**Total ion chromatogram:**

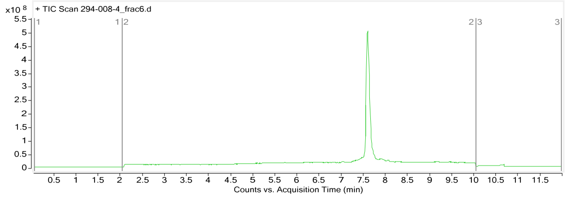


**Mass spectrum of major compound:**

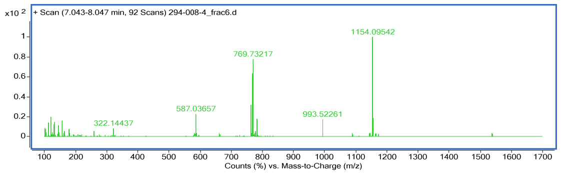


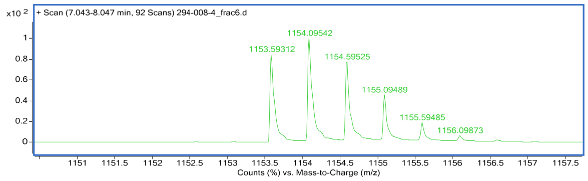


Name: peptide **N6**
HPLC: method A
HRMS (ESI-QTOF): Calcd. for (M +2H)2+: 1152.57, found: 1152.58.

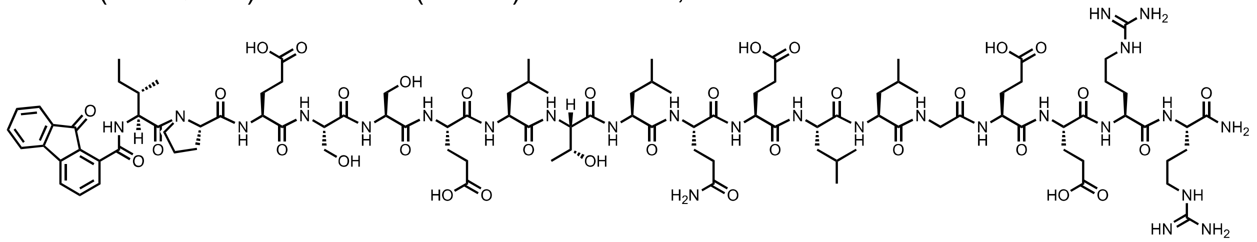

**Total ion chromatogram:**

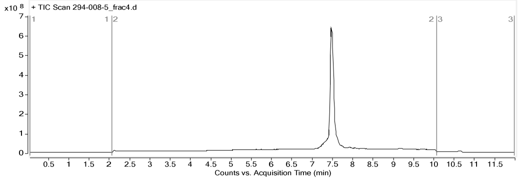


**Mass spectrum of major compound:**

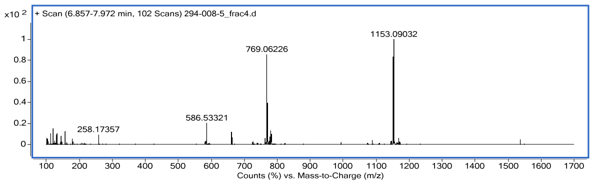


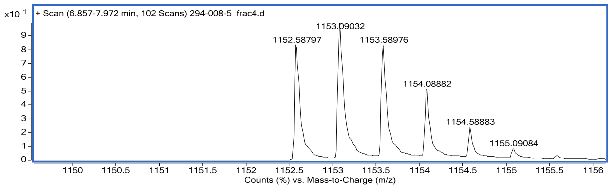


Name: peptide **N7**
HPLC: method A
HRMS (ESI-QTOF): Calcd. for $(M + 2H)^{2+}$: 1167.086, found: 1167.08.

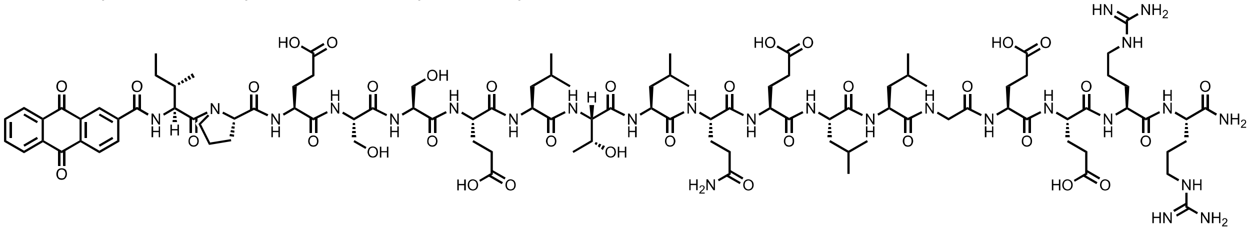

**Total ion chromatogram:**

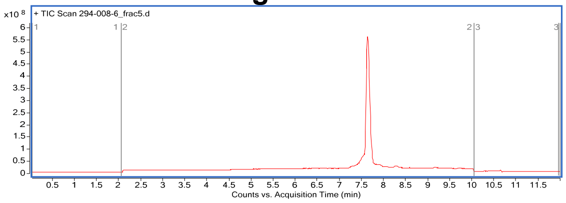


**Mass spectrum of major compound:**

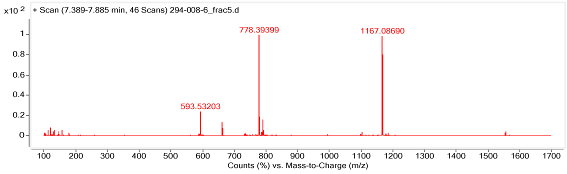


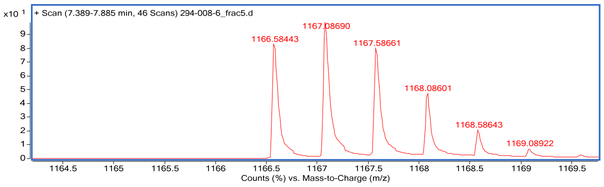


Name: peptide **N8**
HPLC: method A
HRMS (ESI-QTOF): Calcd. for $(M + 2H)^{2+}$: 1153.59, found: 1153.59.

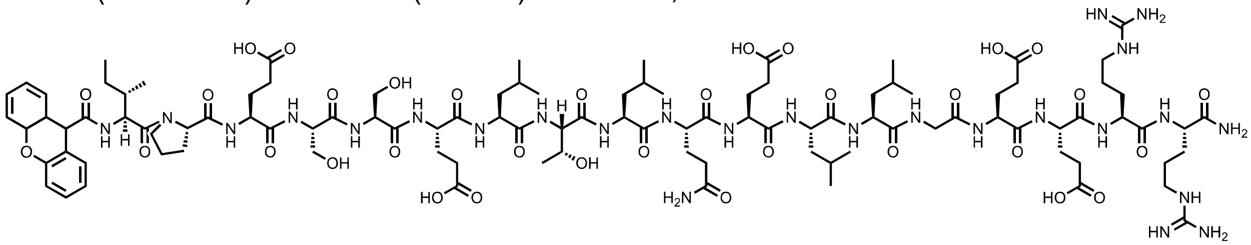

**Total ion chromatogram:**

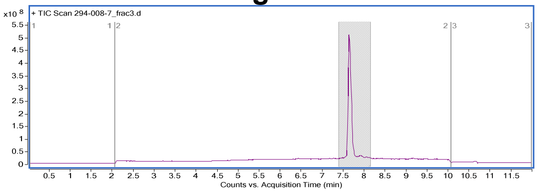


**Mass spectrum of major compound:**

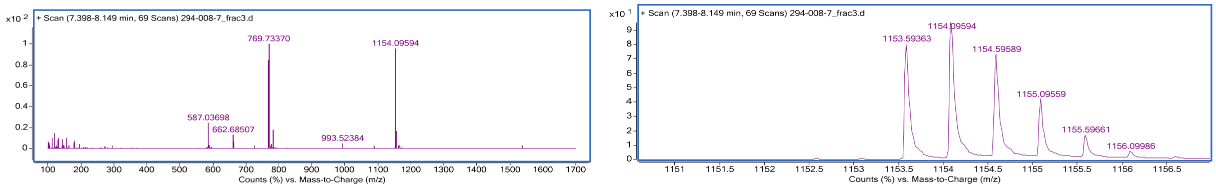


Name: peptide **N9**
HPLC: method A
HRMS (ESI-QTOF): Calcd. for (M +2H)2+: 1152.10, found: 1152.10.

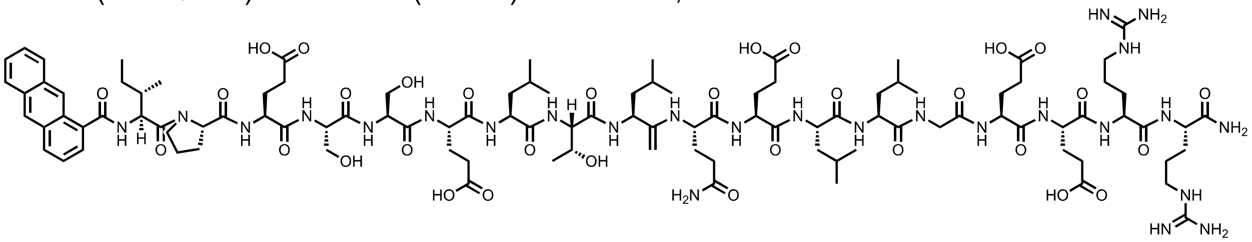

### Total ion chromatogram:

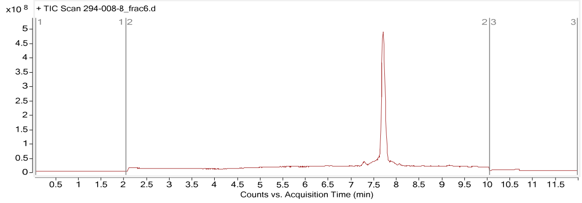


### Mass spectrum of major compound:

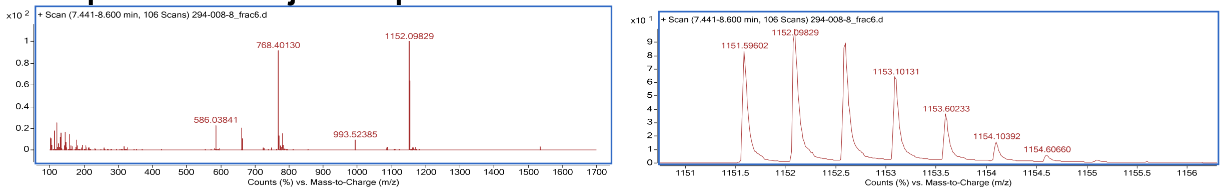


Name: peptide **N10**
HPLC: method A
HRMS (ESI-QTOF): Calcd. for $(M + H)^+$: 2301.19, found: 2301.19

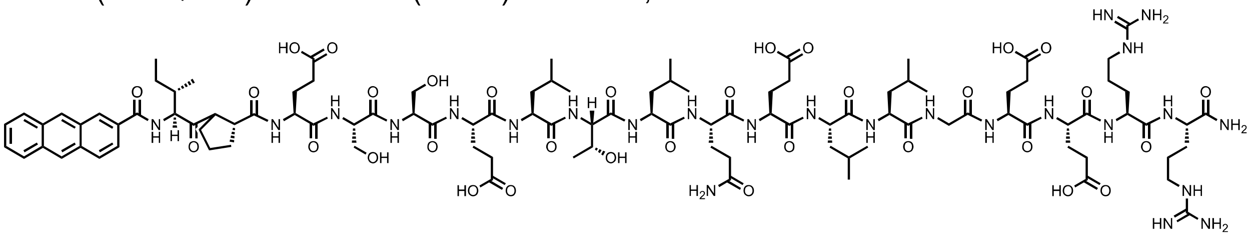

### Total ion chromatogram:

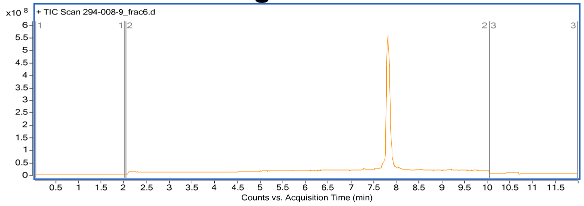


### Mass spectrum of major compound:

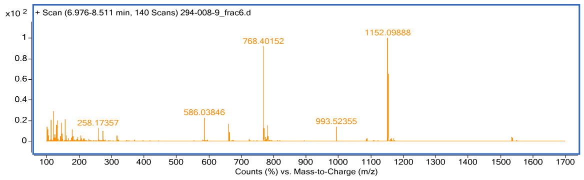


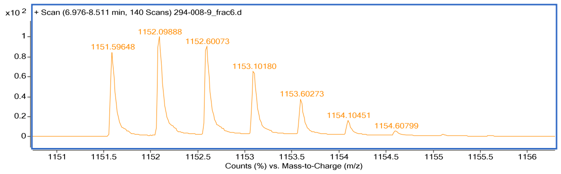


Name: peptide **N11**
HPLC: method A
HRMS (ESI-QTOF): Calcd. for $(M + 2H)^{2+}$: 1131.12 , found: 1131.12.

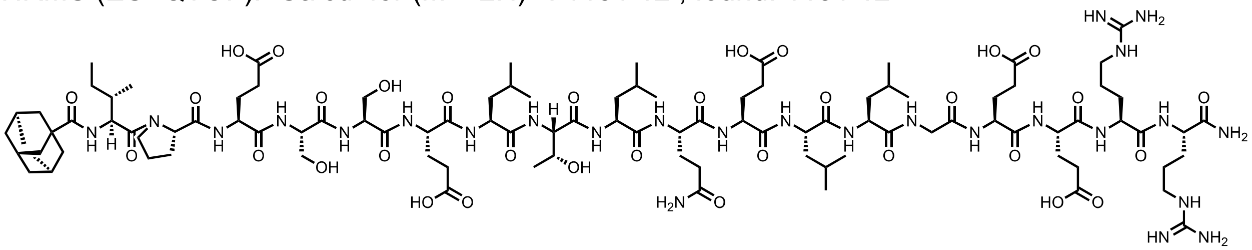

### Total ion chromatogram:

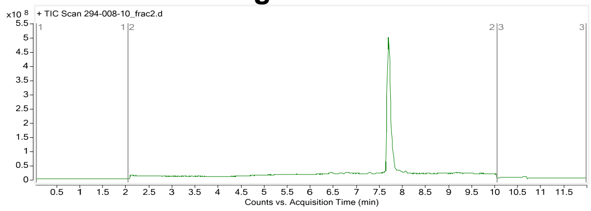


### Mass spectrum of major compound:

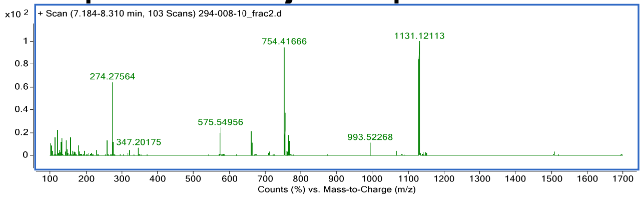


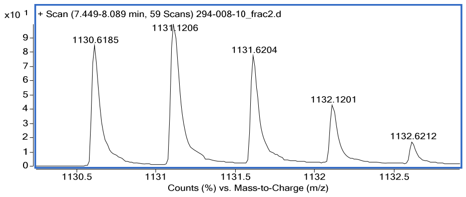


Name: peptide **N12**
HPLC: method A
HRMS (ESI-QTOF): Calcd. for (M +2H)2+: 1184.62, found: 1184.62.

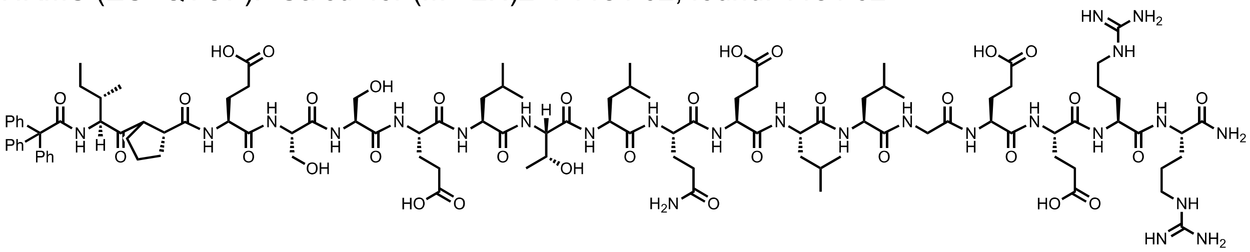

### Total ion chromatogram:

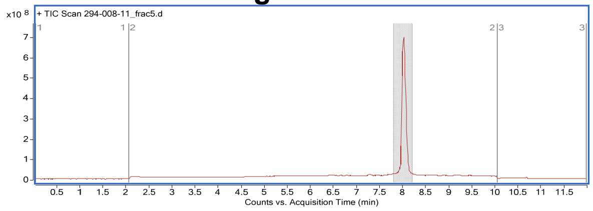


### Mass spectrum of major compound:

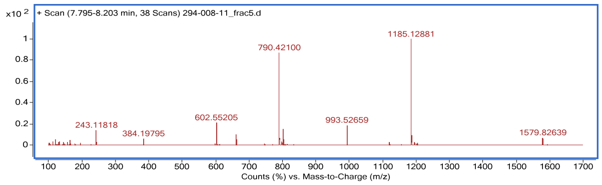


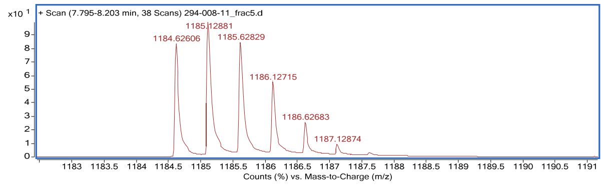


Name: peptide **N13**
HPLC: method A
HRMS (ESI-QTOF): Calcd. for (M +2H)2+: 1146.60, found: 1146.60.

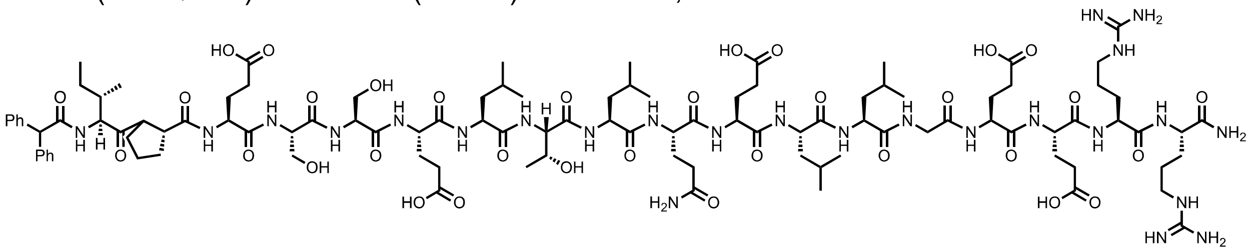

### Total ion chromatogram:

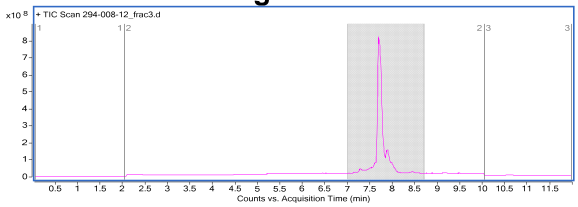


### Mass spectrum of major compound:

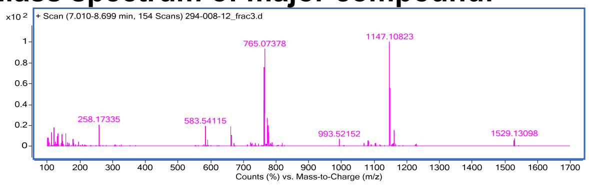


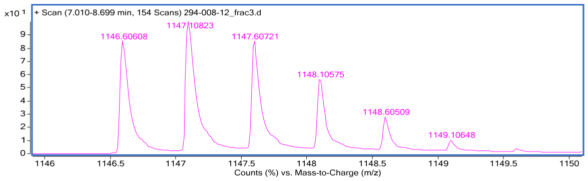


Name: peptide **N14**
HPLC: method E
HRMS (ESI-QTOF): Calcd. for $(M + 2H)^{2+}$: 1168.10, found: 1168.10.

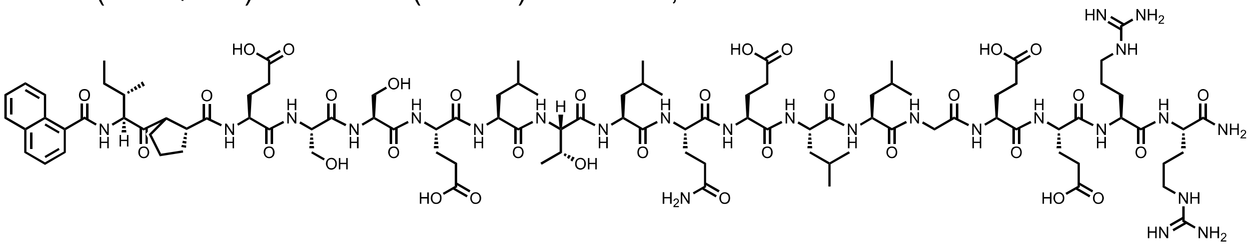

### Total ion chromatogram:

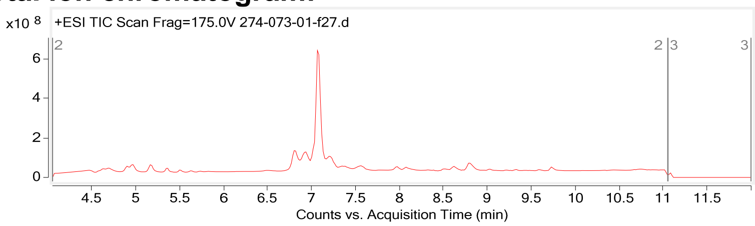


### Mass spectrum of major compound:

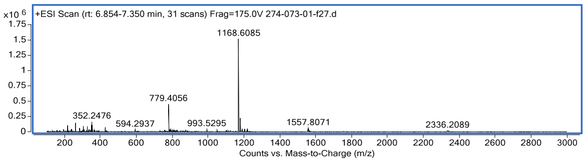


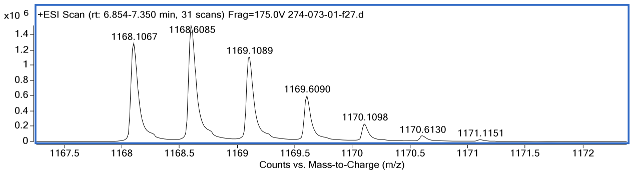


Name: peptide **N15**
HPLC: method E
HRMS (ESI-QTOF): Calcd. for (M + 2H)$^{2+}$: 1197.06, found: 1197.06

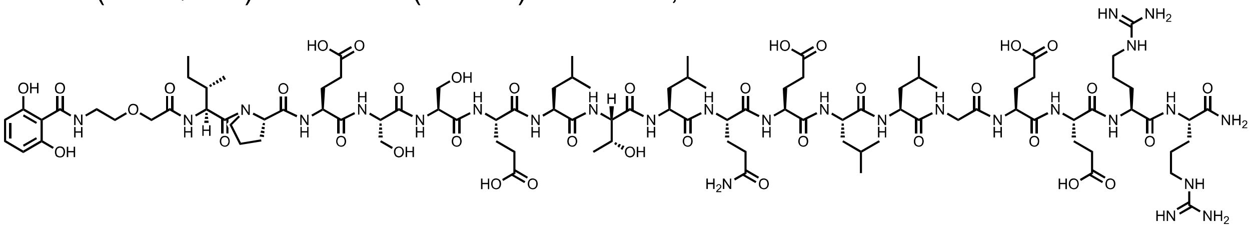

**Total ion chromatogram:**

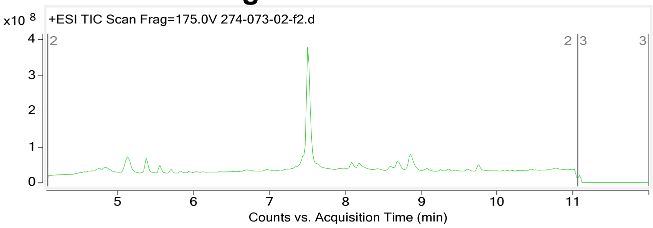


**Mass spectrum of major compound:**

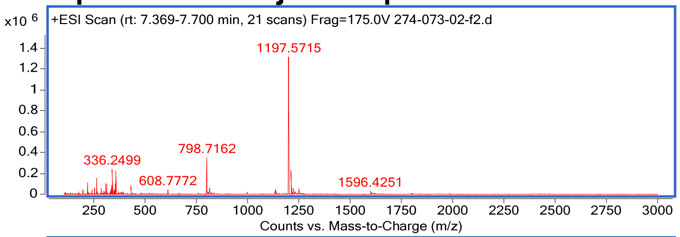


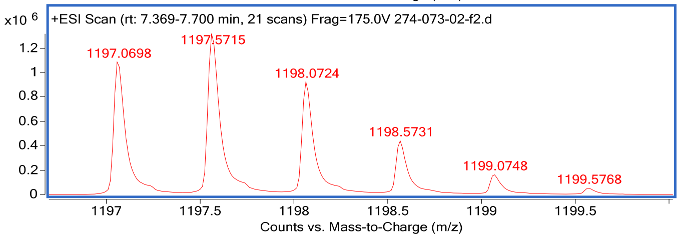


Name: peptide **N16**
HPLC: method E
HRMS (ESI-QTOF): Calcd. for (M + 2H)$^{2+}$: 1185.09, found: 1185.10

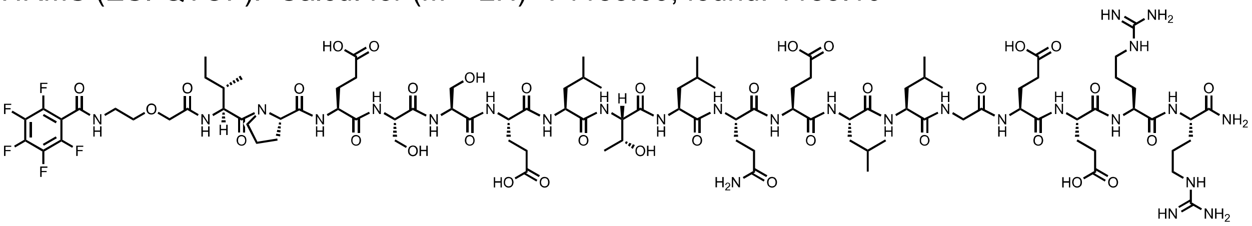

**Total ion chromatogram:**

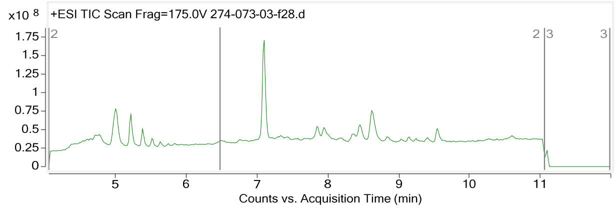


### Mass spectrum of major compound:

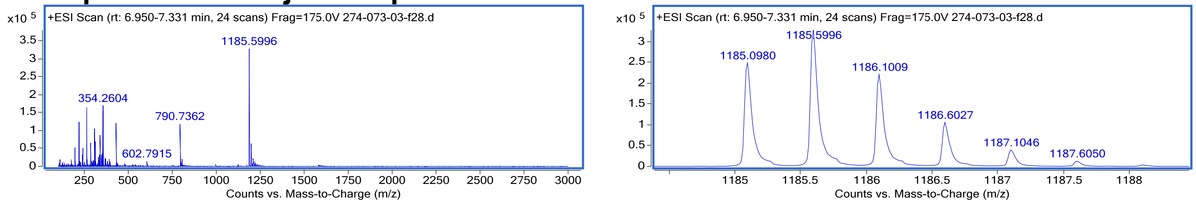


Name: peptide **17**
HPLC: method D
HRMS (ESI-QTOF): Calcd. for $(M + 2H)^{2+}$: 1235.65, found: 1235.65.

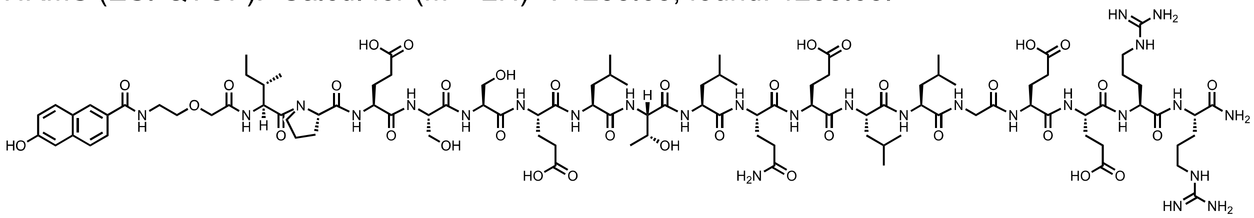

### Total ion chromatogram:

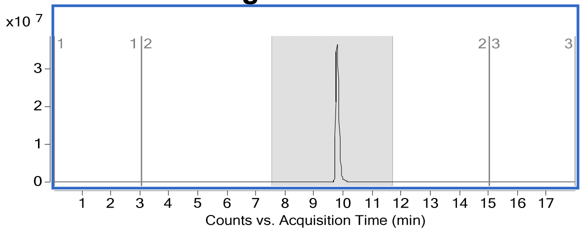


### Mass spectrum of major compound:

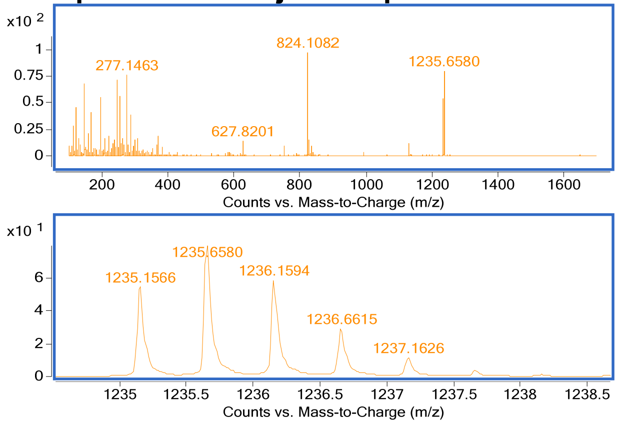


Name: peptide **N18**
HPLC: method E
HRMS (ESI-QTOF): Calcd. for (M +2H)2+: 1190.11, found: 1190.10.

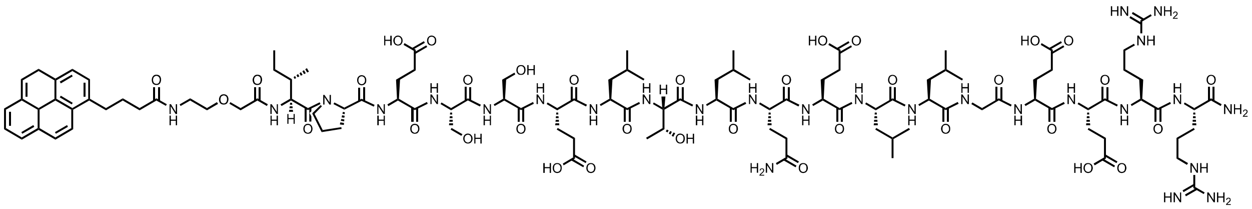

**Total ion chromatogram:**

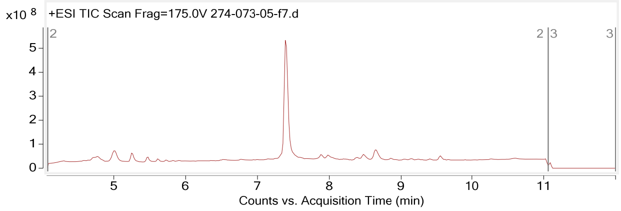


**Mass spectrum of major compound:**

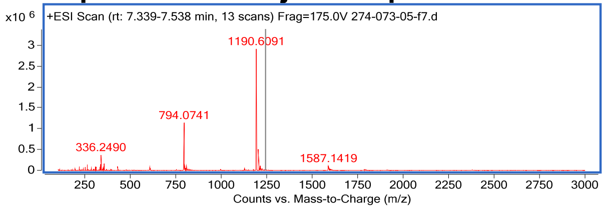


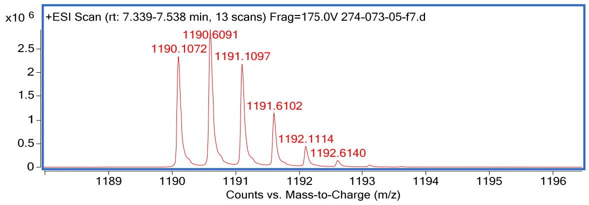


Name: peptide **N19**
HPLC: method E
HRMS (ESI-QTOF): Calcd. for $(M + 2H)^{2+}$: 1208.10, found: 1208.10

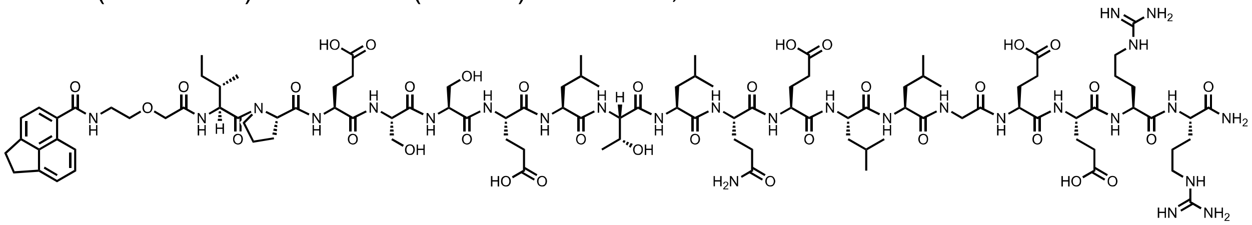

**Total ion chromatogram:**

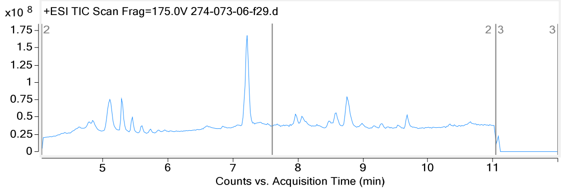


**Mass spectrum of major compound:**

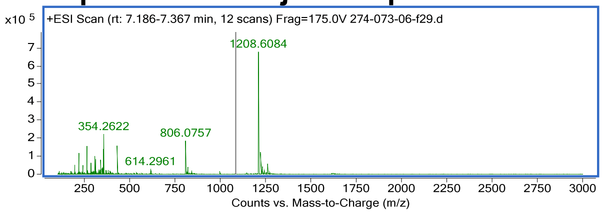


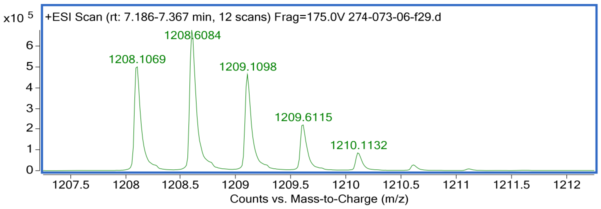

Name: peptide **N20**
HPLC: method E
HRMS (ESI-QTOF): Calcd. for $(M + 2H)^{2+}$: 1211.64, found: 1211.64

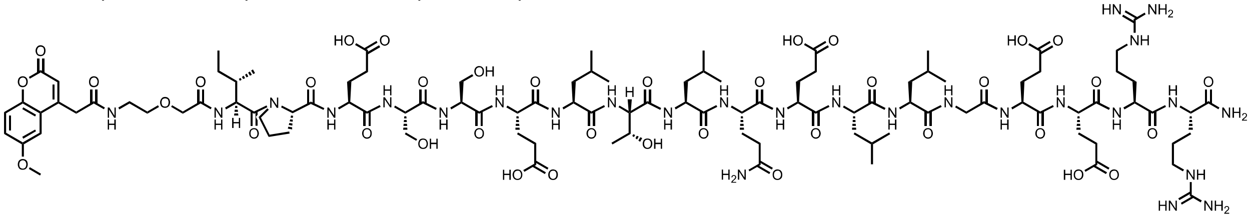

**Total ion chromatogram:**

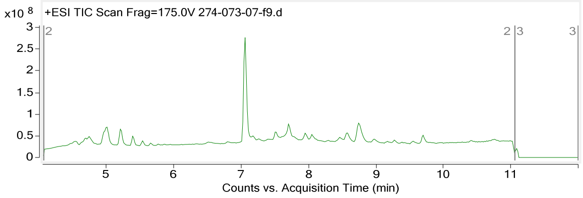


**Mass spectrum of major compound:**

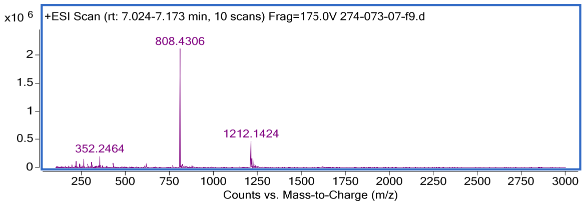


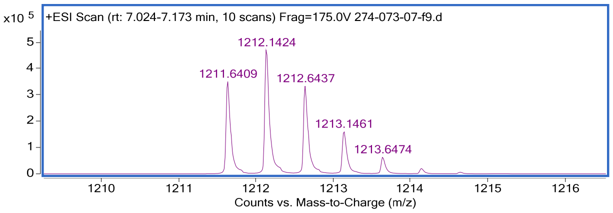


Name: peptide **N21**
HPLC: method E
HRMS (ESI-QTOF): Calcd. for (M +2H)2+: 1211.64, found: 1211.6409.

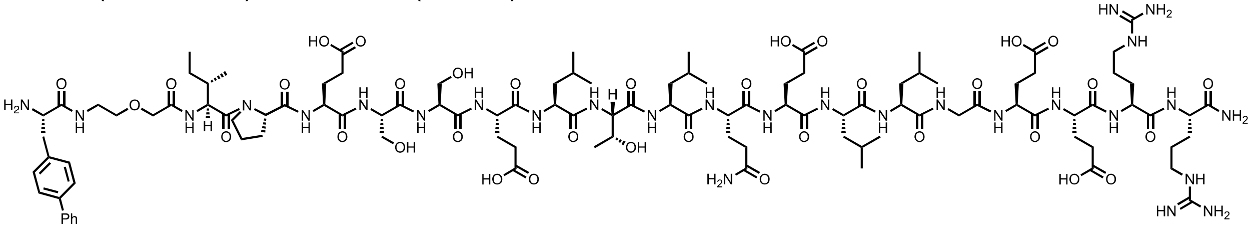

**Total ion chromatogram:**

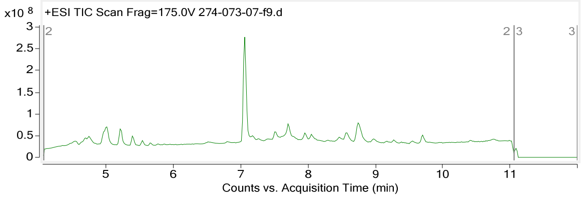


**Mass spectrum of major compound:**

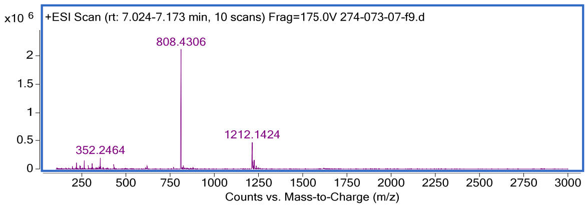


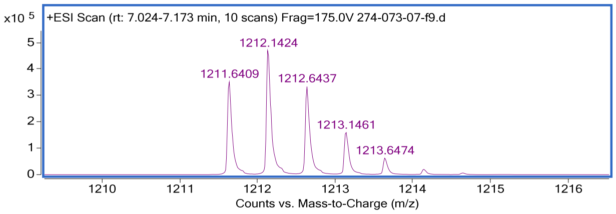


Name: peptide **N22**
HPLC: method E
HRMS (ESI-QTOF): Calcd. for (M +2H)2+: 1176.63, found: 1176.6349

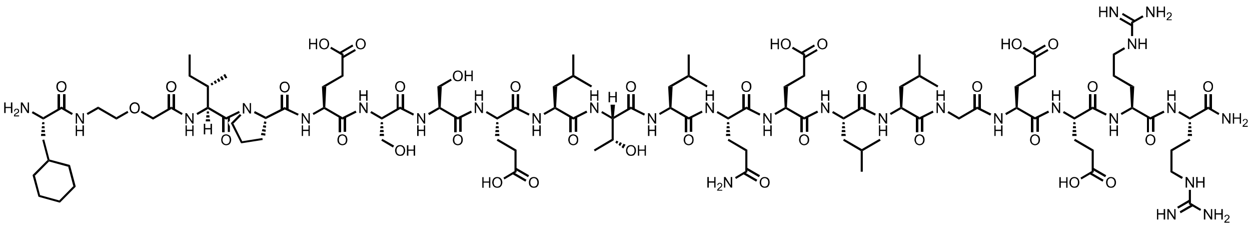

### Total ion chromatogram:

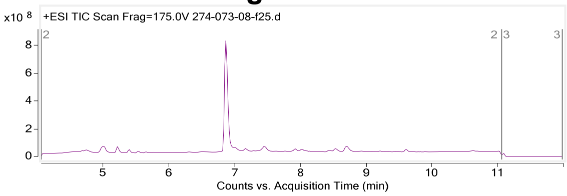


### Mass spectrum of major compound:

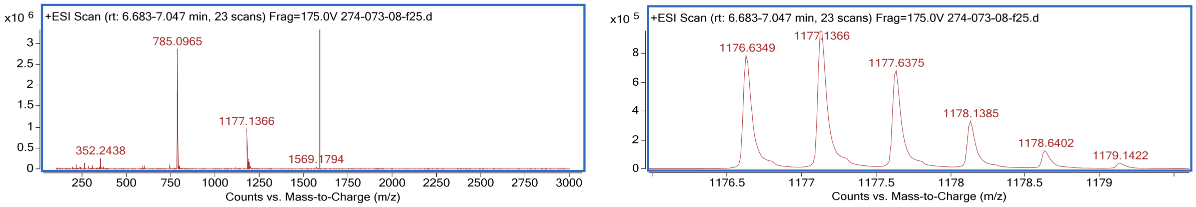


Name: peptide **N-FITC**
HPLC: method B
HRMS (ESI-QTOF): Calcd. for (M +2H)2+: 1279.59, found: 1279.59

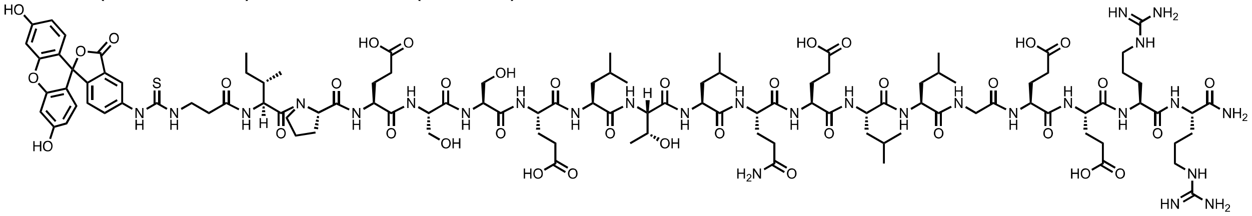

### Total ion chromatogram:

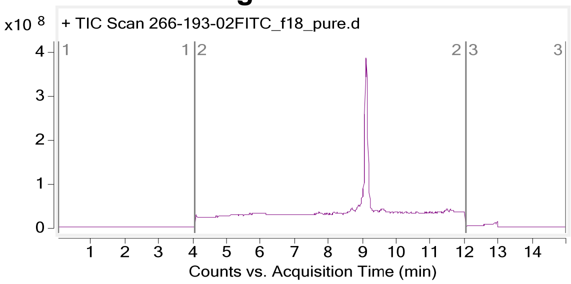


### Mass spectrum of major compound:

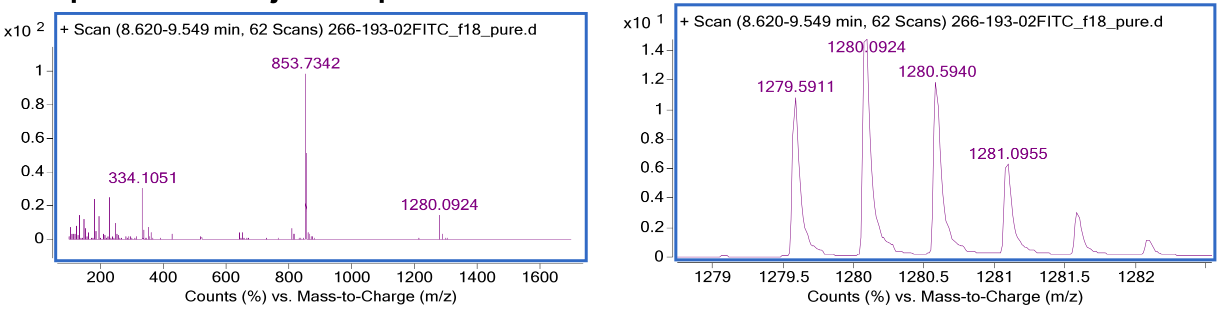

### 2.4 Peptide C1-C13 and C-FITC

Name: peptide **C-FITC**
HPLC: method B
HRMS (ESI-QTOF): Calcd. for $(M + 2H)^{2+}$: 1400.1660, found: 1400.1660

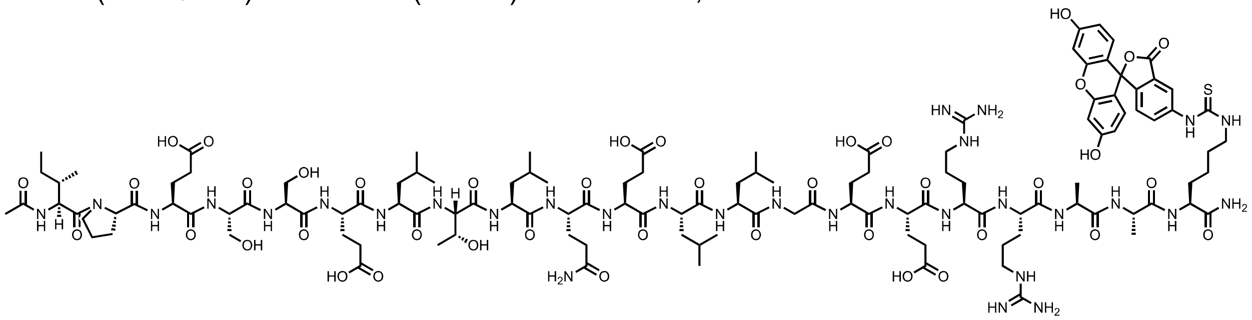

**Total ion chromatogram:**

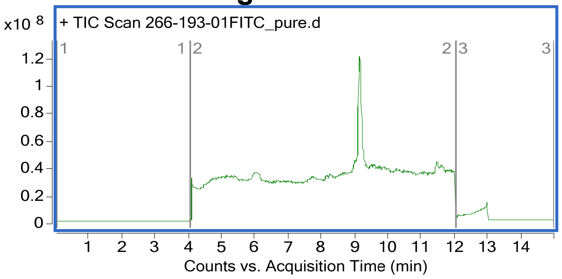


**Mass spectrum of major compound:**

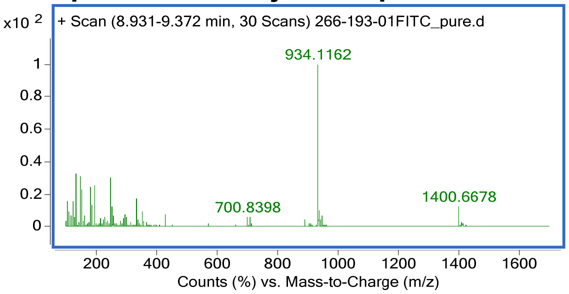


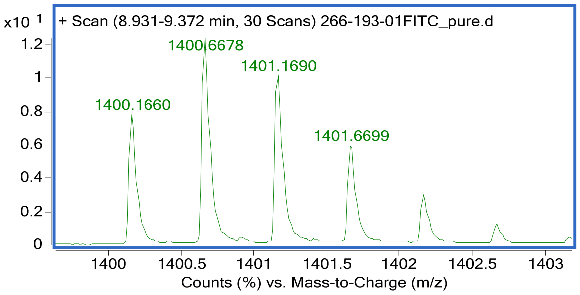


Name: peptide **C1**
HPLC: method B
HRMS (ESI-QTOF): Calcd. for $(M + 2H)^{2+}$: 1308.68, found: 1308.6887

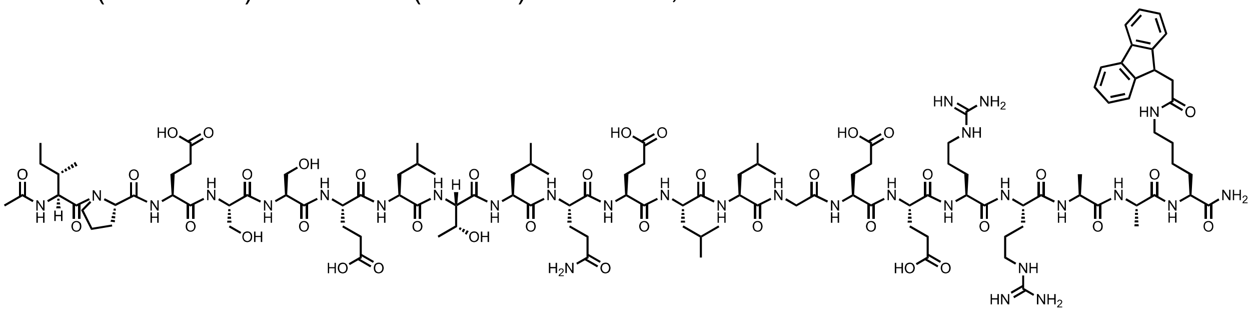

**Total ion chromatogram:**

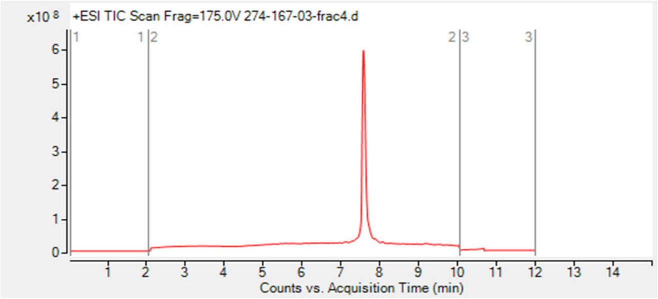


### Mass spectrum of major compound:

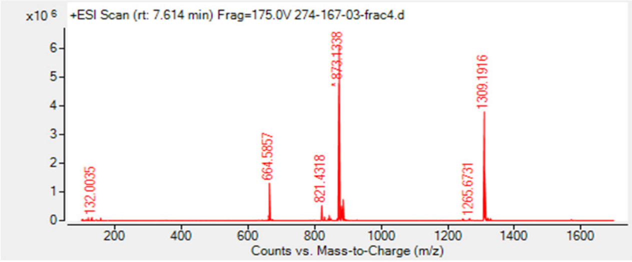


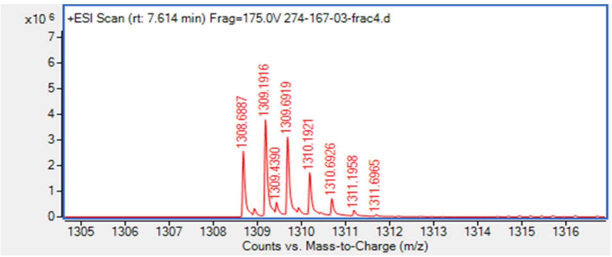


Name: peptide **C2**
HPLC: method B
HRMS (ESI-QTOF): Calcd. for $(M + 2H)^{2+}$: 1277.67, found: 1277.6777

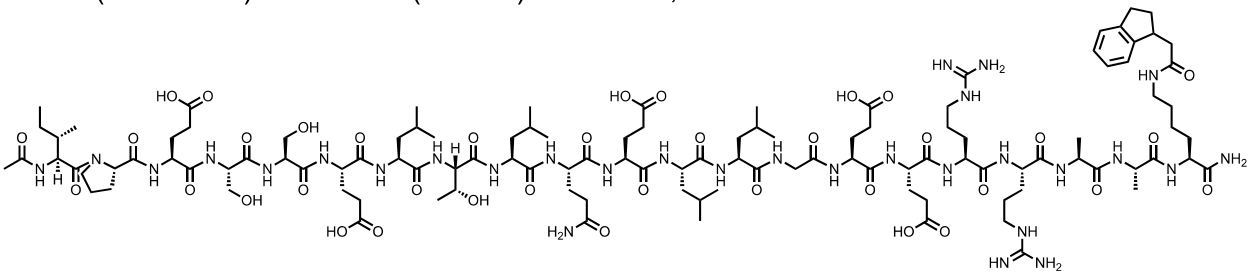

### Total ion chromatogram:

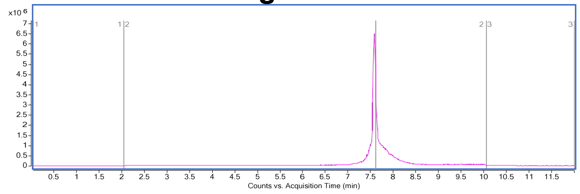


### Mass spectrum of major compound:

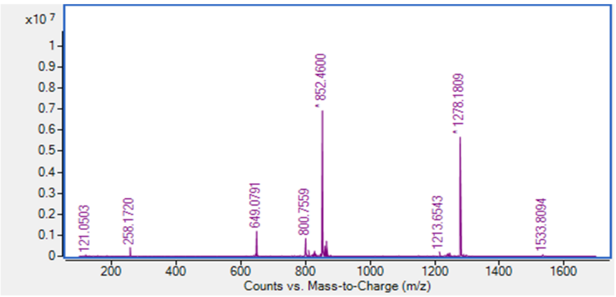


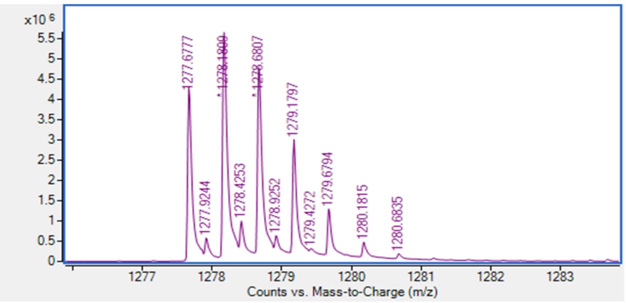


Name: peptide **C3**
HPLC: method B
HRMS (ESI-QTOF): Calcd. for $(M + 2H)^{2+}$: 1301.67, found: 1301.67.

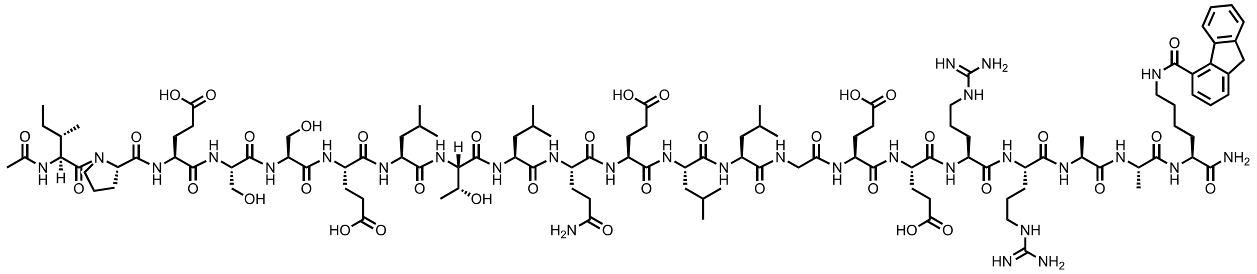

### Total ion chromatogram:

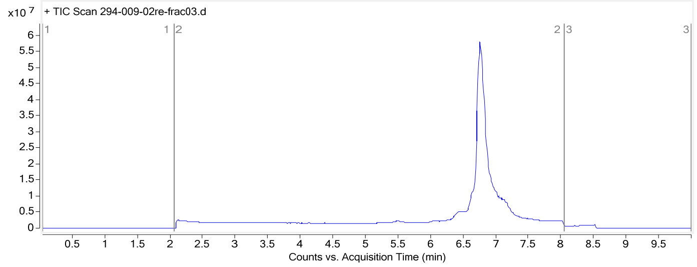


### Mass spectrum of major compound:

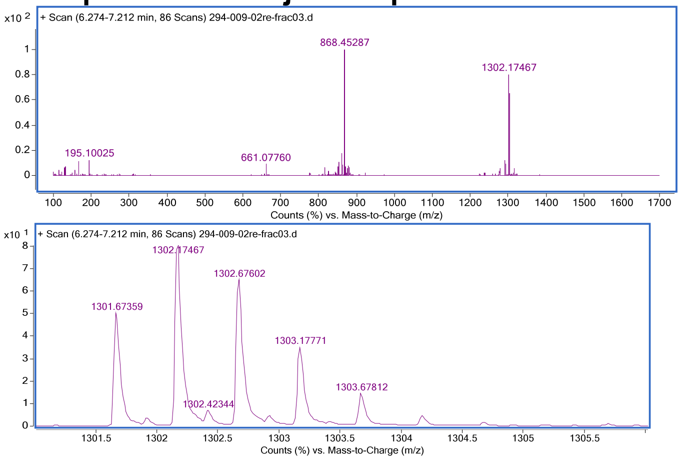


Name: peptide **C4**
HPLC: method B
HRMS (ESI-QTOF): Calcd. for $(M + 2H)^{2+}$: 1308.66, found: 1308.66

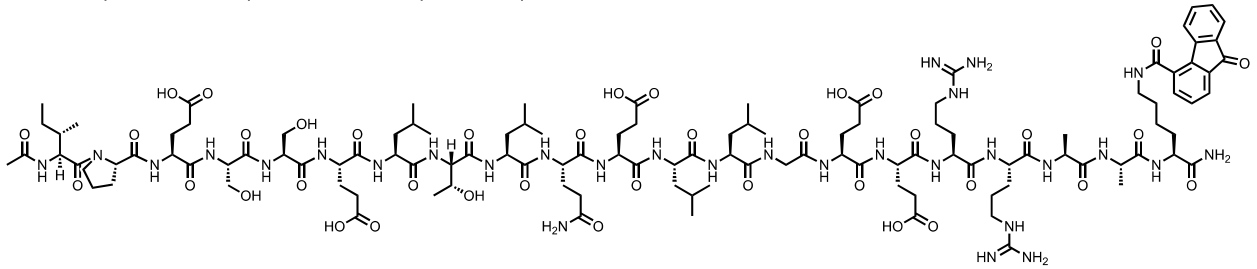

### Total ion chromatogram:

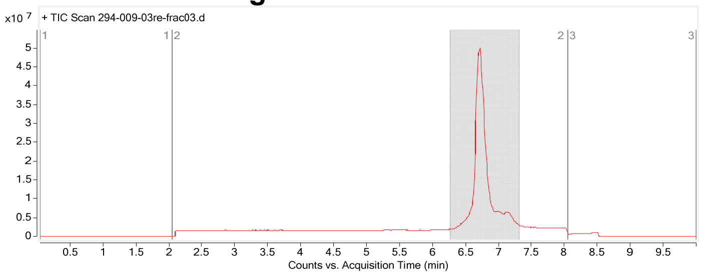

### Mass spectrum of major compound:

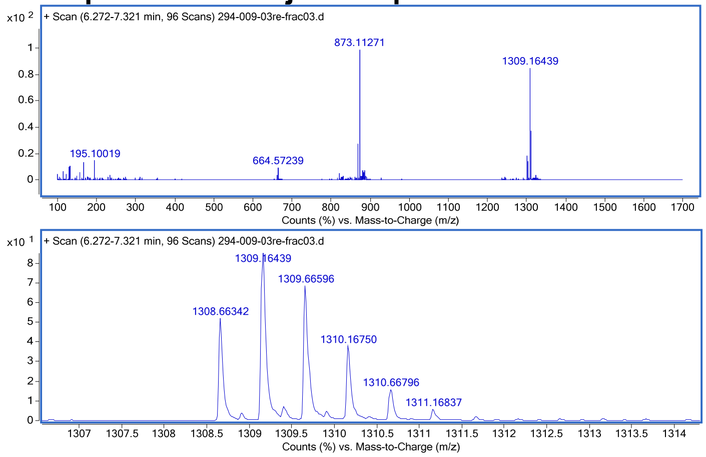


Name: peptide **C5**
HPLC: method B
HRMS (ESI-QTOF): Calcd. for $(M + 2H)^{2+}$: 1308.66, found: 1308.66.

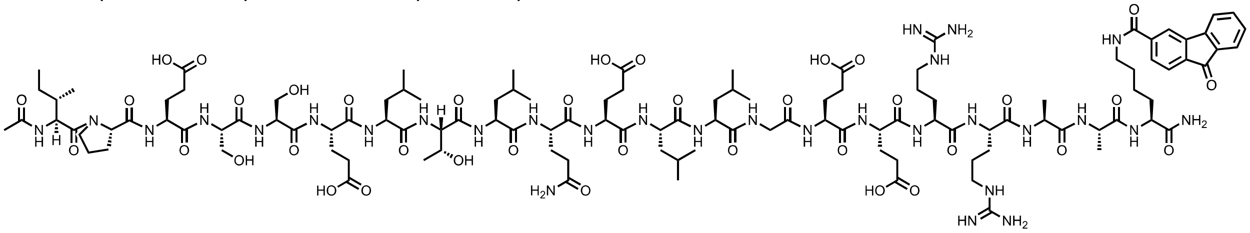

### Total ion chromatogram:

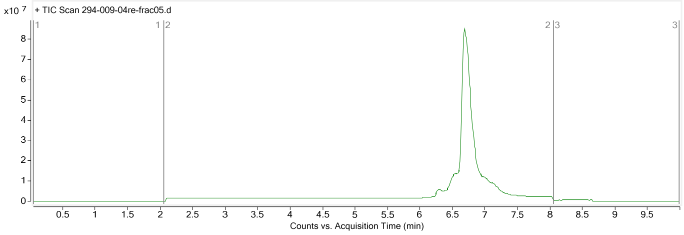


### Mass spectrum of major compound:

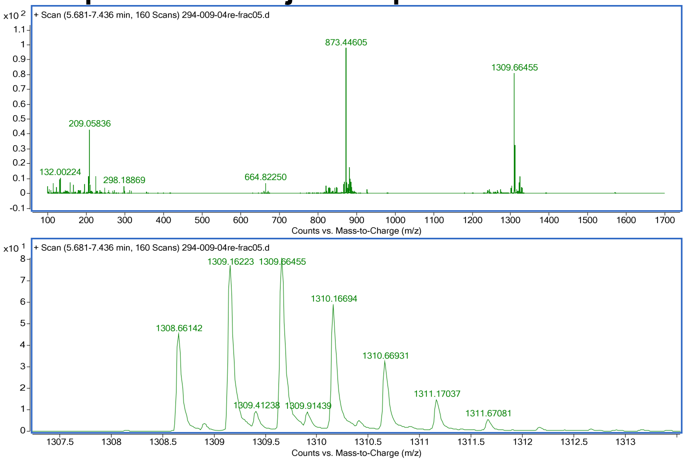


Name: peptide **C6**
HPLC: method B
HRMS (ESI-QTOF): Calcd. for $(M + 2H)^{2+}$: 1308.66, found: 1308.66.

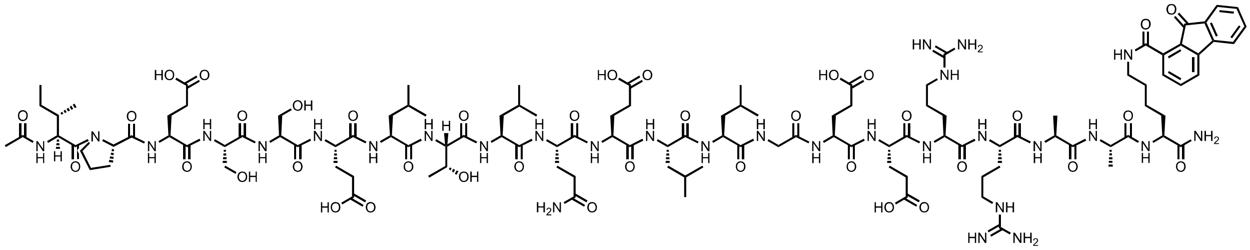

### Total ion chromatogram:

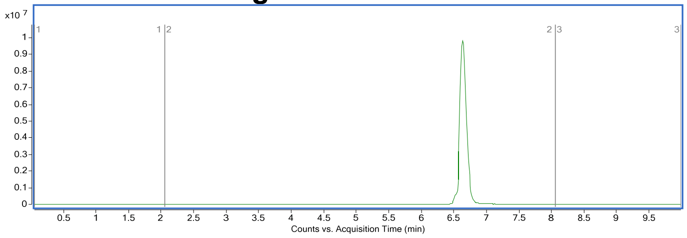


### Mass spectrum of major compound:

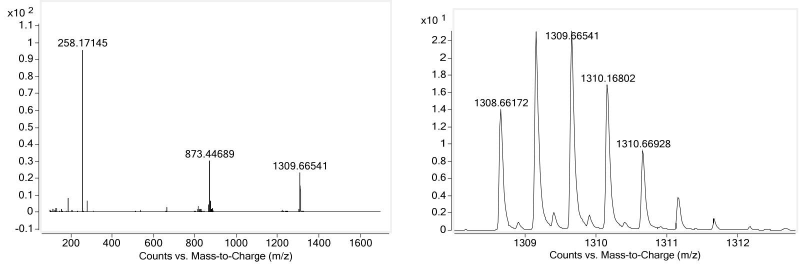


Name: peptide **C7**

HPLC: method B

HRMS (ESI-QTOF): Calcd. for $(M + 2H)^{2+}$: 1226.65, found: 1226.65

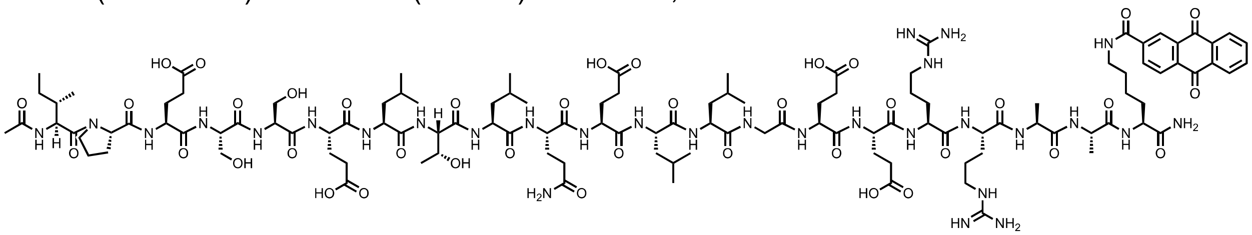

### Total ion chromatogram:

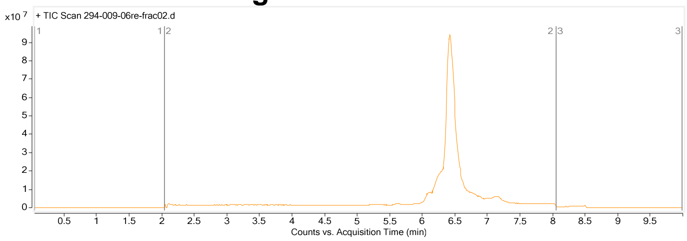


### Mass spectrum of major compound:

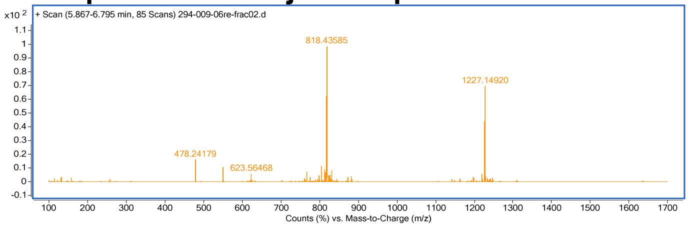

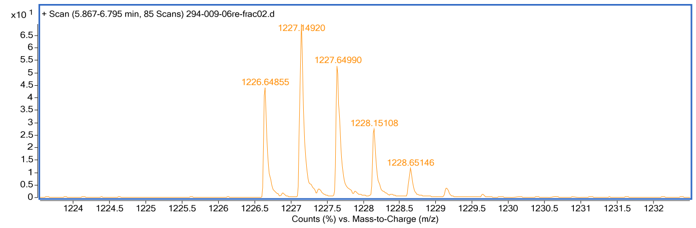


Name: peptide **C8**
HPLC: method B
HRMS (ESI-QTOF): Calcd. for $(M + 2H)^{2+}$: 1316.70, found: 1316.70.

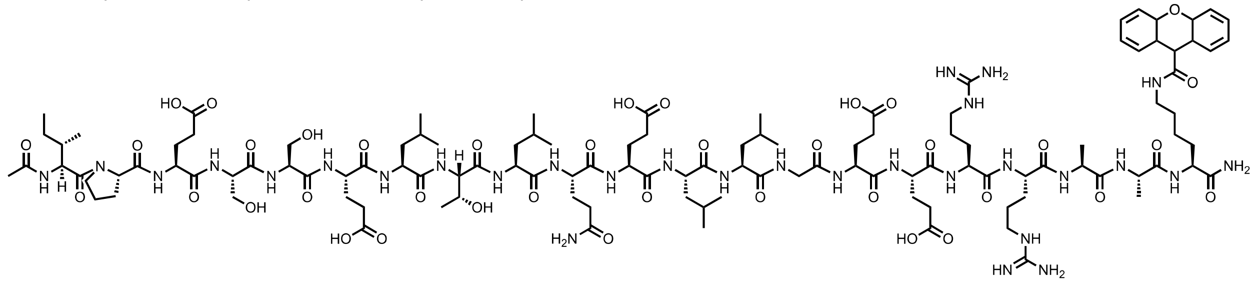

**Total ion chromatogram:**

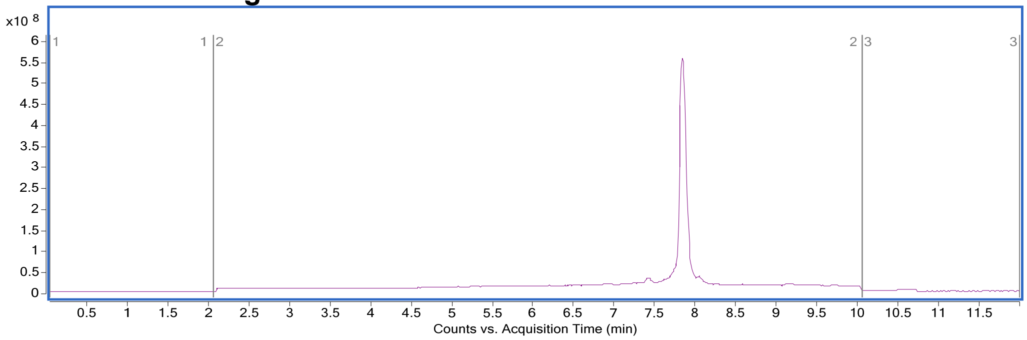


**Mass spectrum of major compound:**

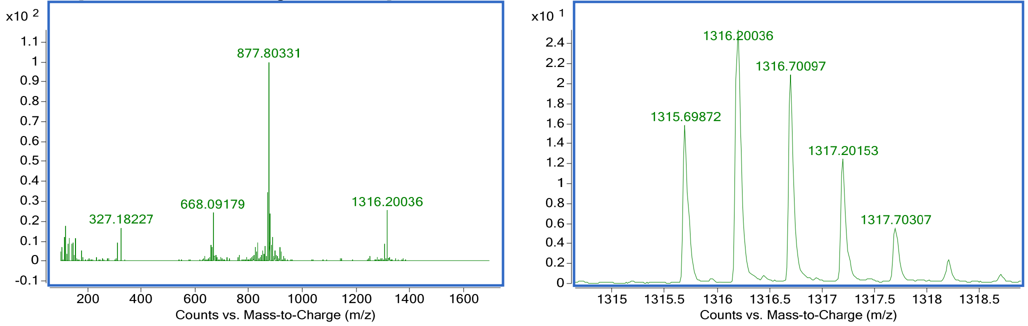


Name: peptide **C9**
HPLC: method B
HRMS (ESI-QTOF): Calcd. for $(M + 2H)^{2+}$: 1307.67, found: 1307.66

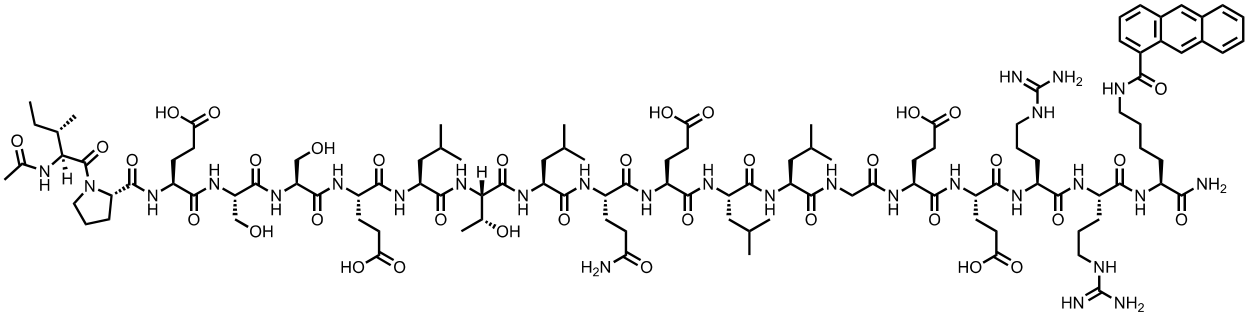

### Total ion chromatogram:

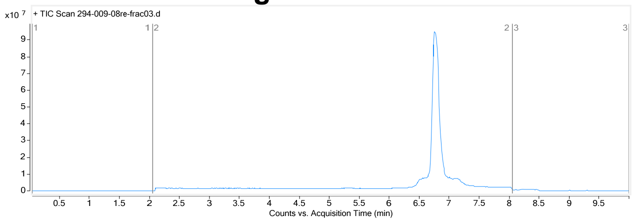


### Mass spectrum of major compound:

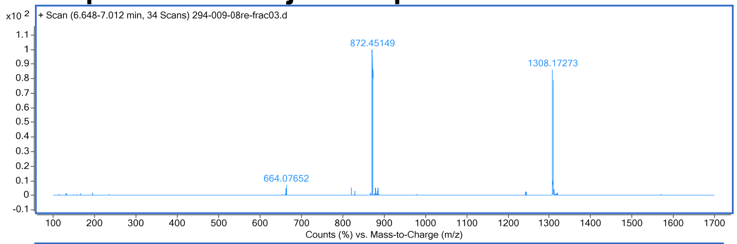


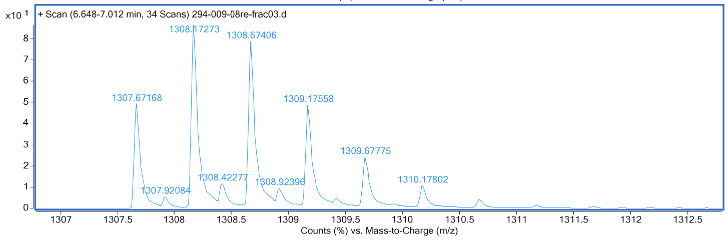


Name: peptide **C10**
HPLC: method B
HRMS (ESI-QTOF): Calcd. for $(M + 2H)^{2+}$: 1307.67, found: 1307.67.

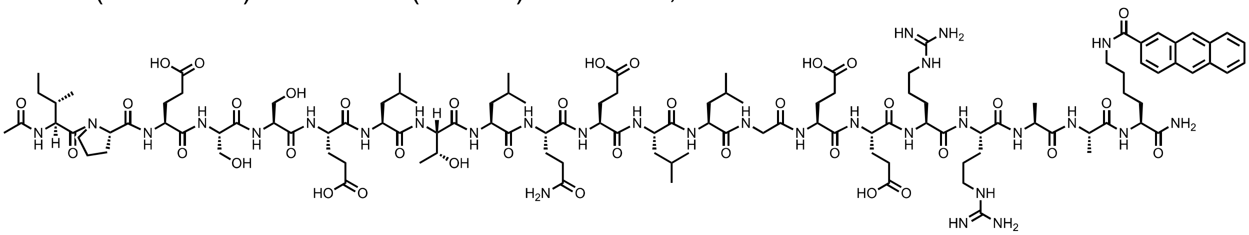

### Total ion chromatogram:

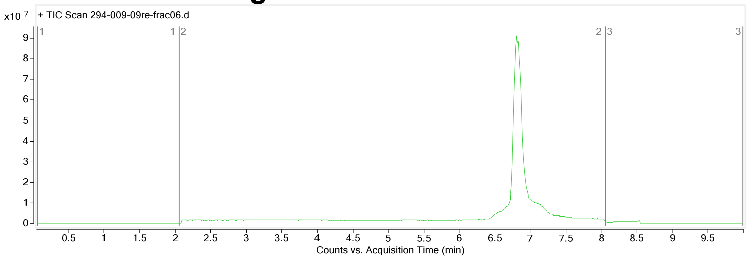


### Mass spectrum of major compound:

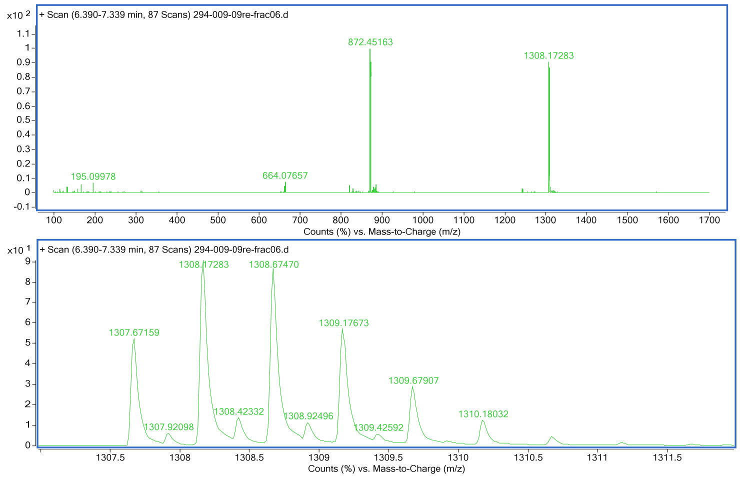


Name: peptide **C11**
HPLC: method B
HRMS (ESI-QTOF): Calcd. for (M +2H)2+: 1286.70, found: 1286.69.

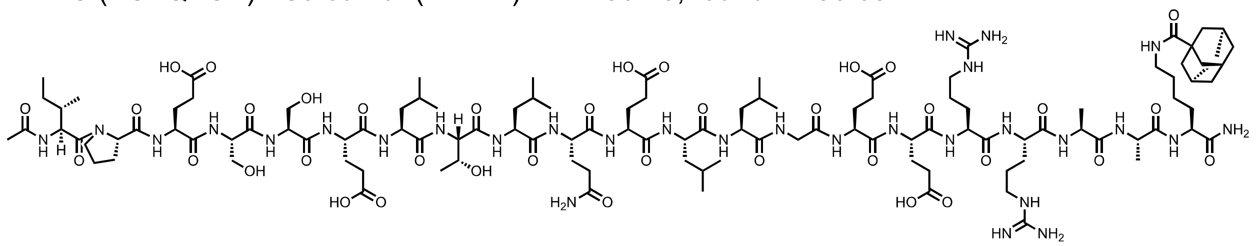

### Total ion chromatogram:

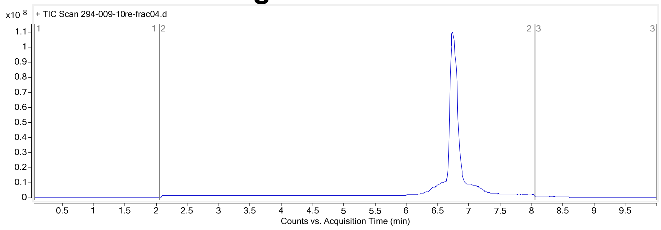


### Mass spectrum of major compound:

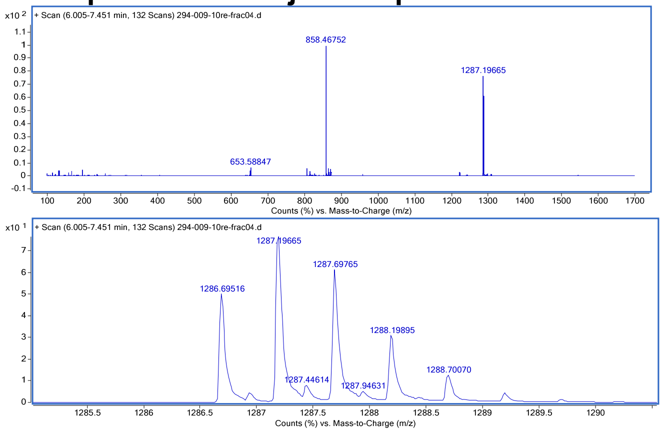


Name: peptide **C12**
HPLC: method B

HRMS (ESI-QTOF): Calcd. for (M +2H)2+: 1302.69, found: 1302.68

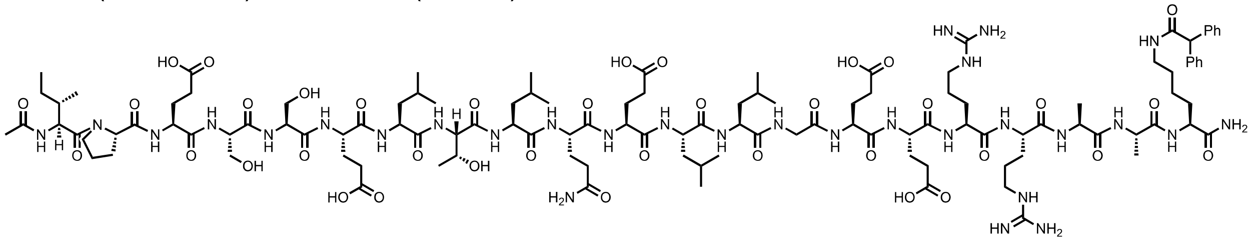

**Total ion chromatogram:**

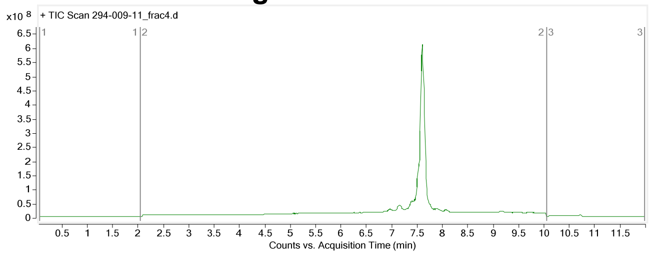


**Mass spectrum of major compound:**

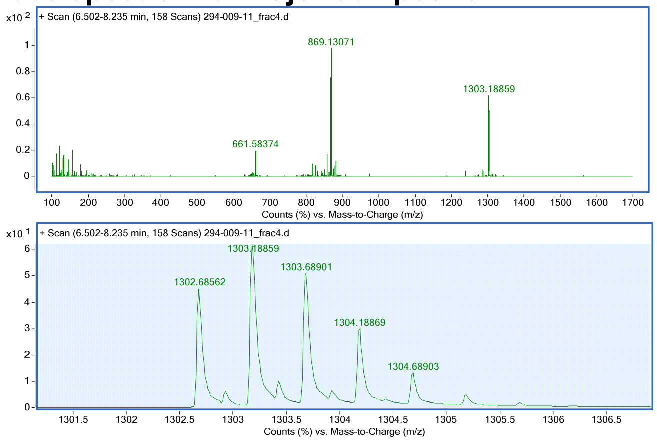


Name: peptide **C13**
HPLC: method B
HRMS (ESI-QTOF): Calcd. for (M +2H)2+: 1340.69, found: 1340.70.
**Total ion chromatogram:**

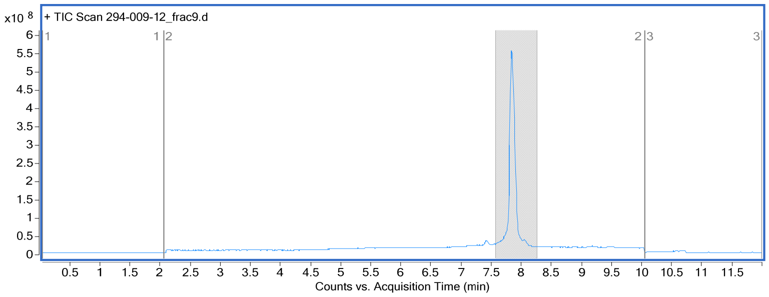


**Mass spectrum of major compound:**

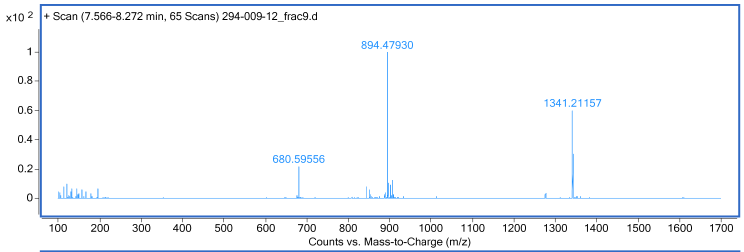


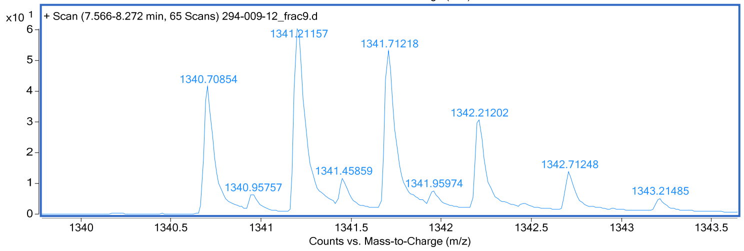


Name: peptide **C14**
HPLC: method B
HRMS (ESI-QTOF): Calcd. for (M +2H)2+: 1282.66, found: 1282.67

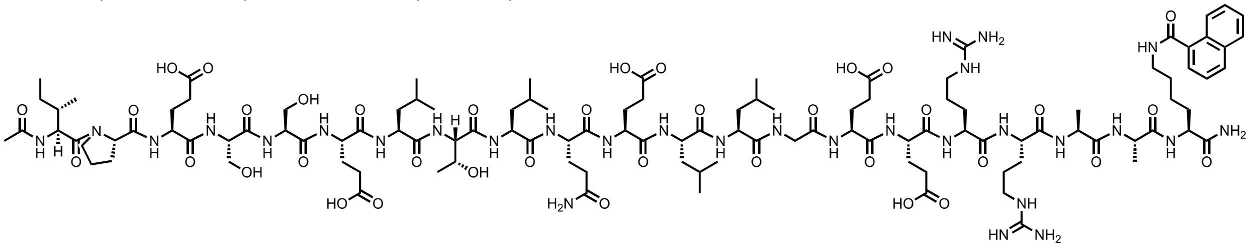

### Total ion chromatogram:

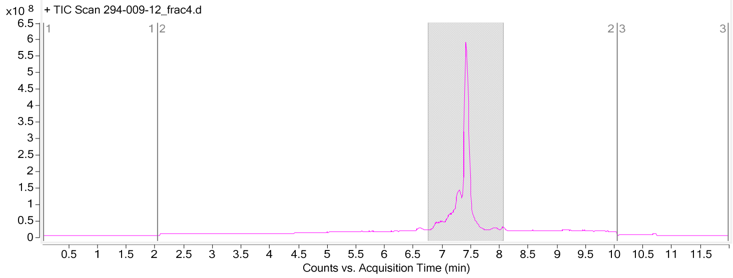


### Mass spectrum of major compound:

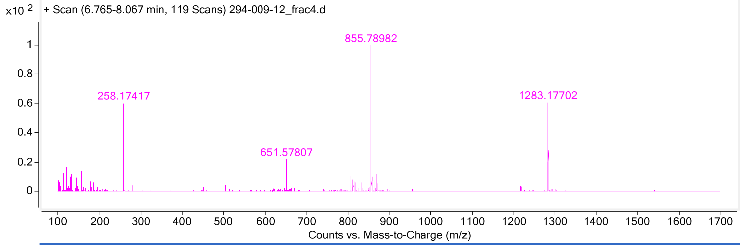


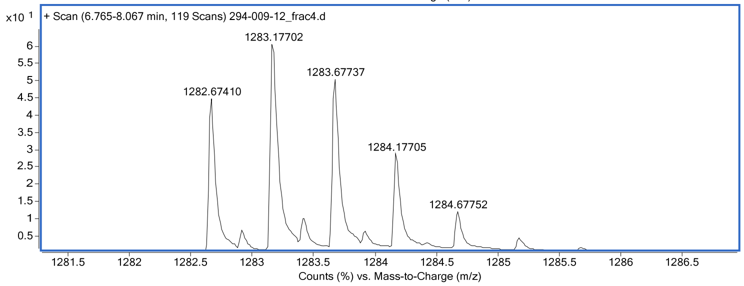

Name: peptide **C15**
HPLC: method B
HRMS (ESI-QTOF): Calcd. for (M +2H)2+: 1265.66, found: 1265.66

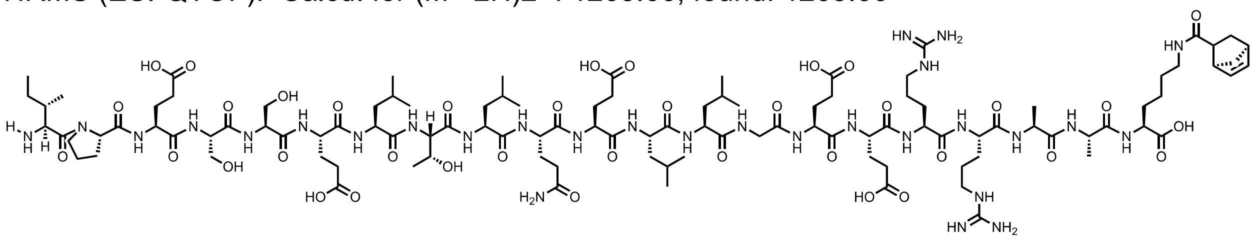

### Total ion chromatogram:

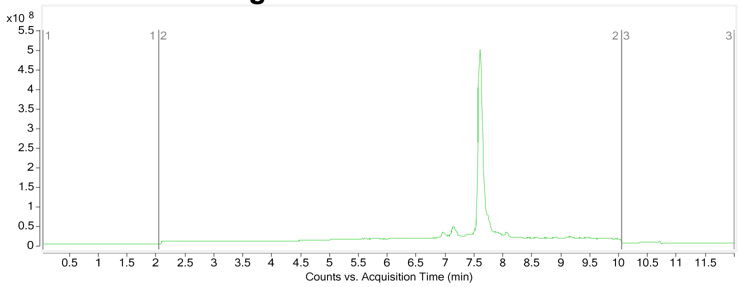


### Mass spectrum of major compound:

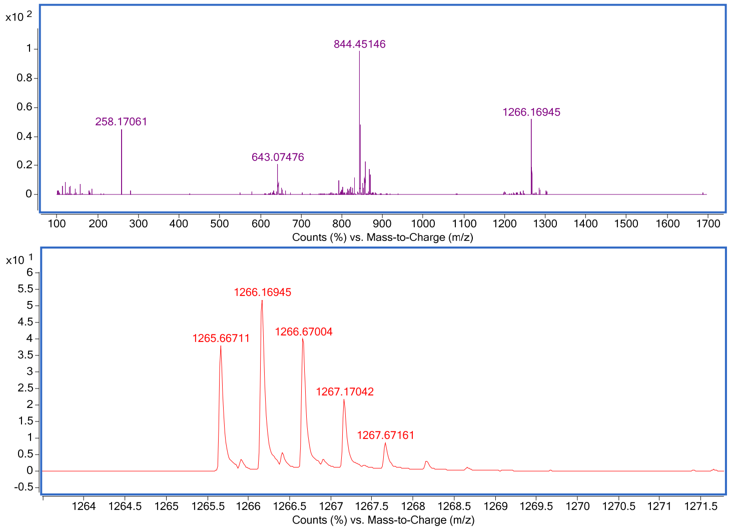

### 2.5 Peptide A1-A17

Name: peptide **A1**
HPLC: method B
HRMS (ESI-QTOF): Calcd. for (M +2H)2+: 1181.06, found: 1181.06.

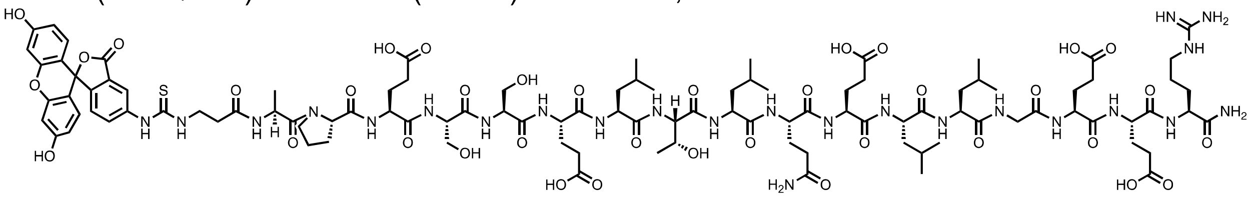

**Total ion chromatogram:**

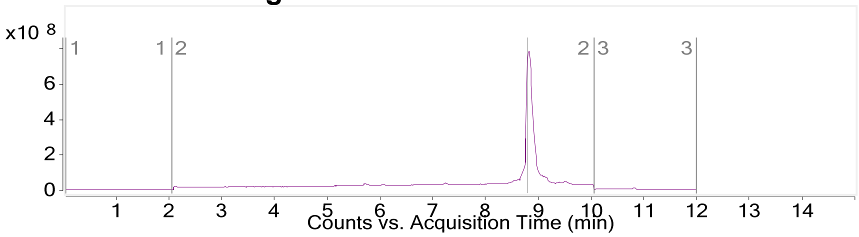


**Mass spectrum of major compound:**

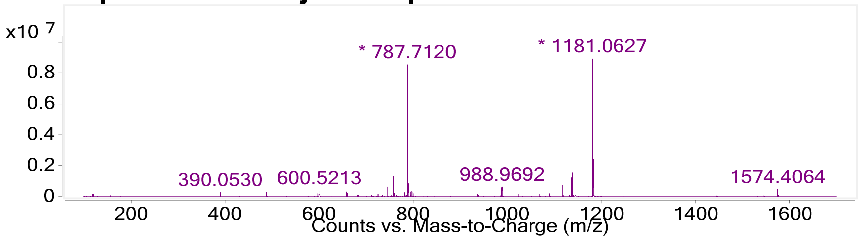


Name: peptide **A2**
HPLC: method B
HRMS (ESI-QTOF): Calcd. for (M +2H)2+: 1189.05, found: 1189.04.

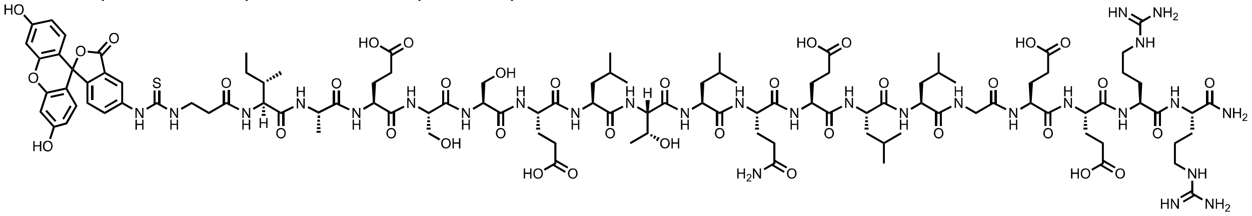

**Total ion chromatogram:**

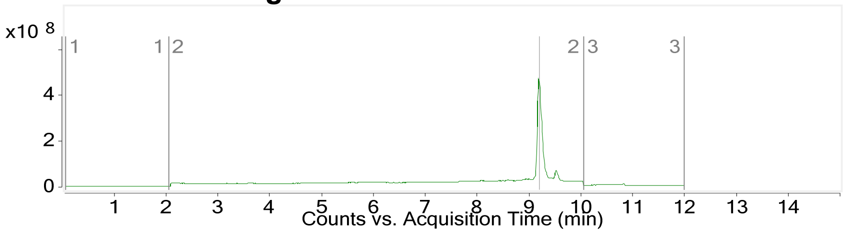


**Mass spectrum of major compound:**

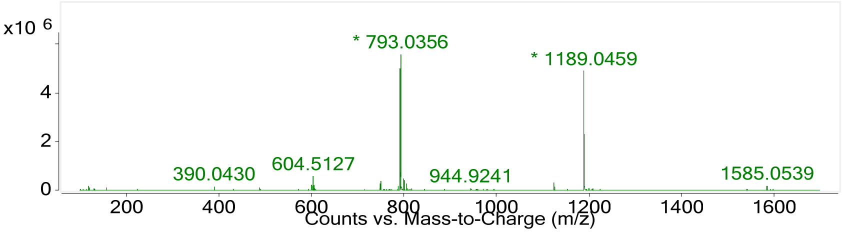


Name: peptide **A3**
HPLC: method B
HRMS (ESI-QTOF): Calcd. for (M +2H)2+: 1173.05, found: 1173.05.

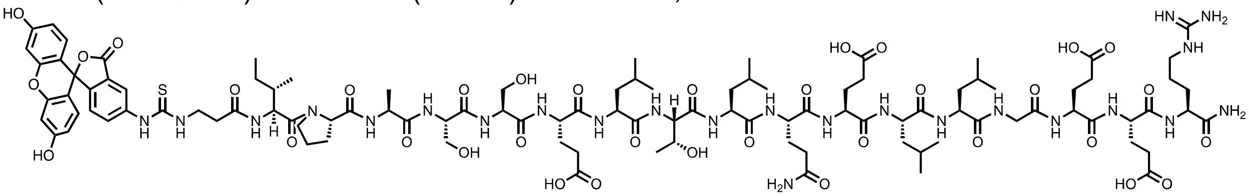

**Total ion chromatogram:**

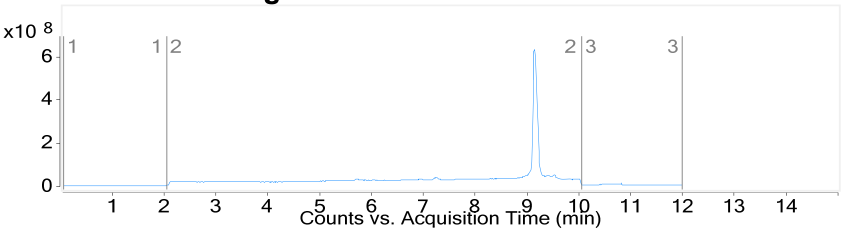


**Mass spectrum of major compound:**

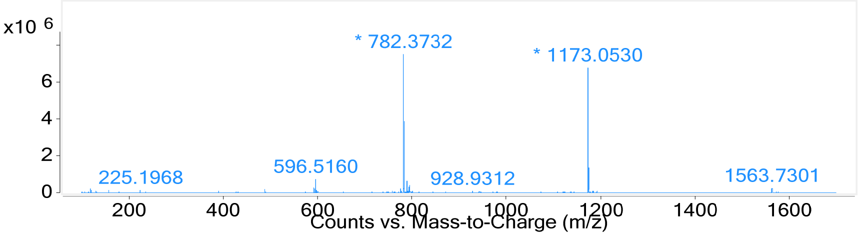


Name: peptide **A4**
HPLC: method B
HRMS (ESI-QTOF): Calcd. for (M +2H)2+: 1194.05 found: 1194.06.

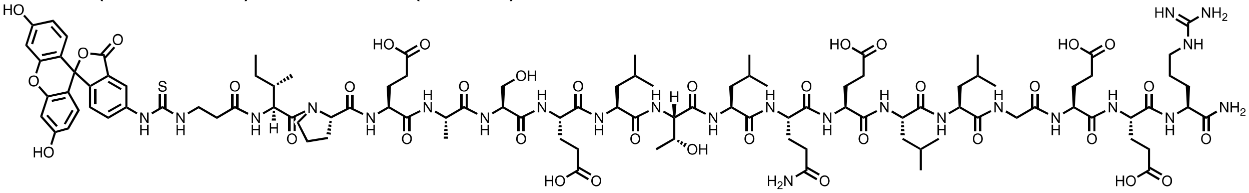

**Total ion chromatogram:**

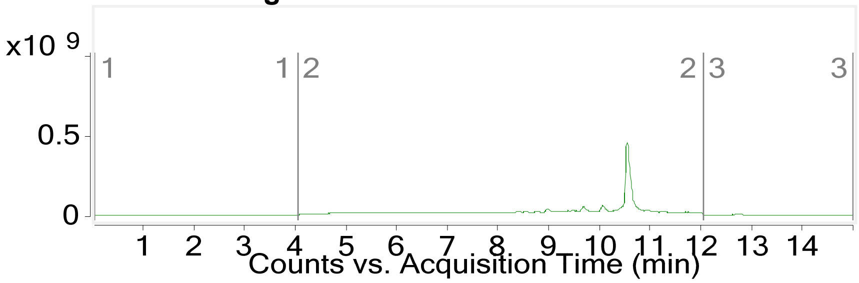


**Mass spectrum of major compound:**

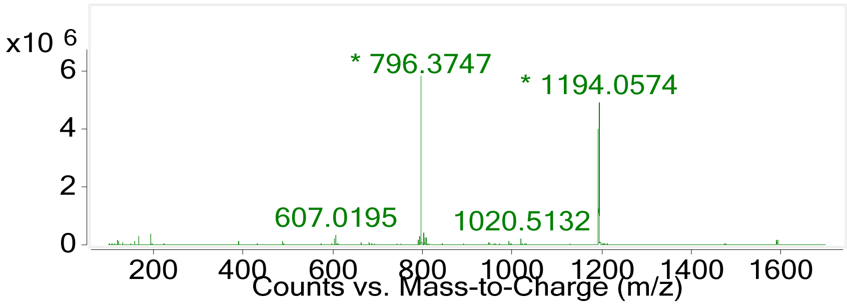


Name: peptide **A5**
HPLC: method B
HRMS (ESI-QTOF): Calcd. for (M +2H)2+: 1194.06, found: 1194.06.

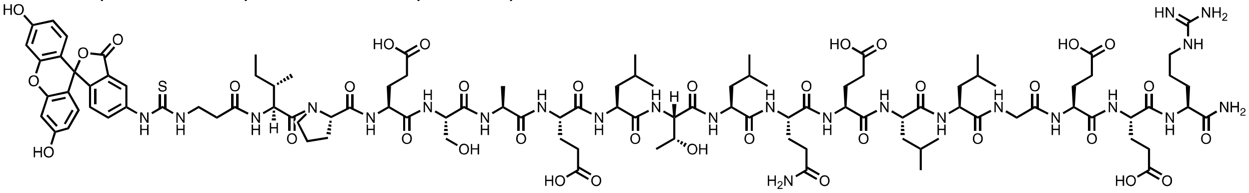

**Total ion chromatogram:**

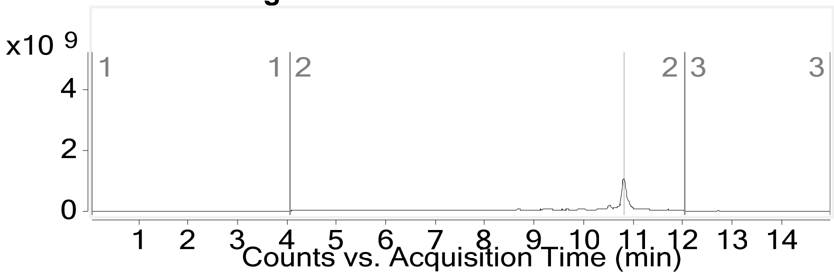


**Mass spectrum of major compound:**

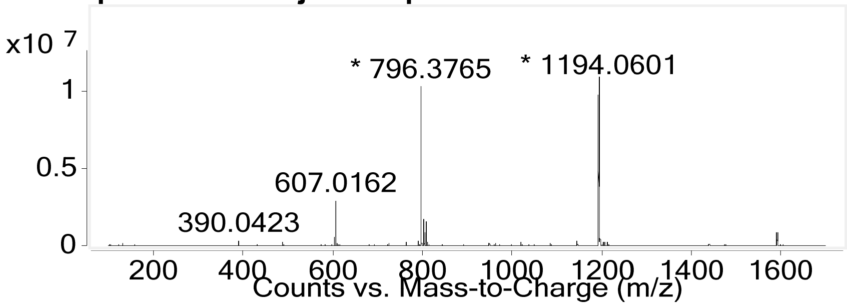


Name: peptide **A6**
HPLC: method B
HRMS (ESI-QTOF): Calcd. for (M +2H)2+: 1173.05, found: 1173.05

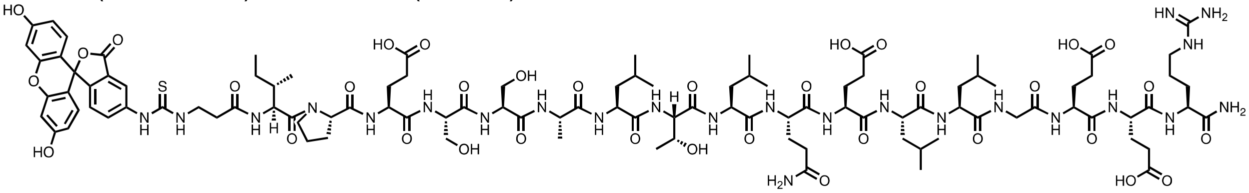

**Total ion chromatogram:**

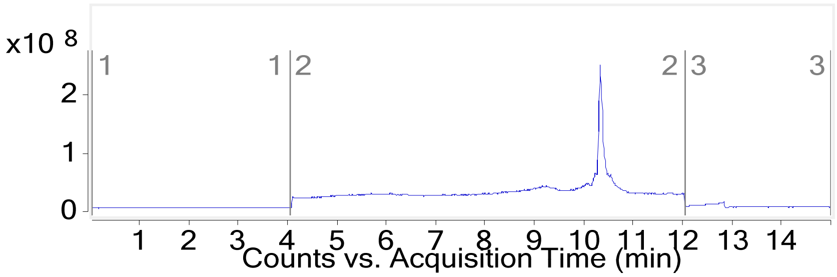


**Mass spectrum of major compound:**

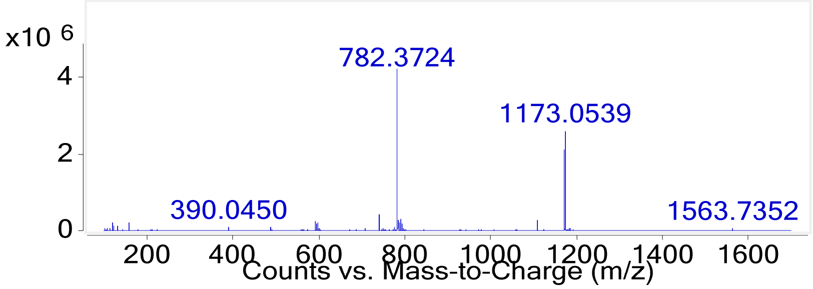


Name: peptide **A7**
HPLC: method B
HRMS (ESI-QTOF): Calcd. for (M +2H)2+: 1181.06, found: 1181.06.

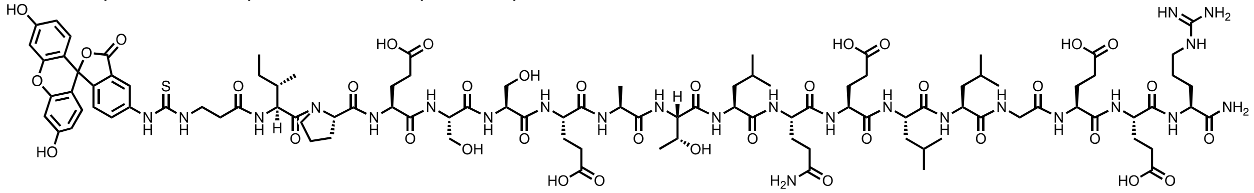

**Total ion chromatogram:**

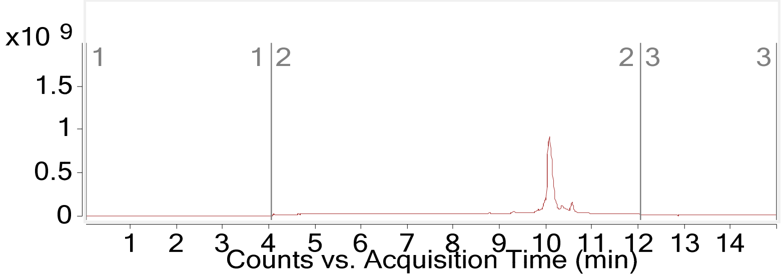


**Mass spectrum of major compound:**

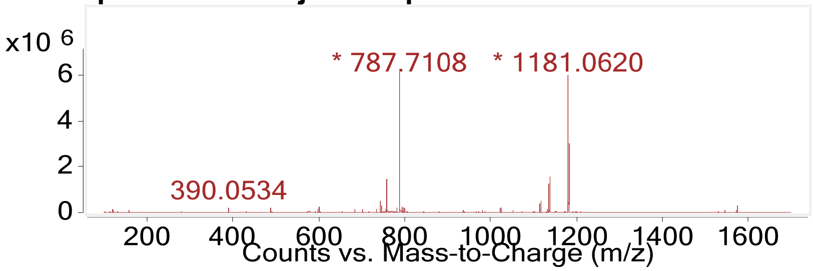


Name: peptide **A8**
HPLC: method B
HRMS (ESI-QTOF): Calcd. for (M +2H)2+: 1187.05, found: 1187.05.

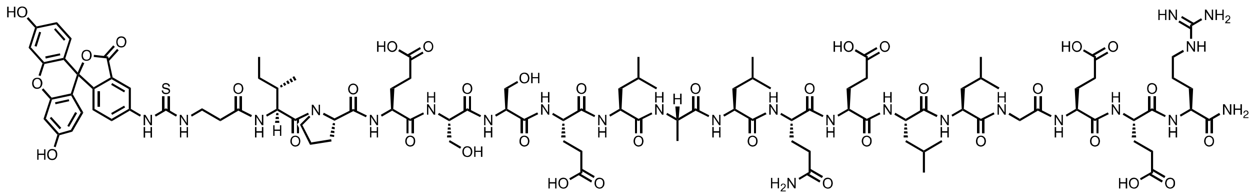

**Total ion chromatogram:**

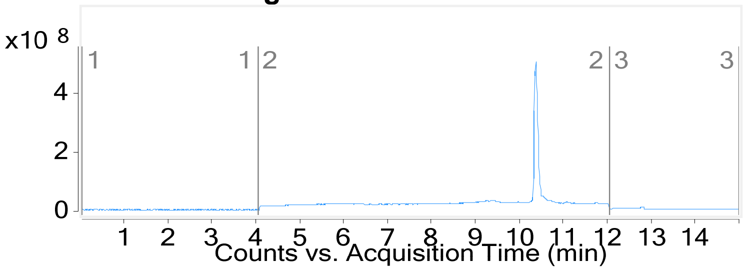


**Mass spectrum of major compound:**

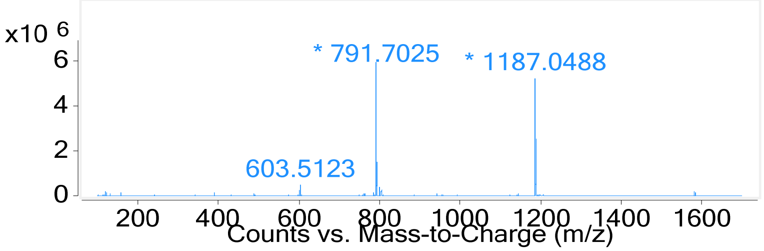


Name: peptide **A9**
HPLC: method B
HRMS (ESI-QTOF): Calcd. for (M +2H)2+: 1181.05, found: 1181.05.

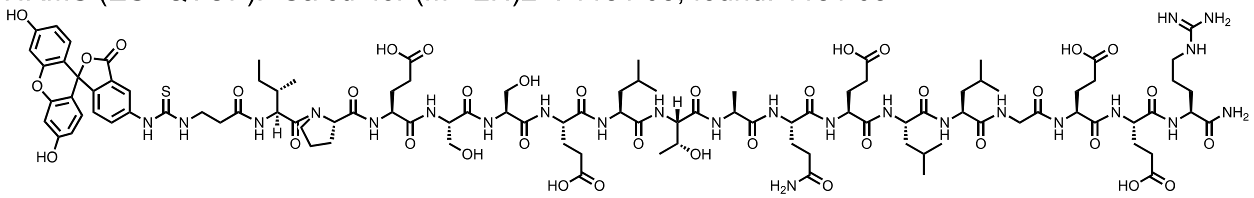

**Total ion chromatogram:**

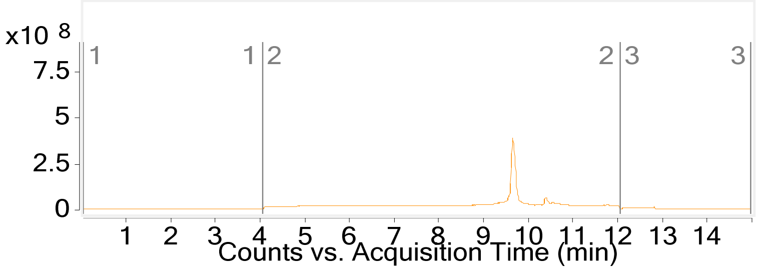


**Mass spectrum of major compound:**

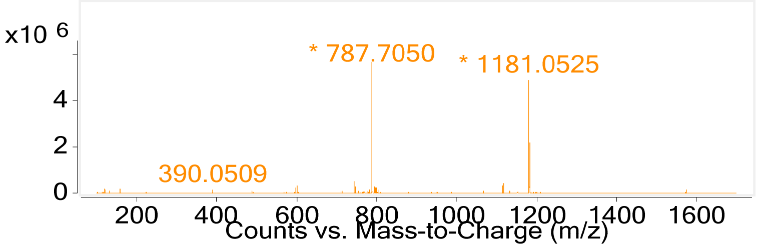


Name: peptide **A10**
HPLC: method B

HRMS (ESI-QTOF): Calcd. for (M +2H)2+: 1173.54, found: 1173.53.

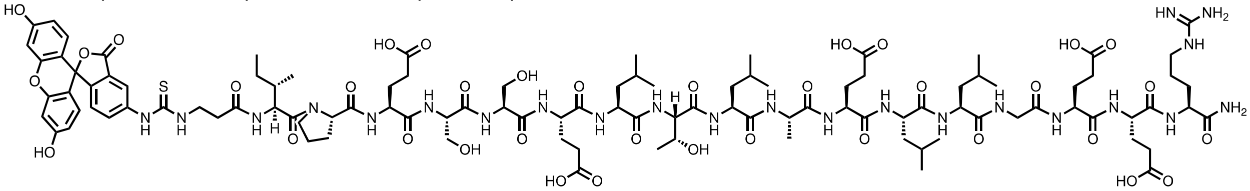

**Total ion chromatogram:**

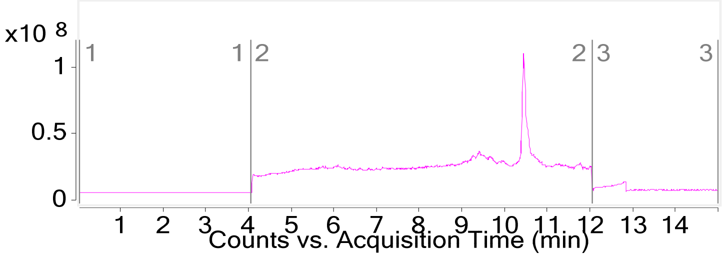


**Mass spectrum of major compound:**

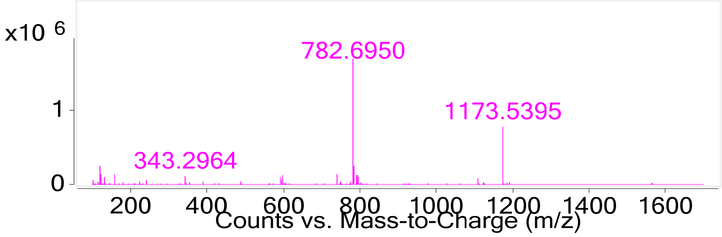


Name: peptide **A11**

HPLC: method B

HRMS (ESI-QTOF): Calcd. for $(M + 2H)^{2+}$: 1173.16, found: 1173.16.

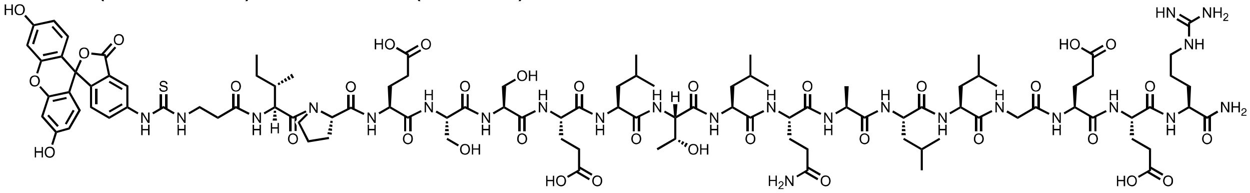

**Total ion chromatogram:**

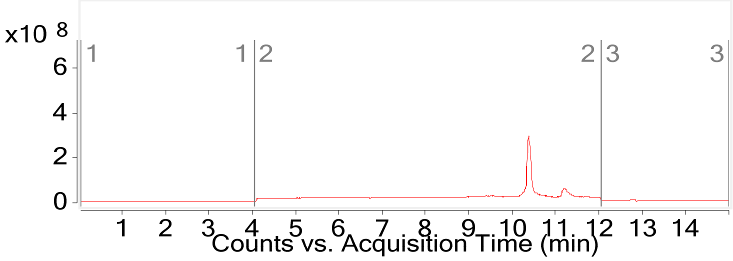


**Mass spectrum of major compound:**

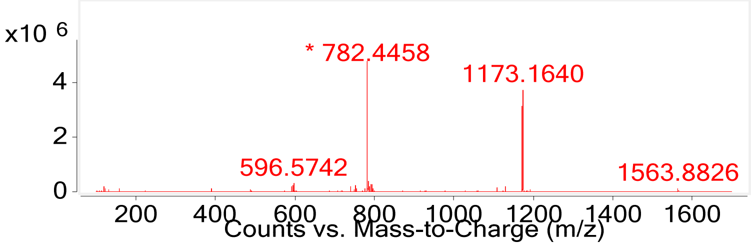


Name: peptide **A12**

HPLC: method B

HRMS (ESI-QTOF): Calcd. for $(M + 2H)^{2+}$:1181.04, found: 1181.04.

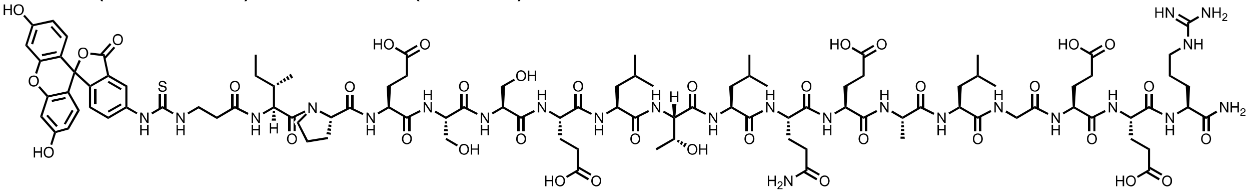

**Total ion chromatogram:**

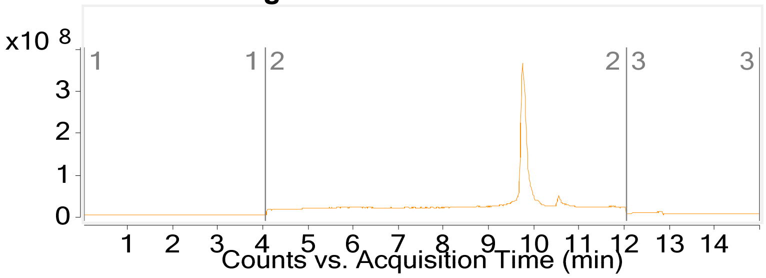


**Mass spectrum of major compound:**

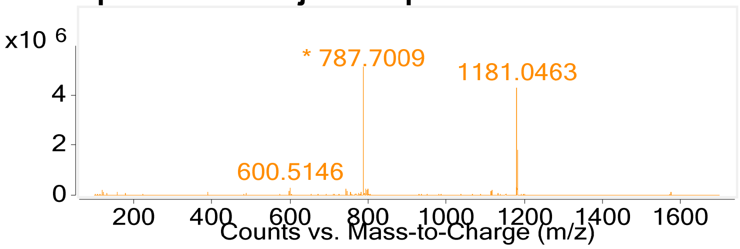


Name: peptide **A13**

HPLC: method B

HRMS (ESI-QTOF): Calcd. for $(M + 2H)^{2+}$: 1181.04, found: 1181.05

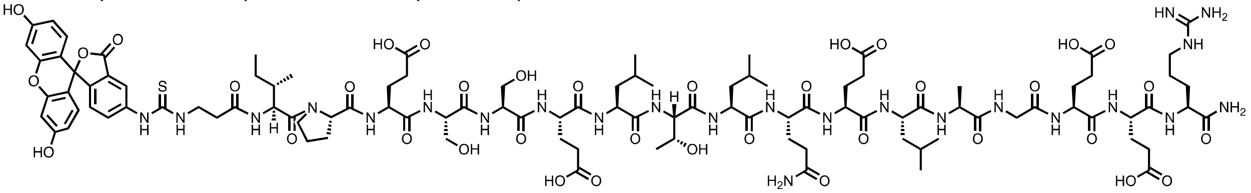

**Total ion chromatogram:**

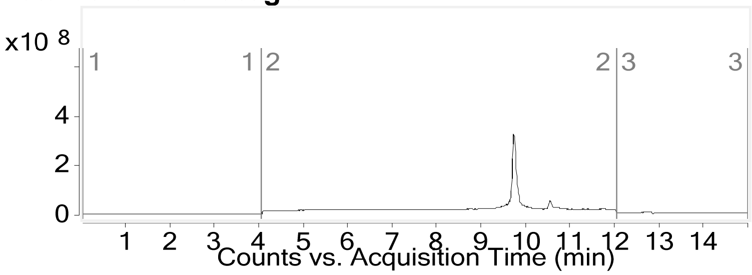


**Mass spectrum of major compound:**

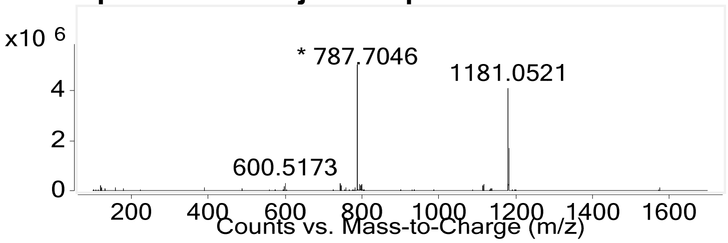


Name: peptide **A14**

HPLC: method B
HRMS (ESI-QTOF): Calcd. for $(M + 2H)^{2+}$: 1209.09, found: 1209.10.

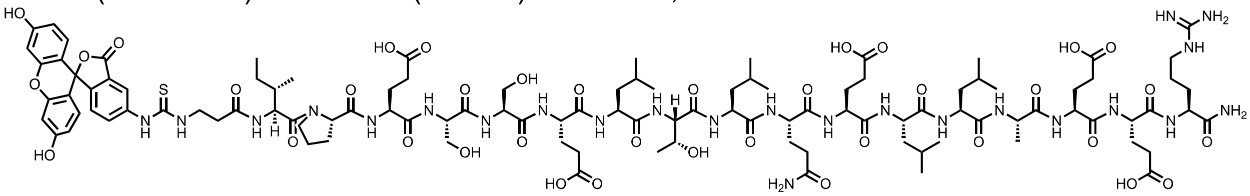

**Total ion chromatogram:**

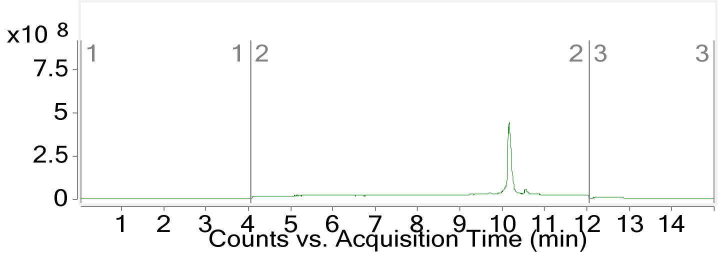


**Mass spectrum of major compound:**

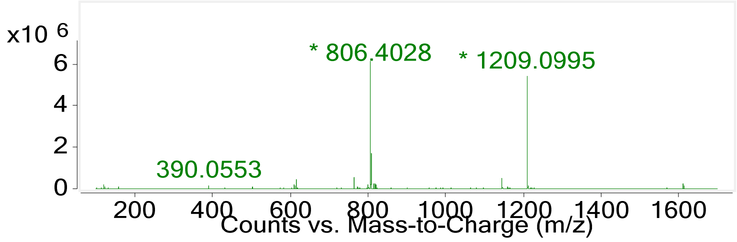


Name: peptide **A15**
HPLC: method B
HRMS (ESI-QTOF): Calcd. for $(M + 2H)^{2+}$: 1173.07, found: 1173.07

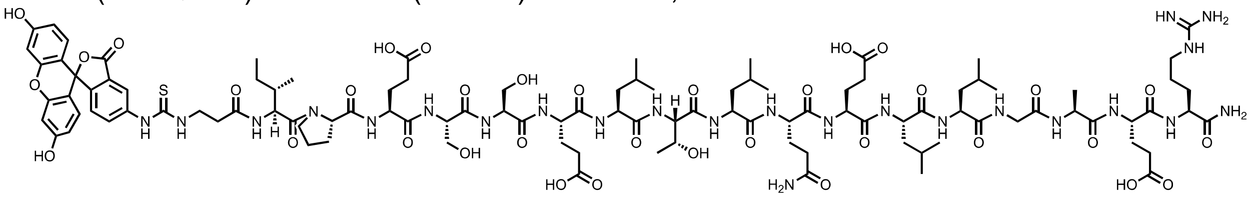

**Total ion chromatogram:**

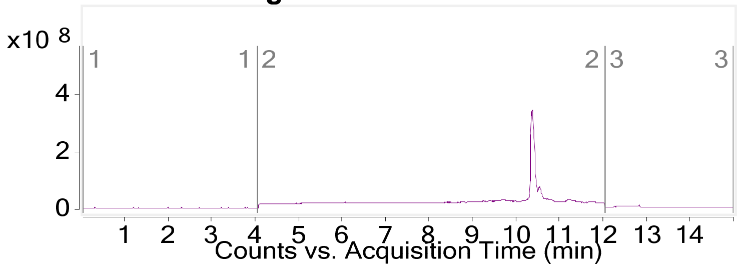


**Mass spectrum of major compound:**

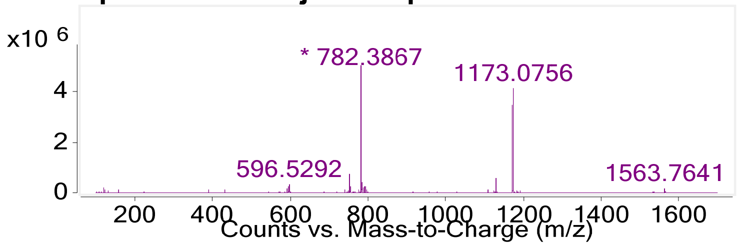

Name: peptide **A16**
HPLC: method B
HRMS (ESI-QTOF): Calcd. for $(M + 2H)^{2+}$: 1173.17, found: 1173.17.

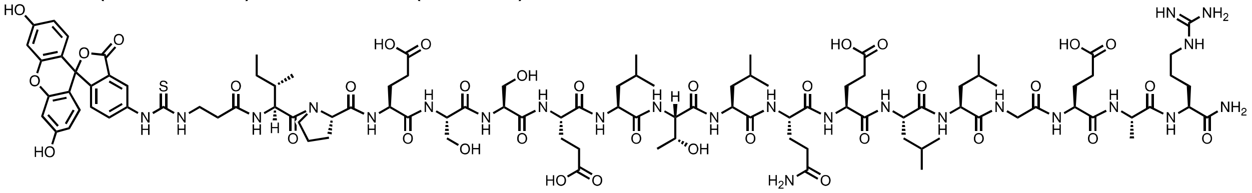

**Total ion chromatogram:**

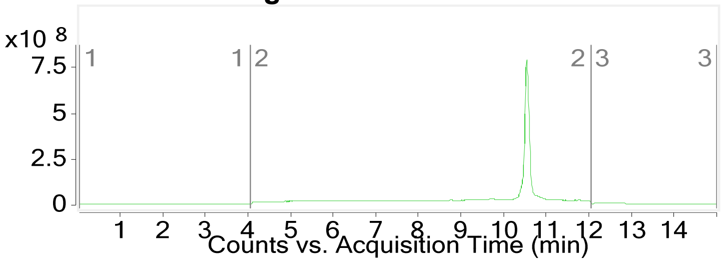


**Mass spectrum of major compound:**

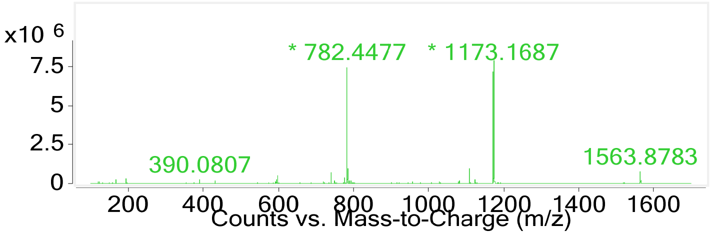


Name: peptide **A17**
HPLC: method B
HRMS (ESI-QTOF): Calcd. for $(M + 2H)^{2+}$: 1159.52, found: 1159.52

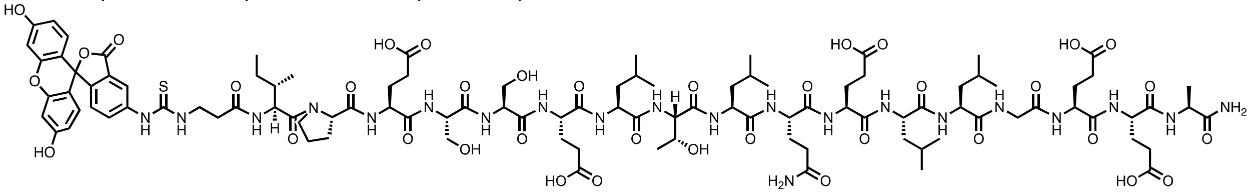

**Total ion chromatogram:**

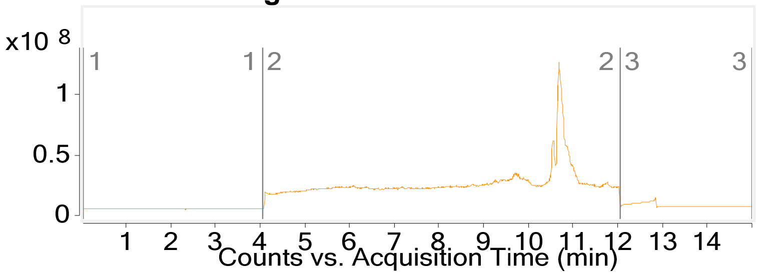


**Mass spectrum of major compound:**

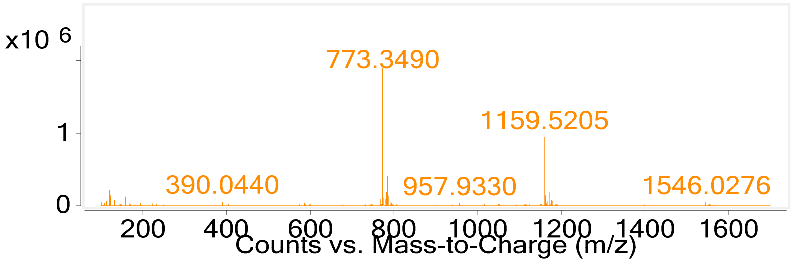


### 2.6 Peptide E1-E11

**Peptide E1**
**Sequence:** AcELTLQELLC(Phacr)EER -*CONH$_2$*

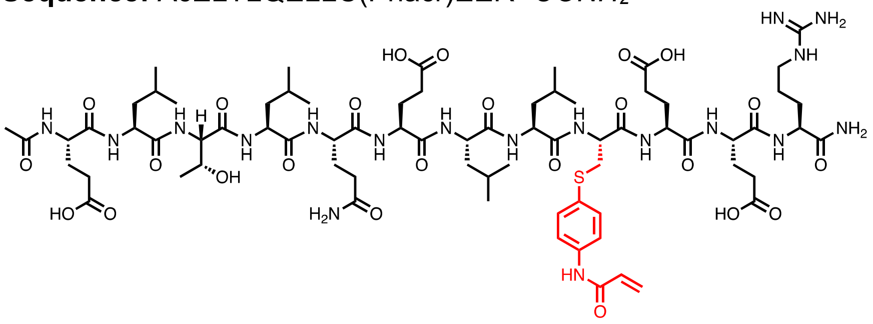


**HPLC:** Method C
**Total ion chromatogram:**

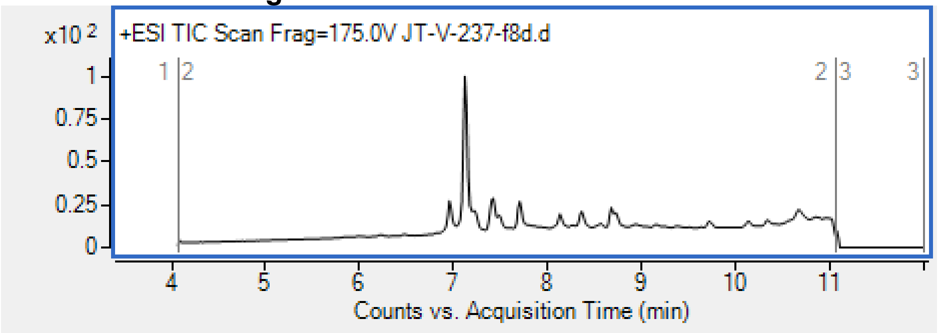


**Mass spectrum of major compound:**

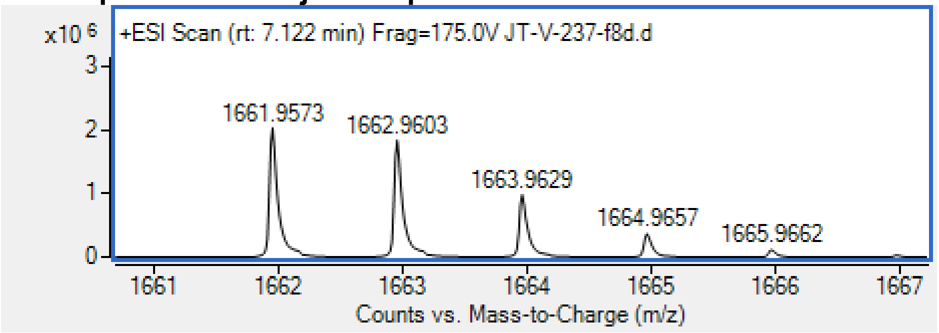


| | | |
|---|---|---|
| Exact Mass: | Calculated for $C_{73}H_{117}N_{18}O_{24}S$ $[M+H]^+$ | 1661.82 Da |
| | found | 1661.96 Da |

**Peptide E2**
**Sequence:** AcELTLQELLGC(Phacr)ER -*CONH2*

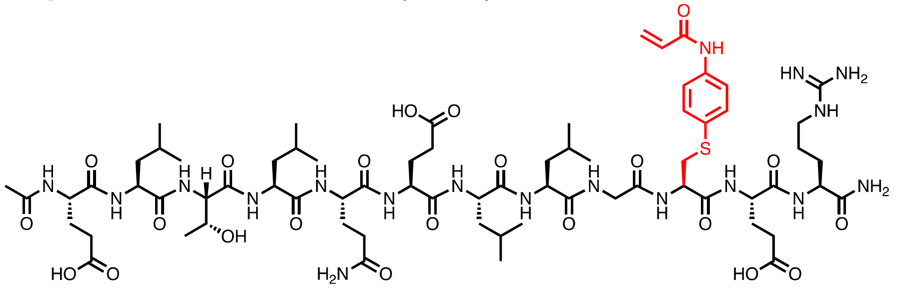

**HPLC:** Method C
**Total ion chromatogram:**

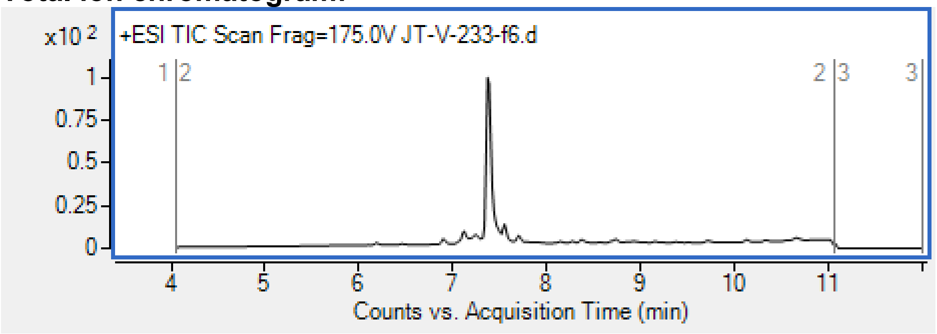


**Mass spectrum of major compound:**

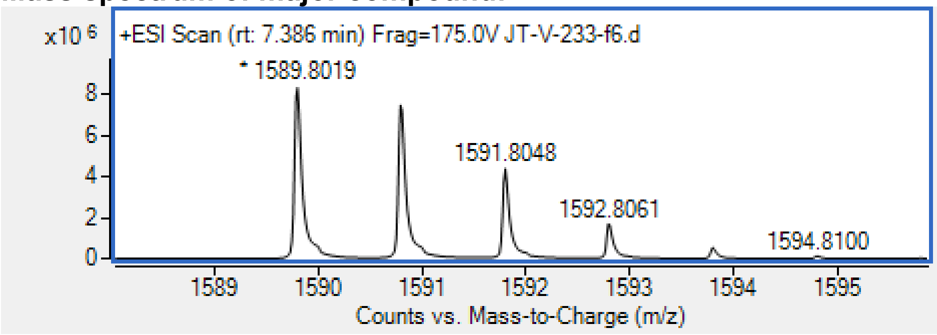


| | | | | |
|---|---|---|---|---|
| Exact Mass: | Calculated $[M+H]^+$ | for | $C_{70}H_{113}N_{18}O_{22}S$ | 1589.80 Da |
| | found | | | 1589.80 Da |

**Peptide E3**
**Sequence:** AcELTLQELL(dha)EER-*CONH2*

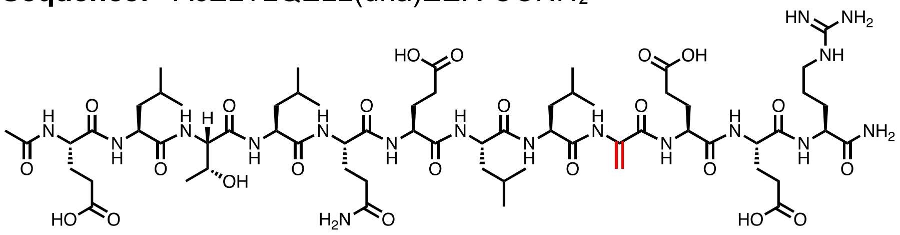

**HPLC:** Method C
**Total ion chromatogram:**

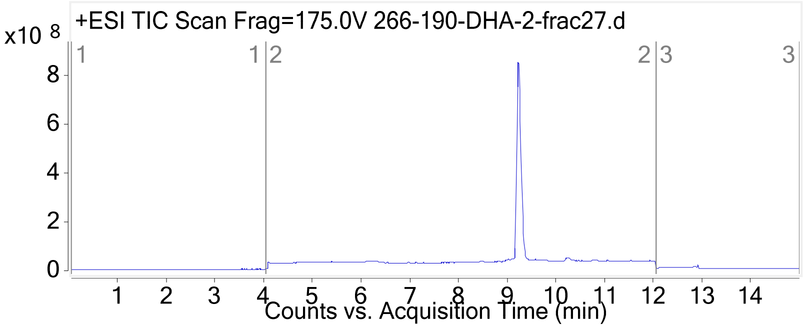


**Mass spectrum of major compound:**

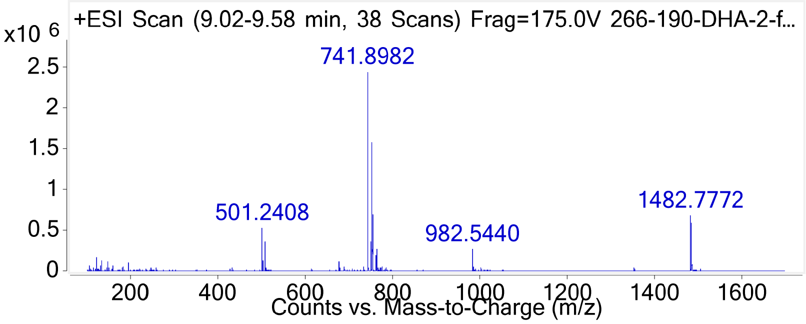


**HRMS (ESI-QTOF):** Calcd. $C_{64}H_{107}N_{17}O_{23}$ for $(M + H)^+$: 1482.77, found: 1482.77.

**Peptide E4**

**Sequence:** AcELTLQELLG(dha)ER -*CONH₂*

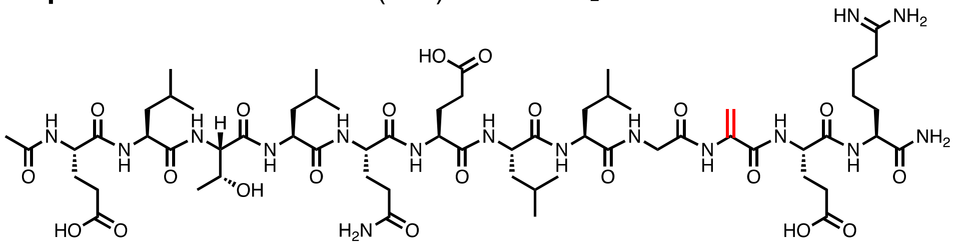

**HPLC:** Method C

**Total ion chromatogram:**

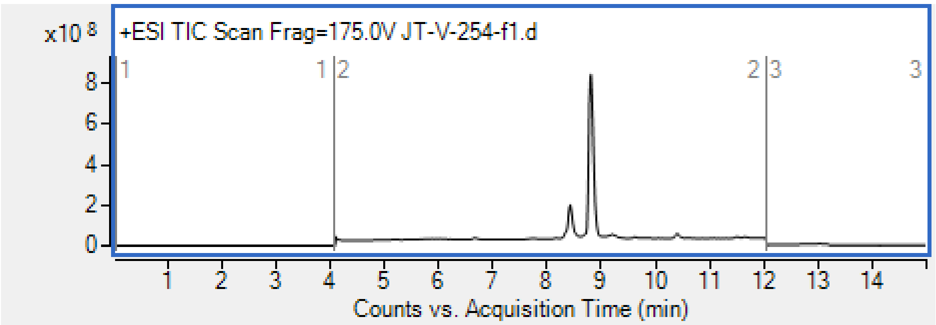


**Mass spectrum of major compound:**

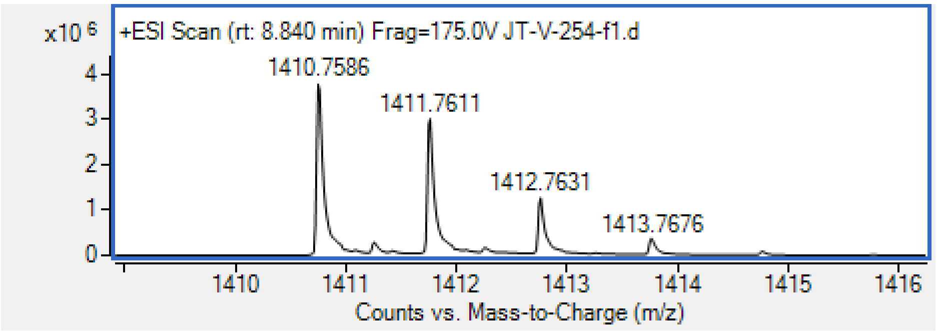


| | | |
|---|---|---|
| Exact Mass: | Calculated for $C_{61}H_{104}N_{17}O_{21}$ [M+H]$^+$ | 1410.76 Da |
| | found | 1410.76 Da |

**Peptide E5**
**Sequence:** AcELTLQELLGE(dha)R-*CONH₂*

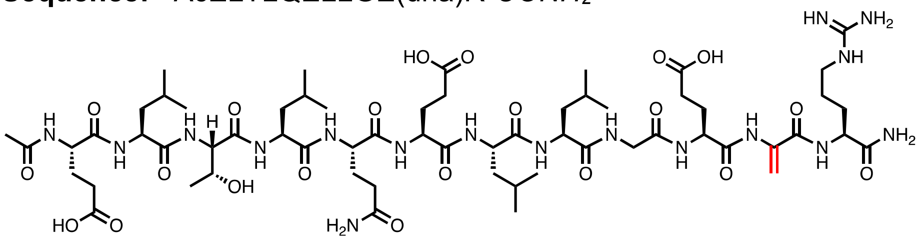

**HPLC:** Method C
**Total ion chromatogram:**

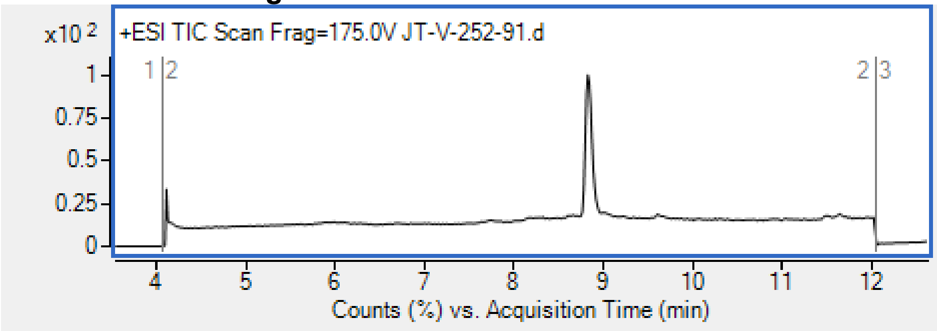


**Mass spectrum of major compound:**

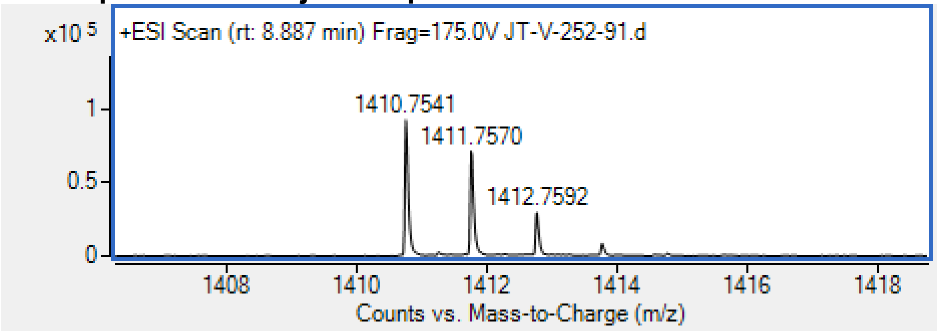


| | | |
|---|---|---|
| Exact Mass: | Calculated for $C_{62}H_{105}N_{16}O_{21}$ $[M+H]^+$ | 1410.75 Da |
| | found | 1410.75 Da |

**Peptide E6**
**Sequence:** AcELTLQELL(dap-acr)EER -*CONH₂*

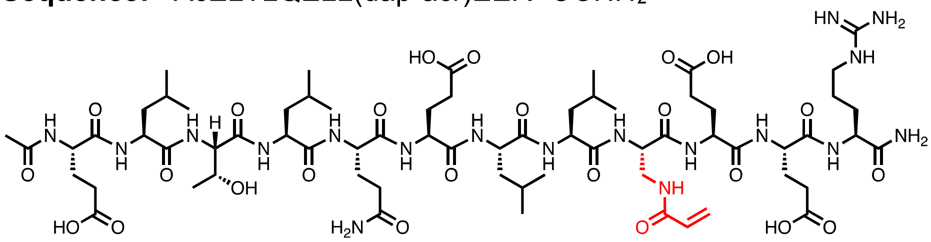

**HPLC:** Method C
**Total ion chromatogram:**

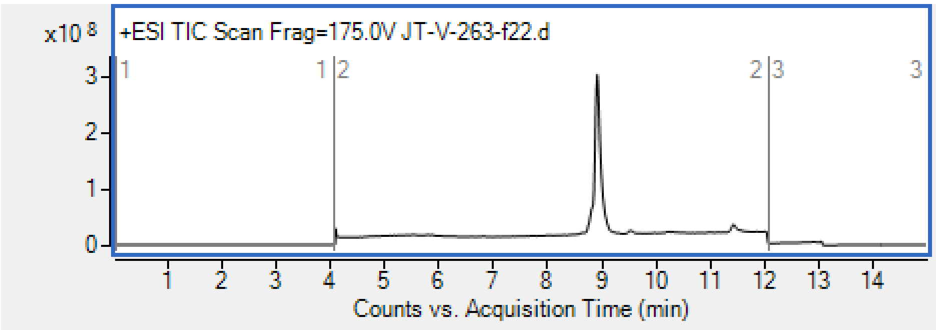


**Mass spectrum of major compound:**

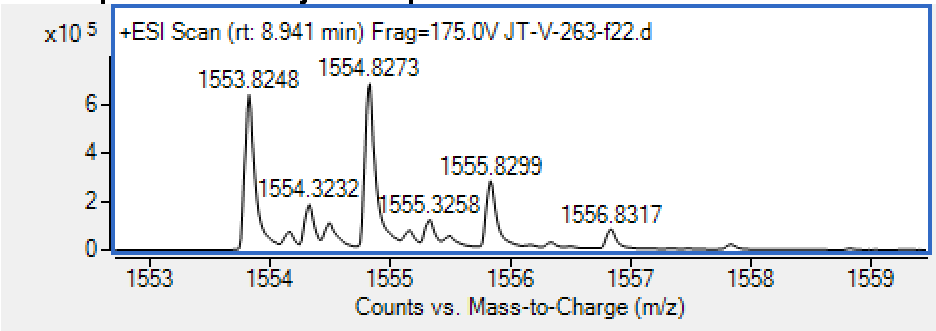


| | | |
|---|---|---|
| Exact Mass: | Calculated for $C_{67}H_{113}N_{18}O_{24}[M+H]^+$ | 1553.82 Da |
| | found | 1553.82 Da |

**Peptide E7**

**Sequence:** AcELTLQELLG(dap-acr)ER-*CONH₂*

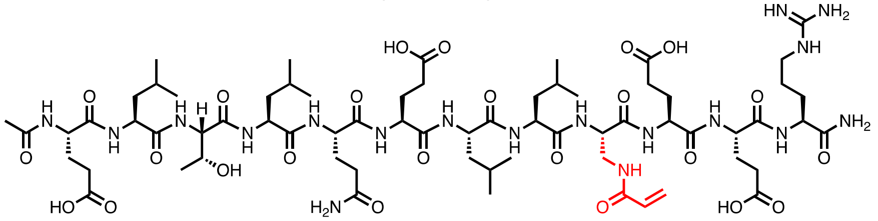

**HPLC:** Method D

**Total ion chromatogram:**

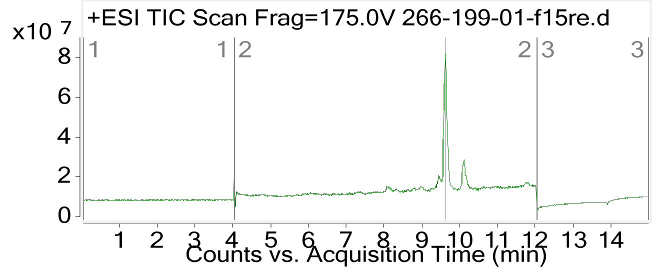


**Mass spectrum of major compound:**

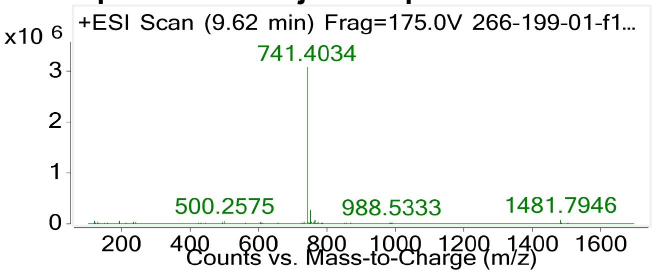

**HRMS (ESI-QTOF):** Calcd. $C_{64}H_{108}N_{18}O_{22}$ for $(M + H)^+$: 1481.79, found: 1481.79.

**Peptide E8**
**Sequence:** AcELTLQELLGE(dap-acr)R -*CONH₂*

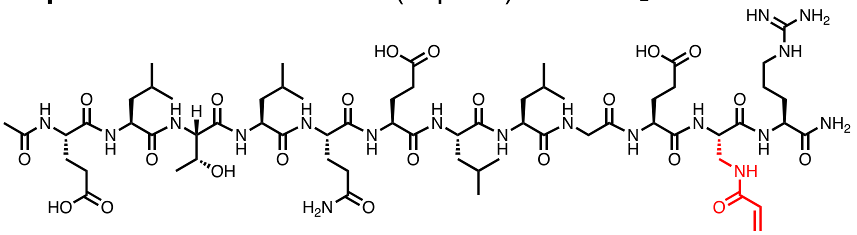

**HPLC:** Method D
**Total ion chromatogram:**

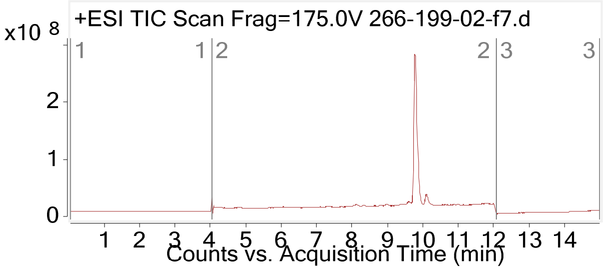


**Mass spectrum of major compound:**

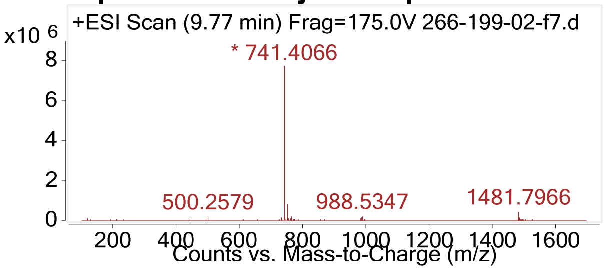


**HRMS (ESI-QTOF):** Calcd. $C_{64}H_{108}N_{18}O_{22}$ for $(M + H)^+$: 1481.79, found: 1481.79.

**Peptide E9**
**Sequence:** AcELTLQELLG(dab-acr)ER-*CONH₂*

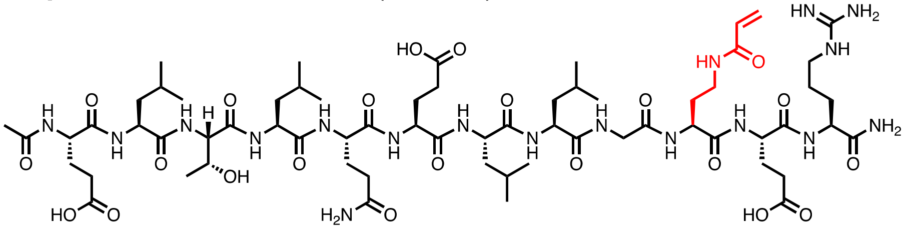

**HPLC:** Method D
**Total ion chromatogram:**

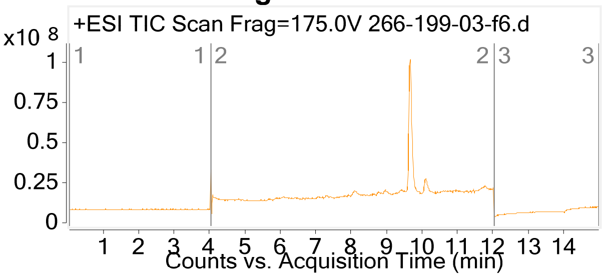

**Mass spectrum of major compound:**

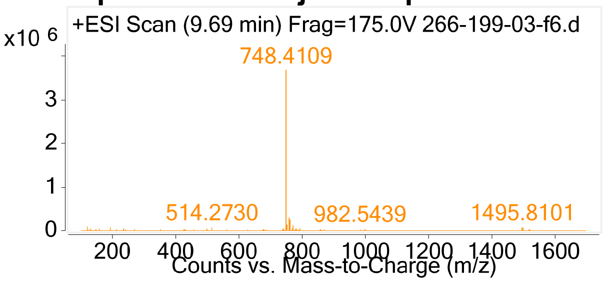


**HRMS (ESI-QTOF):** Calcd. $C_{65}H_{110}N_{18}O_{22}$ for $(M + H)^+$: 1495.81, found: 1495.81.

**Peptide E10**

**Sequence:** AcELTLQELLGE(dab-acr)R -*CONH₂*

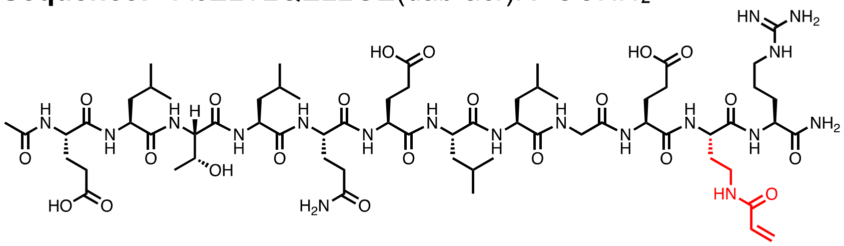

**HPLC:** Method D

**Total ion chromatogram:**

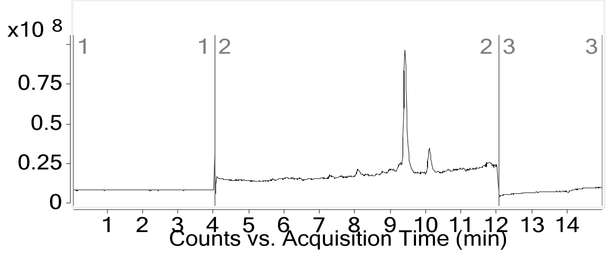


**Mass spectrum of major compound:**

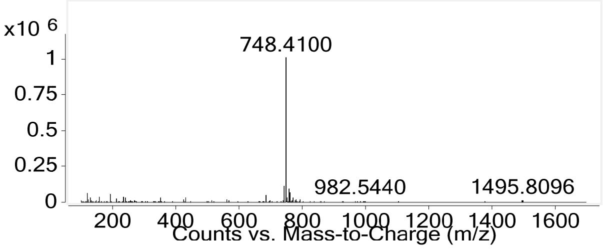


**HRMS (ESI-QTOF):** Calcd. $C_{65}H_{110}N_{18}O_{22}$ for $(M + H)^+$: 1495.81, found: 1495.81.

**Peptide E11**

**Sequence:** AcELTLQELLGE(dap-ppa)R-*CONH₂*

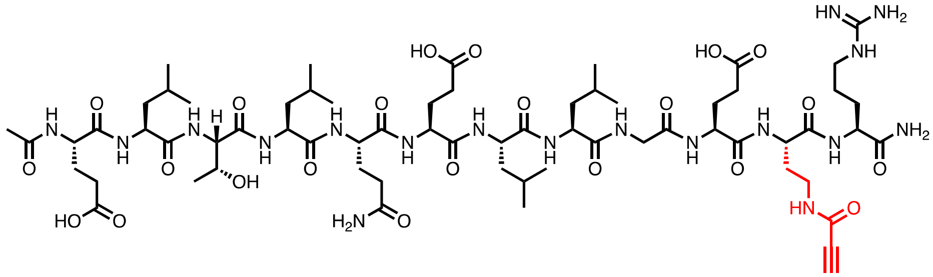

**HPLC:** Method C

**Total ion chromatogram:**

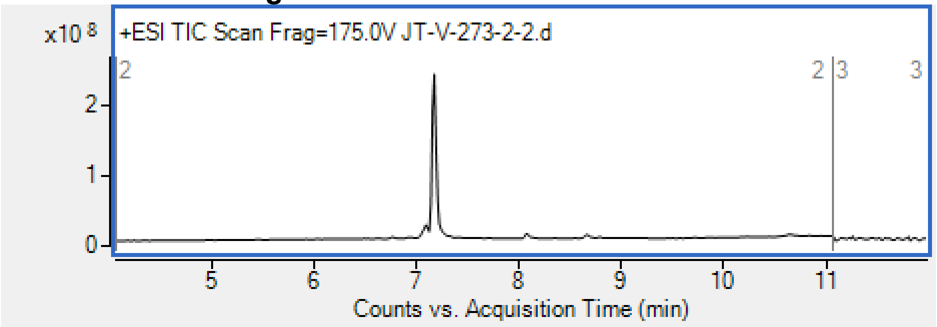


**Mass spectrum of major compound:**

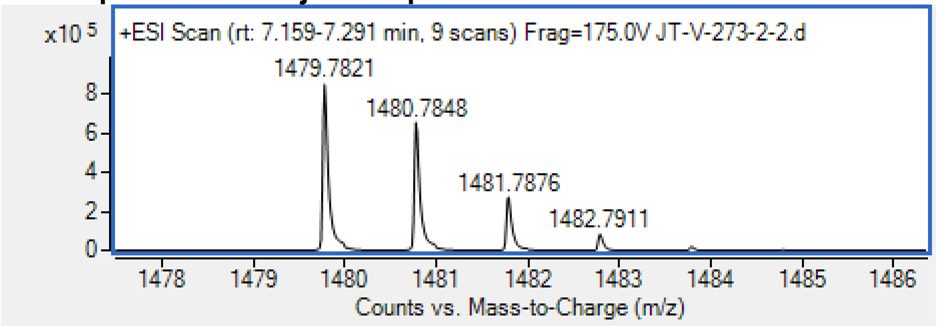


| | | |
|---|---|---|
| Exact Mass: | Calculated $C_{64}H_{107}N_{18}O_{22}$ $[M+H]^+$ | 1479.78 Da |
| | found | 1479.78 Da |

### 3. Binding characterization

Peptide **N1** binding validation. Approximately 100 nM MBP-16E6 and peptide were mixed following the protocol for **Competitive binding assay by BLI** (N=2). A representative binding curve was shown. Free [MBP-16E6] was estimated using the biotin-IPESS calibration curve, and the obtained titration curve was fitted.

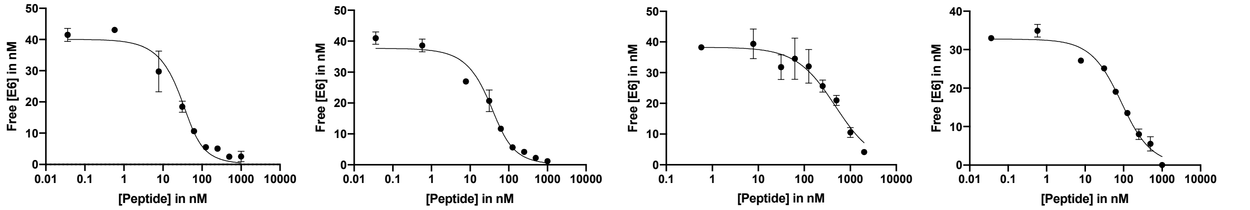


| N1 | N2 | N3 | N4 |
|---|---|---|---|
| $K_D$= 10±6.8 nM | $K_D$= 13±6.3 nM | $K_D$= 425±78 nM | $K_D$= 79±14 nM |

Peptide **N1, N2, N3 and N4** binding validation. Approximately 100 nM MBP-16E6 and peptide (1000 nM, 500 nM, 250 nM, 125nM, 63 nM, 32 nM, 7.8 nM, 0.6 nM, and 0.03 nM) were mixed following the protocol for **Competitive binding assay by BLI** (N=2). A representative binding curve was shown. Free [MBP-16E6] was estimated using the biotin-IPESS calibration curve, and the obtained titration curve was fitted.

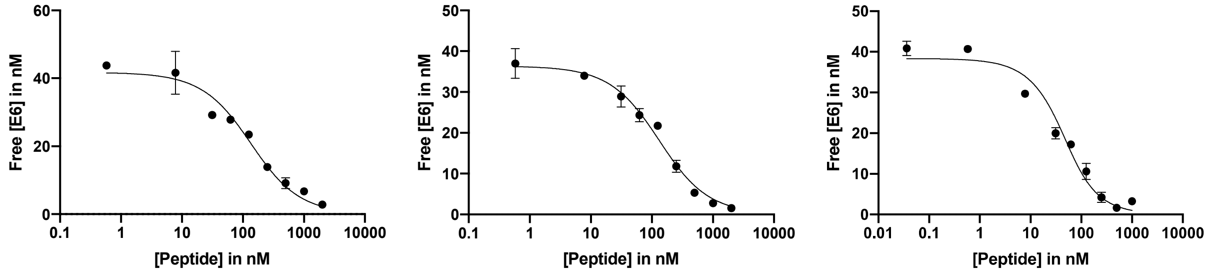


| N5 | N6 | N7 |
|---|---|---|
| $K_D$= 110±21 nM | $K_D$= 102±18 nM | $K_D$= 27±6.3 nM |

Peptide **N5, N6, and N7** binding validation. Approximately 40 nM MBP-16E6 and peptide (1000 nM, 500 nM, 250 nM, 125nM, 63 nM, 32 nM, 7.8 nM, 0.6 nM, and 0.03 nM) were mixed following the protocol for **Competitive binding assay by BLI** (N=2). A representative binding curve was shown. Free [MBP-16E6] was estimated using the biotin-IPESS calibration curve, and the obtained titration curve was fitted.

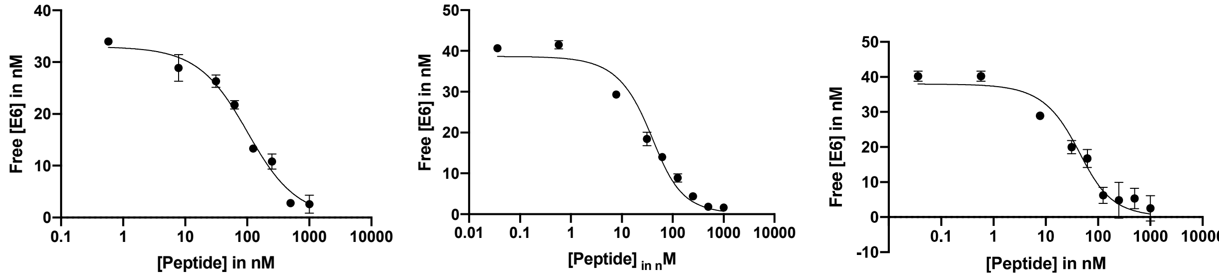


| N8 | N9 | N10 |
|---|---|---|
| $K_D$= 75±19 nM | $K_D$= 22±7.2 nM | $K_D$= 23±9.0 nM |

Peptide **N8, N9, and N1** binding validation. Approximately 40 nM MBP-16E6 and peptide (1000 nM, 500 nM, 250 nM, 125nM, 63 nM, 32 nM, 7.8 nM, 0.6 nM, and 0.03 nM) were mixed together following the protocol for **Competitive binding assay by BLI** (N=2). A representative binding curve was shown. Free [MBP-16E6] was estimated using the biotin-IPESS calibration curve, and the obtained titration curve was fitted.

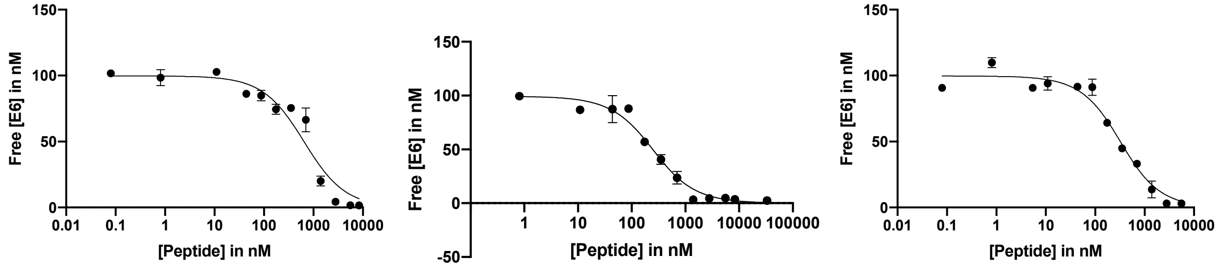


| N11 | N12 | N13 |
|---|---|---|
| $K_D$= 595±152 nM | $K_D$= 199±37nM | $K_D$= 274±51nM |

Peptide **N10, N11, and N12** binding validation. Approximately 100 nM MBP-16E6 and peptide (33333 nM, 16667 nM, 8333 nM, 4166 nM, 2083 nM, 1041 nM, 520 nM, 260 nM, 65 nM, 32nM, 4.8 nM 0.5 nM were mixed together following the protocol for **Competitive binding assay by BLI** (N=2). A representative binding curve was shown. Free [MBP-16E6] was estimated using the biotin-IPESS calibration curve, and the obtained titration curve was fitted.

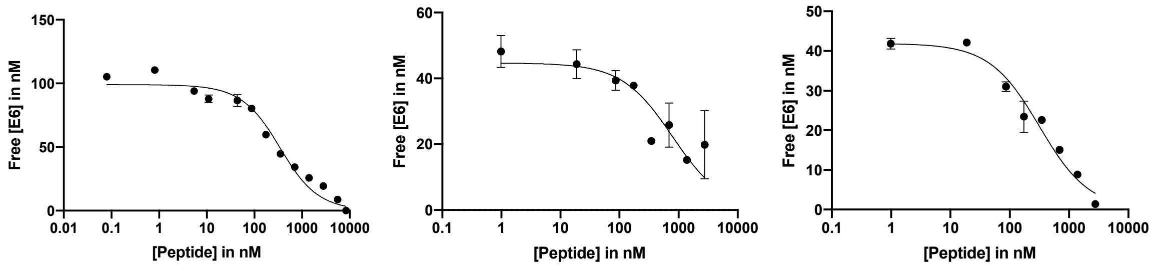


| | | |
|---|---|---|
| N14 | N15 | N16 |
| $K_D$= 292±54 nM | $K_D$= 767±295 nM | $K_D$= 228±74 nM |

Peptide **N14, N15, and N16** binding validation. Approximately 100 nM or 40nM MBP-16E6 and peptide **N14** (33333 nM, 16667 nM, 8333 nM, 4166 nM, 2083 nM, 1041 nM, 520 nM, 260 nM, 65 nM, 32nM, 4.8 nM, 0.5 nM) or **N15, N16** (2777 nM, 1388 nM, 694 nM, 347 nM, 173 nM, 86 nM, 18 nM, 1nM) were mixed together following the protocol for **Competitive binding assay by BLI** (N=2). A representative binding curve was shown. Free [MBP-16E6] was estimated using the biotin-IPESS calibration curve, and the obtained titration curve was fitted.

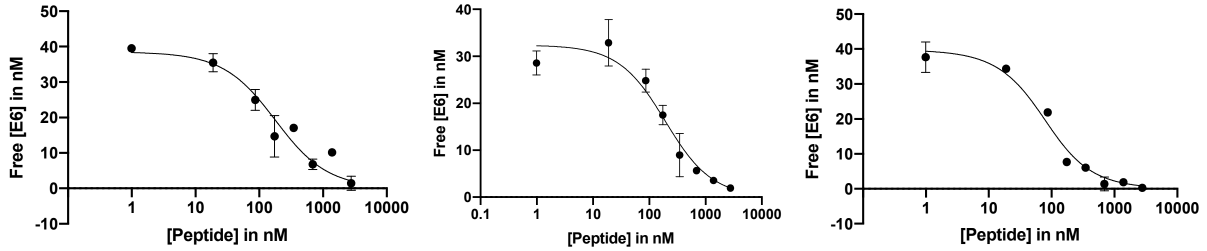


| | | |
|---|---|---|
| N17 | N18 | N19 |
| $K_D$= 159±47 nM | $K_D$= 106±27 nM | $K_D$= 55±12 nM |

Peptide **N16, N17, and N18** binding validation. Approximately 40 nM MBP-16E6 and peptide (2777 nM, 1388 nM, 694 nM, 347 nM, 173 nM, 86 nM, 18 nM, 1nM) were mixed together following the protocol for **Competitive binding assay by BLI** (N=2). A representative binding curve was shown. Free [MBP-16E6] was estimated using the biotin-IPESS calibration curve, and the obtained titration curve was fitted.

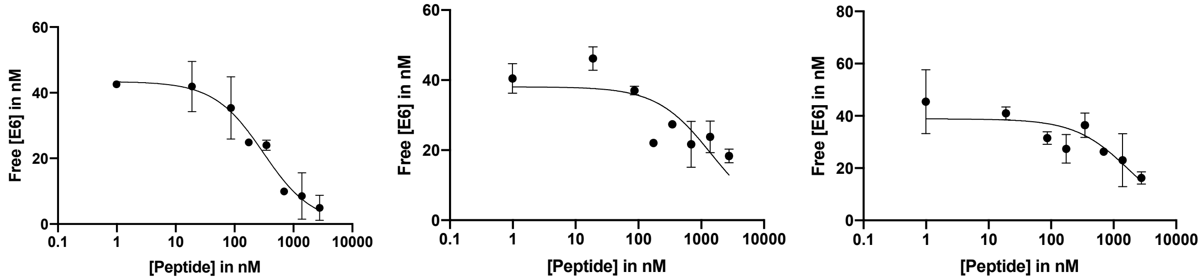


| | | |
|---|---|---|
| N20 | N21 | N22 |
| $K_D$= 213±49 nM | $K_D$= 1412±822 nM | $K_D$= 1658±705 nM |

Peptide **N19, N20, and N21** binding validation. Approximately 40 nM MBP-16E6 and peptide (2777 nM, 1388 nM, 694 nM, 347 nM, 173 nM, 86 nM, 18 nM, 1nM) were mixed together following the protocol for **Competitive binding assay by BLI** (N=2). A representative binding curve was shown. Free [MBP-16E6] was estimated using the biotin-IPESS calibration curve, and the obtained titration curve was fitted.

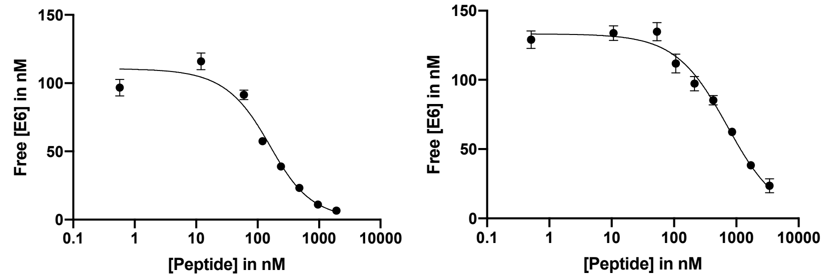


| N-FITC | C-FITC |
|---|---|
| $K_D$= 95±17 nM | $K_D$= 640±82 nM |

Peptide **N-FITC** and **C-FITC** binding validation. Approximately 100 nM or 120nM MBP-16E6 and peptide (1915 nM, 957nM, 478nM, 239nM, 119nM, 59nM, 12nM, and 0.5nM) were mixed together following the protocol for **Competitive binding assay by BLI** (N=2). A representative binding curve was shown. Free [MBP-16E6] was estimated using the biotin-IPESS calibration curve, and the obtained titration curve was fitted.

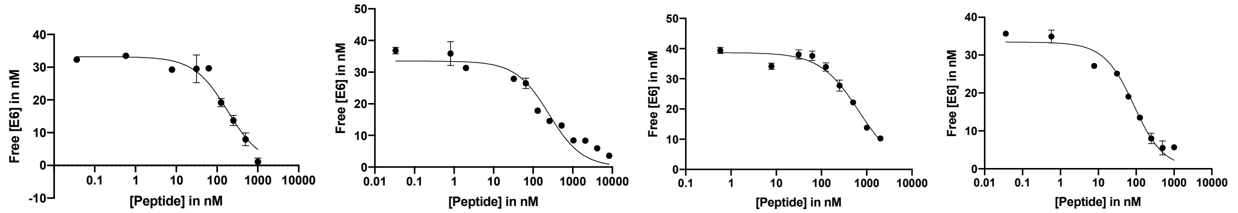


| C1 | C2 | C3 | C4 |
|---|---|---|---|
| $K_D$=165±35 nM | $K_D$=245±59 nM | $K_D$=645±92 nM | $K_D$= 71±14 nM |

Peptide **C1, C2, C3, and C4** binding validation. Approximately 40 nM MBP-16E6 and peptide (1000 nM, 500 nM, 250 nM, 125nM, 63 nM, 32 nM, 7.8 nM, 0.6 nM, and 0.06 nM) were mixed together following the protocol for **Competitive binding assay by BLI** (N=2). A representative binding curve was shown. Free [MBP-16E6] was estimated using the biotin-IPESS calibration curve, and the obtained titration curve was fitted.

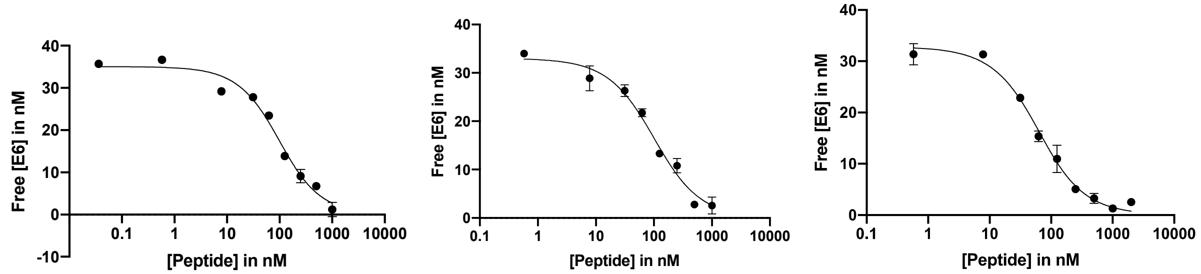


| C5 | C6 | C7 |
|---|---|---|
| $K_D$= 81±13 nM | $K_D$=84±13 nM | $K_D$= 46±5.5 nM |

Peptide **C5, C6, and C7** binding validation. Approximately 30 nM MBP-16E6 and peptide (1000 nM, 500 nM, 250 nM, 125nM, 63 nM, 32 nM, 7.8 nM, 0.6 nM) were mixed together following the protocol for **Competitive binding assay by BLI** (N=2). A representative binding curve was shown. Free [MBP-16E6] was estimated using the biotin-IPESS calibration curve, and the obtained titration curve was fitted.

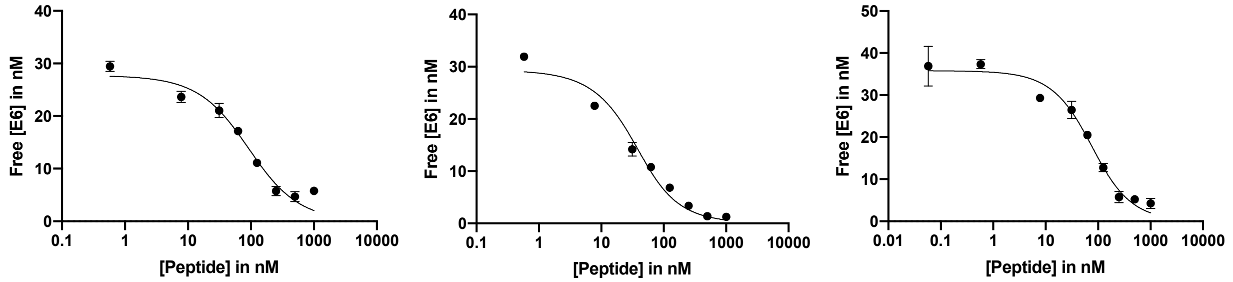


| C8 | C9 | C10 |
|---|---|---|
| $K_D$= 78±18 nM | $K_D$= 22±5.8 nM | $K_D$= 64±10 nM |

Peptide **C8, C9, and C10** binding validation. Approximately 30 or 15 nM MBP-16E6 and peptide (1000 nM, 500 nM, 250 nM, 125nM, 63 nM, 32 nM, 7.8 nM, 0.6 nM) were mixed together following the protocol for **Competitive binding assay by BLI** (N=2). A representative binding curve was shown. Free [MBP-16E6] was estimated using the biotin-IPESS calibration curve, and the obtained titration curve was fitted.

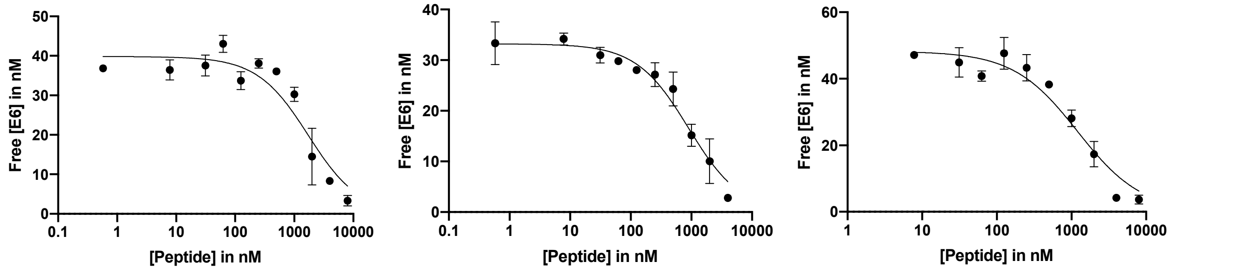


| C11 | C12 | C13 |
|---|---|---|
| $K_D$= 1459±474 nM | $K_D$= 878±125 nM | $K_D$= 1184±264 nM |

Peptide **C11, C12, and C13** binding validation. Approximately 40 nM MBP-16E6 and peptide (8000 nM, 4000 nM, 2000 nM, 1000 nM, 500 nM, 250 nM, 125nM, 63 nM, 32 nM, 7.8 nM, 0.6 nM, and 0.03 nM) were mixed together following the protocol for **Competitive binding assay by BLI** (N=2). A representative binding curve is shown. Free [MBP-16E6] was estimated using the biotin-IPESS calibration curve, and the titration curve was fitted.

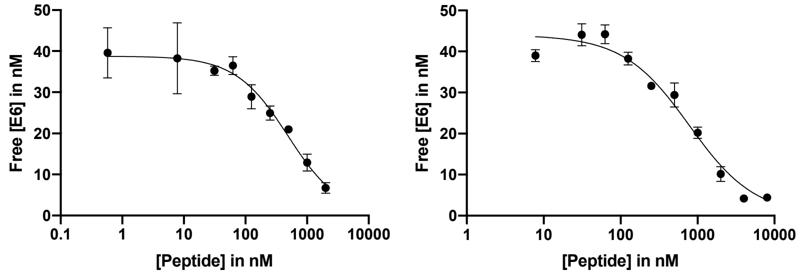


| C14 | C15 |
|---|---|
| $K_D$= 507±65nM | $K_D$= 756±128 nM |

Peptide **C14** and **C15** binding validation. Approximately 40 nM MBP-16E6 and peptide (4000 nM, 2000 nM, 1000 nM, 500 nM, 250 nM, 125nM, 63 nM, 32 nM, 7.8 nM, 0.6 nM, and 0.03 nM) were mixed together following the protocol for **Competitive binding assay by BLI** (N=2). A representative binding curve is shown. Free [MBP-16E6] was estimated using the biotin-IPESS calibration curve, and the obtained titration curve was fitted.

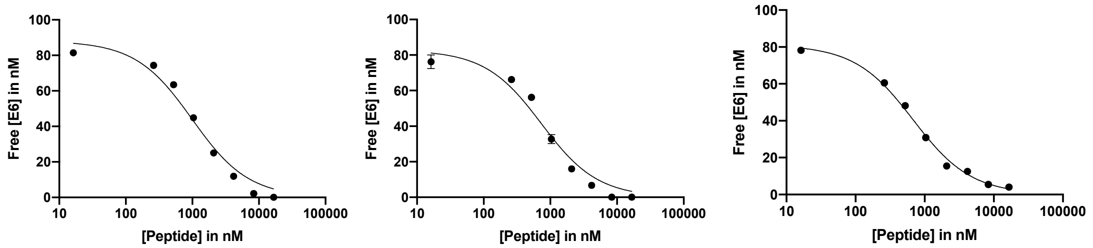


| A1 | A2 | A3 |
|---|---|---|
| FITC-APESSELTLQELLGEER | FITC-IAESSELTLQELLGEER | FITC-IPASSELTLQELLGEER |
| $K_D$= 944±182 nM | $K_D$= 691±157 nM | $K_D$= 638±54 nM |

Peptide **A1, A2, and A3** binding validation. Approximately 80 nM MBP-16E6 and peptide (16667 nM, 8333 nM, 4166 nM, 2083 nM, 1041 nM, 520 nM, 260 nM, 16 nM, and 1.6 nM) were mixed together following the protocol for **Competitive binding assay by BLI** (N=2). A representative binding curve was shown. Free [MBP-16E6] was estimated using the biotin-IPESS calibration curve, and the obtained titration curve was fitted.

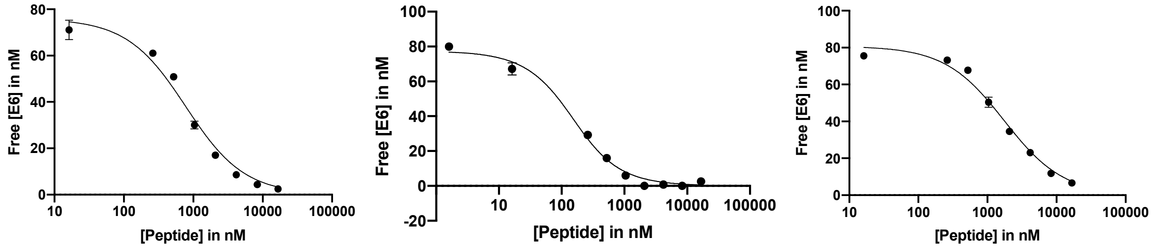


| A4 | A5 | A6 |
|---|---|---|
| FITC-IPEASELTLQELLGEER | FITC-IPESAELTLQELLGEER | FITC-IPESSALTLQELLGEER |
| $K_D$= 739±113 nM | $K_D$=125±17 nM | $K_D$= 1706±209 nM |

Peptide **A4, A5, and A6** binding validation. Approximately 80 nM MBP-16E6 and peptide (16667 nM, 8333 nM, 4166 nM, 2083 nM, 1041 nM, 520 nM, 260 nM, 16 nM, and 1.6 nM) were mixed together following the protocol for **Competitive binding assay by BLI** (N=2). A representative binding curve was shown. Free [MBP-16E6] was estimated using the biotin-IPESS calibration curve, and the obtained titration curve was fitted.

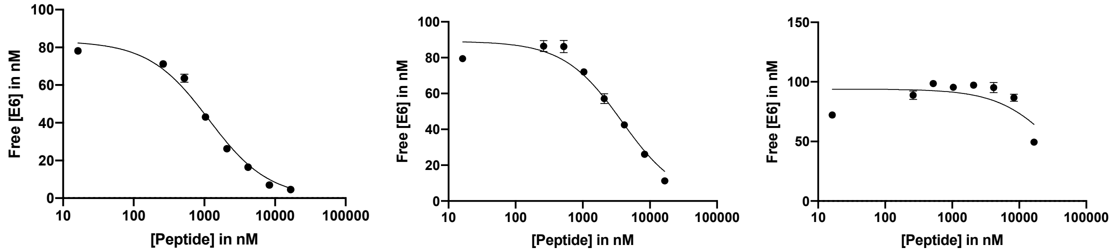


| A7 | A8 | A9 |
|---|---|---|
| FITC-IPESSEATLQELLGEER | FITC-IPESSELALQELLGEER | FITC-IPESSELTAQELLGEER |
| $K_D$= 1114±152 nM | $K_D$=3748±680 nM | $K_D$>10000 nM |

Peptide **A7, A8, and A9** binding validation. Approximately 80 nM MBP-16E6 and peptide (16667 nM, 8333 nM, 4166 nM, 2083 nM, 1041 nM, 520 nM, 260 nM, 16 nM) were mixed together following the protocol for **Competitive binding assay by BLI** (N=2). A representative binding curve was shown. Free [MBP-16E6] was estimated using the biotin-IPESS calibration curve, and the obtained titration curve was fitted.

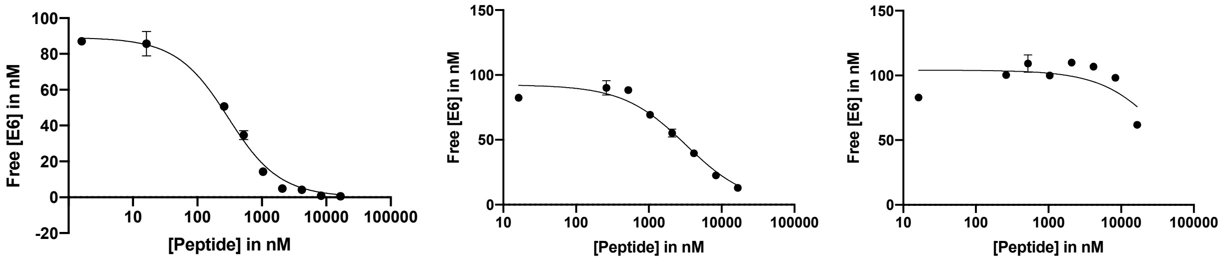


| A10 | A11 | A12 |
|---|---|---|
| FITC-IPESSELTLAELLGEER | FITC-IPESSELTLQALLGEER | FITC-IPESSELTLQEALGEER |
| $K_D$= 242±31 nM | $K_D$=3158±568 nM | $K_D$>10000 nM |

Peptide **A10, A11, and A12** binding validation. Approximately 80 nM MBP-16E6 and peptide (16667 nM, 8333 nM, 4166 nM, 2083 nM, 1041 nM, 520 nM, 260 nM, 16 nM) were mixed together following the protocol for **Competitive binding assay by BLI** (N=2). A representative binding curve was shown. Free [MBP-16E6] was estimated using the biotin-IPESS calibration curve, and the obtained titration curve was fitted.

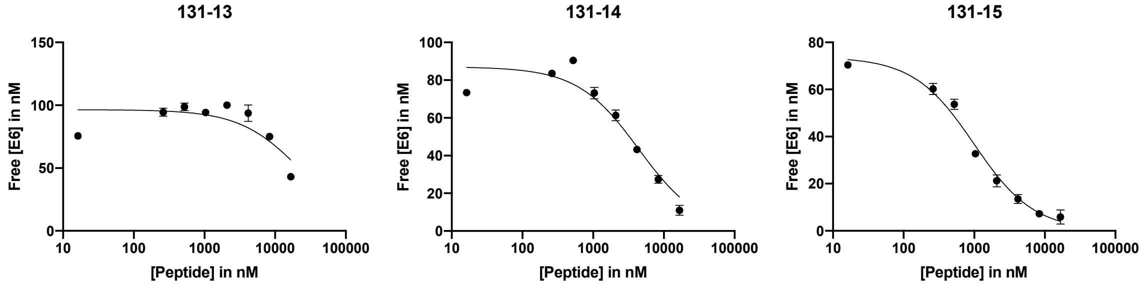


| A13 | A14 | A15 |
|---|---|---|
| FITC-IPESSELTLQELAGEER | FITC-IPESSELTLQELLAEER | FITC-IPESSELTLQELLGAER |
| $K_D$>10000 nM | $K_D$= 4342±1177 nM | $K_D$=953±119 nM |

Peptide **A13, A14, and A15** binding validation. Approximately 80 nM MBP-16E6 and peptide (16667 nM, 8333 nM, 4166 nM, 2083 nM, 1041 nM, 520 nM, 260 nM, 16 nM) were mixed together following the protocol for **Competitive binding assay by BLI** (N=2). A representative binding curve was shown. Free [MBP-16E6] was estimated using the biotin-IPESS calibration curve, and the obtained titration curve was fitted.

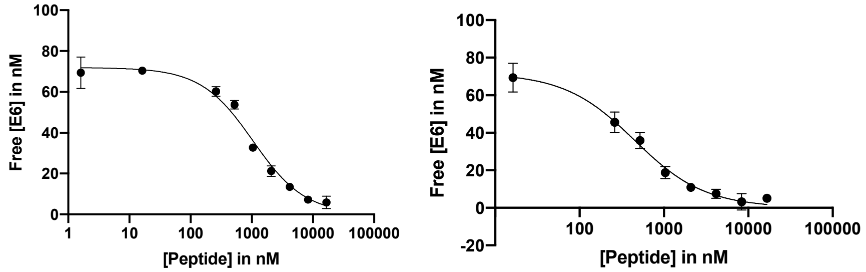


| A16 | A17 |
|---|---|
| FITC-IPESSELTLQELLGEAR | FITC-IPESSELTLQELLGEEA |
| $K_D$= 274±21 nM | $K_D$=413±39 nM |

Peptide **A16 and A17** binding validation. Approximately 70 nM MBP-16E6 and peptide (16667 nM, 8333 nM, 4166 nM, 2083 nM, 1041 nM, 520 nM, 260 nM, 16 nM) were mixed together following the protocol for **Competitive binding assay by BLI** (N=2). A representative binding curve was shown. Free [MBP-16E6] was estimated using the biotin-IPESS calibration curve, and the obtained titration curve was fitted.

Peptide 1 binding see Figure S1.

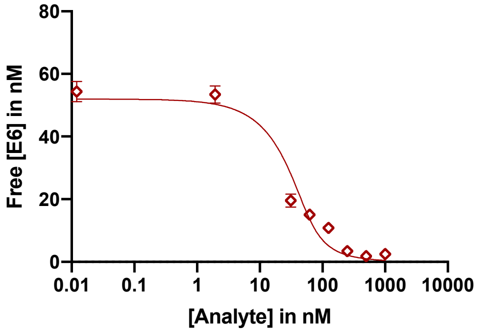


**Peptide 2** binding validation. Approximately 50 nM MBP-16E6 and peptide (1000 nM, 500 nM, 250 nM, 125 nM, 62 nM, 31 nM, 1.9 nM, 0.012 nM) were mixed together following the protocol for **Competitive binding assay by BLI** (N=3). A representative binding curve was shown. Free [MBP-16E6] was estimated using the biotin-IPESS calibration curve, and the obtained titration curve was fitted.

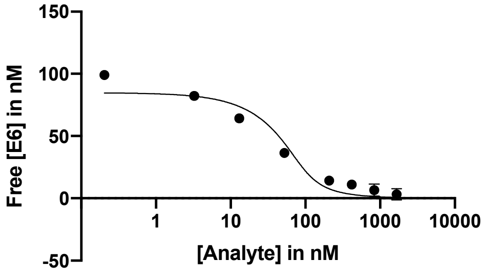


**Peptide 3** binding validation. Approximately 100 nM MBP-16E6 and peptide (1667 nM, 833 nM, 416 nM, 208 nM, 52 nM, 13 nM, 3.0 nM, and 0.2 nM) were mixed together following the protocol for **Competitive binding assay by BLI** (N=3). A representative binding curve was shown. Free [MBP-16E6] was estimated using the biotin-IPESS calibration curve, and the obtained titration curve was fitted.

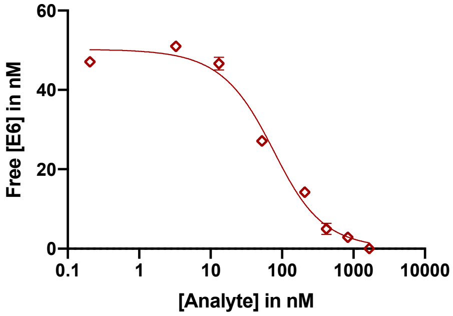


**Peptide 4** binding validation. Approximately 50 nM MBP-16E6 and peptide (1667 nM, 833 nM, 416 nM, 208 nM, 52 nM, 13 nM, 3.0 nM, and 0.2 nM) were mixed together

following the protocol for **Competitive binding assay by BLI** (N=3). A representative binding curve was shown. Free [MBP-16E6] was estimated using the biotin-IPESS calibration curve, and the obtained titration curve was fitted.

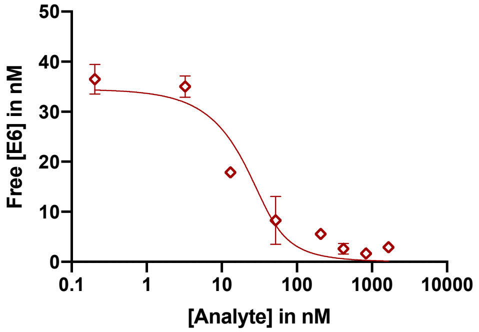


**Peptide 5** binding validation. Approximately 40 nM MBP-16E6 and peptide (1667 nM, 833 nM, 416 nM, 208 nM, 52 nM, 13 nM, 3.0 nM, and 0.2 nM) were mixed together following the protocol for **Competitive binding assay by BLI** (N=3). A representative binding curve was shown. Free [MBP-16E6] was estimated using the biotin-IPESS calibration curve, and the obtained titration curve was fitted. Peptide **5** $K_D$= 6.7±3.9 nM,

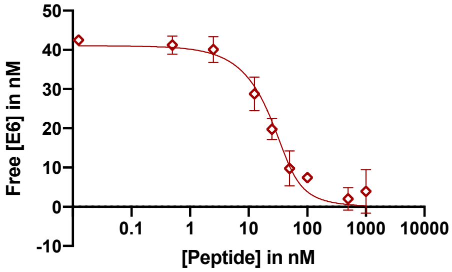


**Peptide 6** binding validation. Approximately 40 nM MBP-16E6 and peptide (1000 nM, 500 nM, 100 nM, 50 nM, 25 nM, 12 nM, 2.5 nM, 0.5 nM, and 0.0125 nM)were mixed together following the protocol for **Competitive binding assay by BLI** (N=2). A representative binding curve was shown. Free [MBP-16E6] was estimated using the biotin-IPESS calibration curve, and the obtained titration curve was fitted. Peptide **6** $K_D$= 3.7±1.9 nM,

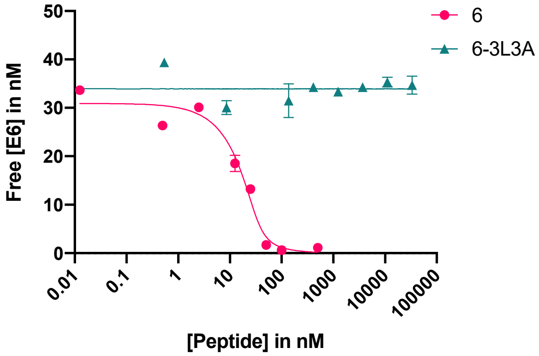


**Peptide 6' ** and **6'-3L3A**. Binding validation. Approximately 40 nM MBP-16E6 and peptide **6** (500 nM, 100 nM, 50 nM, 25 nM, 12 nM, 2.5 nM, 0.5 nM, and 0.0125 nM) were mixed

together following the protocol for **Competitive binding assay by BLI** (N=3). With peptide **6-3L3A** 33333 nM, 11111 nM, 3703 nM, 1234 nM, 411 nM, 137 nM, 8.5 nM, and 0.5 nM (N=2). Free [MBP-16E6] was estimated using the biotin-IPESS calibration curve, and the obtained titration curve was fitted. Peptide **6** $K_D$= 2.7±1.2 nM, Peptide **6-3L3A** $K_D$ > 33,333 nM

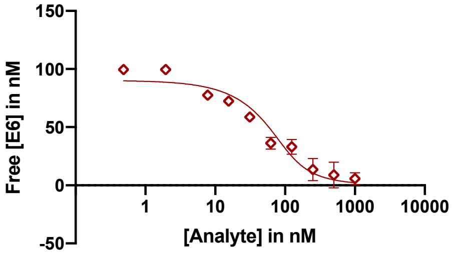


**Peptide 7** binding validation. Approximately 100 nM MBP-16E6 and peptide (1000 nM, 500 nM, 250 nM, 125 nM, 62 nM, 31 nM, 15 nM, 7.8 nM, 3.9 nM, 1.9 nM, and 0.5 nM) were mixed together following the protocol for **Competitive binding assay by BLI** (N=2). A representative binding curve was shown. Free [MBP-16E6] was estimated using the biotin-IPESS calibration curve, and the obtained titration curve was fitted. Estimated apparent $K_i$ = 23±8.1nM.

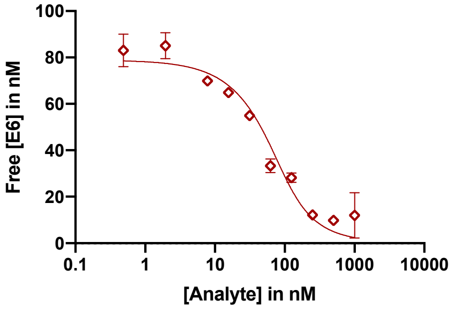


**Peptide 8** binding validation. Approximately 80 nM MBP-16E6 and peptide (1000 nM, 500 nM, 250 nM, 125 nM, 62 nM, 31 nM, 15 nM, 7.8 nM, 3.9 nM, 1.9 nM, and 0.5 nM) were mixed together following the protocol for **Competitive binding assay by BLI** (N=3). A representative binding curve was shown. Free [MBP-16E6] was estimated using the biotin-IPESS calibration curve, and the obtained titration curve was fitted. Estimated apparent $K_i$ = 39±6.1nM.

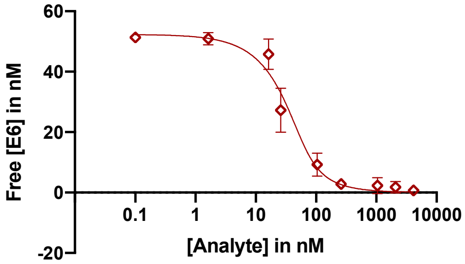


**Peptide 9** binding validation. Approximately 80 nM MBP-16E6 and peptide (4167 nM, 2084 nM, 1042 nM, 260 nM, 104 nM, 26 nM, 16 nM, 1.6 nM, 0.1 nM) were mixed together following the protocol for **Competitive binding assay by BLI** (N=3). A representative binding curve was shown. Free [MBP-16E6] was estimated using the biotin-IPESS

calibration curve, and the obtained titration curve was fitted. Estimated apparent $K_i$ = 11±3.7 nM.

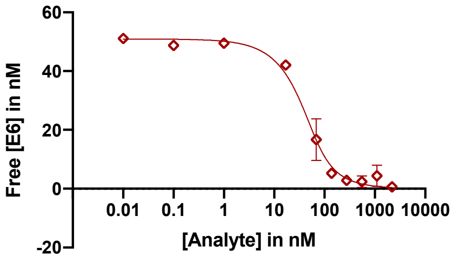


**Peptide 10** binding validation. Approximately 80 nM MBP-16E6 and peptide (4167 nM, 2084 nM, 1042 nM, 260 nM, 104 nM, 26 nM, 16 nM, 1.6 nM, 0.1 nM) were mixed together following the protocol for **Competitive binding assay by BLI** (N=3). A representative binding curve was shown. Free [MBP-16E6] was estimated using the biotin-IPESS calibration curve, and the obtained titration curve was fitted. Estimated apparent $K_i$ = 16±2.6 nM.

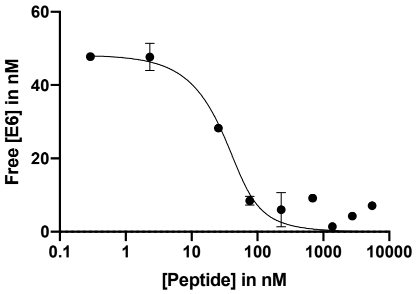


**Peptide 11** binding validation. Approximately 50 nM MBP-16E6 and peptide (5500 nM, 2750 nM, 1375 nM, 687 nM, 229 nM, 76 nM, 25 nM, 2.3 nM, and 0.2 nM) were mixed together following the protocol for **Competitive binding assay by BLI** (N=3). A representative binding curve was shown. Free [MBP-16E6] was estimated using the biotin-IPESS calibration curve, and the obtained titration curve was fitted. Estimated apparent $K_i$ 11±4.9nM.

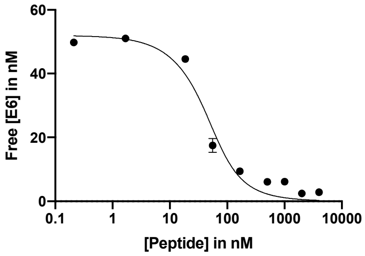


**Peptide 12** binding validation. Approximately 80 nM MBP-16E6 and peptide (4000 nM, 2000 nM, 1000 nM, 500 nM, 166 nM, 55 nM, 18 nM, 1.7 nM, and 0.2 nM) were mixed together following the protocol for in-solution competition assay (N=3). A representative binding curve was shown. Free [MBP-16E6] was estimated using the biotin-IPESS calibration curve, and the obtained titration curve was fitted. Estimated apparent $K_i$ = 20±5.4nM.

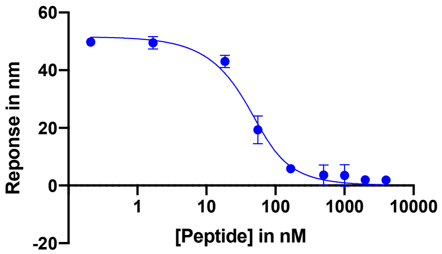


**Peptide 13** binding validation. Approximately 80 nM MBP-16E6 and peptide (4000 nM, 2000 nM, 1000 nM, 500 nM, 166 nM, 55 nM, 18 nM, 1.7 nM, and 0.2 nM) were mixed together following the protocol for in-solution competition assay (N=3). A representative binding curve was shown. Free [MBP-16E6] was estimated using the biotin-IPESS calibration curve, and the obtained titration curve was fitted. Estimated apparent $K_i$ = 17±3.9nM
